# CheckAMG: Accurate Identification of Auxiliary Viral Genes with Genome Language Models

**DOI:** 10.64898/2026.09.23.753886

**Authors:** James C. Kosmopoulos, Cody Martin, James M. Wainaina, Benjamin Bolduc, Marion Urvoy, Matthew B. Sullivan, Karthik Anantharaman

## Abstract

Viruses shape microbial metabolism through auxiliary viral genes (AVGs) that reprogram host metabolism, physiology, and gene regulation during infection. Identifying AVGs is complicated by ambiguous viral versus cellular assignment, inconsistent curation, and reliance on sequence homology that misses divergent functions. Here we present CheckAMG, which pairs annotation-based AVG prediction, curated with a per-gene viral scoring system and ontological mapping of metabolic function, with a finetuned genome-language model for annotation-independent discovery. Benchmarked against DRAM-V and VIBRANT across three ecosystems, CheckAMG produced more strongly supported auxiliary metabolic gene (AMG) calls, and uniquely among AMG tools reports auxiliary physiological (APGs) and regulatory (AReGs) genes. Applied to 1,005,980 fragmented or complete soil and human-gut viral genomes, CheckAMG identified 541,199 AVGs tracking biome-specific protein clusters and functions, including 109,107 with no detectable sequence similarity to any reference database. CheckAMG thus provides a reproducible, scalable framework for identifying the genes through which viruses reprogram host biology.

## Main

Viruses of microorganisms are fundamental drivers of global biogeochemical cycles and host microbiome dynamics, in part through their ability to reprogram host metabolism using tools such as auxiliary viral genes (AVGs)^1^. By expressing host-derived^2^ genes during infection that redirect cellular metabolic pathways (auxiliary metabolic genes, AMGs)^3,4^, alter cellular physiology (auxiliary physiological genes, APGs), or modulate host or auxiliary viral gene expression (auxiliary regulatory genes, AReGs), phages and viruses actively manipulate host cellular state^5,6^, nutrient flux^4,7–9^, and community structure^10,11^ across diverse ecosystems. Examples range from cyanophages using photosynthesis genes to maintain host photosystem activity during infection^12–14^, to viral-encoded CRISPR-Cas subunits that modulate host immunity^15–17^, and to viral-encoded sigma factors that reprogram host transcriptional networks associated with sporulation and dormancy^6,18^, to name a few. However, while metagenomic sequencing has expanded the known virosphere by orders of magnitude^19^, our ability to decode the functional mechanisms through which viruses manipulate their hosts remains disproportionately limited^20^.

Extracting these functional elements from vast metagenomic datasets presents a formidable computational challenge, with reliable AVG cataloging constrained by two compounding bottlenecks^21^. First, because AVGs are host-derived genes encoded by viruses, non-viral sequence contamination is problematic. Imprecisely predicted prophage boundaries^22^, degraded prophages^23^, nested mobile elements^24^, and/or chimeric assemblies^25^ can all contain genes that resemble the very host-like sequence expected of a genuine AVG^26^. This contamination can therefore introduce false positives^1^, because a non-viral gene cannot be a genuine AVG. Second, existing automated annotation tools, such as DRAM-V^27^ and VIBRANT^28^, apply inconsistent and often overly permissive curation criteria^1^ to their predictions. AVG predictions must be curated to ensure the sequence is genuinely viral and to ensure the encoded function is auxiliary, rather than merely involved in ultimate production of a virus particle^1^. The conventions of such curation vary by the environment being studied^21,26,29^, yielding conflicting AVG catalogs from identical datasets. These limitations bias current approaches towards genes encoded by well-characterized viruses and acting in well-characterized pathways, leaving a vast fraction of the viral functional landscape entirely hidden in complex and fragmented metagenomic datasets.

An additional factor that compounds both problems is that existing workflows for AVG detection rely almost exclusively on sequence-homology-dependent approaches against static databases^20,30^, and are therefore unable to recognize auxiliary genes whose function has diverged beyond recognizable similarity to any reference. This leaves an unknown but likely substantial share of the AVG pool as uncharacterized viral dark matter^20^. Protein language models and structure-aware, annotation-independent approaches have begun to close comparable gaps in homology detection elsewhere in microbial genomics, from general-purpose protein structure prediction^31^ to genome-context models that identify anti-phage defense systems directly from sequence^32^. However, these deep learning paradigms have not yet been adapted to the coupled challenge of simultaneously predicting viral origin and auxiliary function of genes.

Here we present CheckAMG, a bioinformatic pipeline that can identify putative AVGs (including AMGs, APGs, and AReGs) using both a traditional, reference-dependent annotation approach and an AI-based approach that is independent of sequence homology to any reference database of proteins with labeled functions. CheckAMG pairs interpretable annotation-based curation, a genome-context-based classifier of per-protein viral origin using a viral scoring system (V-scores)^33^, and an auxiliary-function weighting scheme with a protein-based genome language model finetuned from a foundational model (Protein Set Transformer) developed specifically for viromics^34^. We show that CheckAMG places a larger share of its high-confidence AMG calls on geNomad-viral sequences than DRAM-V^27^ does at its default setting, benchmarking against DRAM-V and VIBRANT^28^ across metagenomes, viromes, and viral genomes drawn from three different ecosystems (aquatic, soil, and the human gut). Applied to more than one million fragmented or complete soil and human-gut viral genomes^19^, all uncultivated, CheckAMG recovers several hundred thousand AVGs, including over 100,000 with no detectable sequence similarity to any tested reference database (but strong AVG-like signal in genome language model embedding space), revealing auxiliary functions and broad gene clusters that track the known biochemistry of each biome. CheckAMG thus provides a reproducible, scalable framework for identifying and interpreting the auxiliary genes through which viruses shape microbial metabolism across Earth’s ecosystems.

## Results

### An overview of the CheckAMG workflow

CheckAMG is an automated, scalable, and modular pipeline for identifying, curating, and classifying AVGs on viral or mixed metagenomic sequences (Figure 1). It operates as two complementary modules that can be run independently or together. First, the *annotate* module delineates AMG, APG, and AReG predictions for each open reading frame (ORF) in a scaffold. Each prediction comes with interpretable supporting evidence in CheckAMG’s output that combines (i) curated reference-based functional annotation against a set of protein profile HMM databases with labeled functions, (ii) a per-gene confidence score of how auxiliary-like the gene is (the AMG weight, detailed below), and (iii) an annotation-derived viral or nonviral gene origin classification (Supplemental Figure 1). Second, the *de-novo* module uses a finetuned protein-based genome language model, CheckAMG-PST, to identify viral and AVG-like proteins without relying on sequence homology to a reference database of proteins with labeled, specific functions (Supplemental Figure 2). By scoring proteins in a learned embedding space rather than against a fixed reference based on sequence homology, this module extends detection to auxiliary genes whose function is too divergent, or simply absent, from any curated database, which the annotate module cannot recover by design.

**Figure 1.**
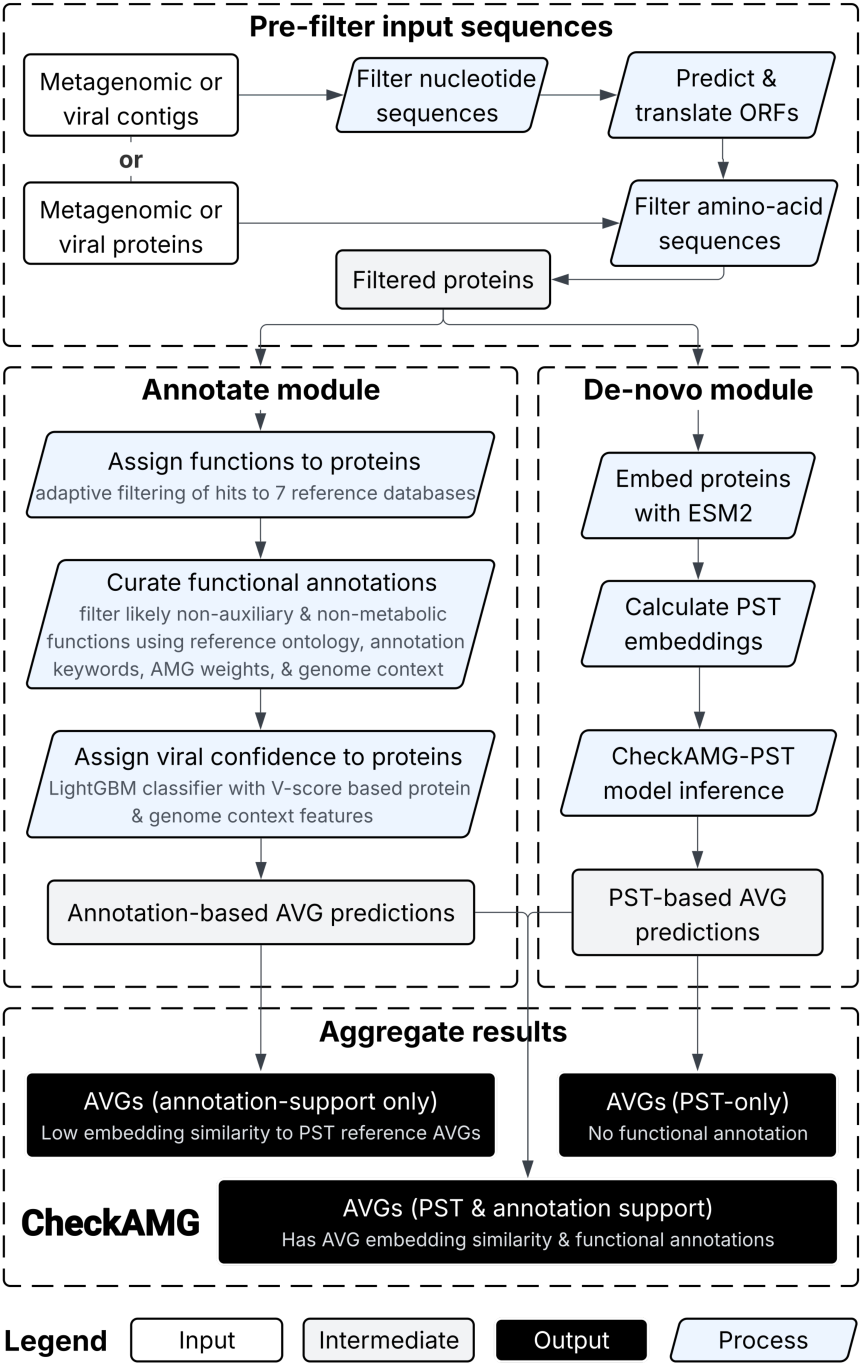
Parallel annotation- and deep-learning-based AVG identification in CheckAMG. Overview of the *end-to-end* workflow. Filtered proteins from metagenomes or viral genomes enter two parallel modules. The *annotate* module (left) assigns functions by adaptive HMM searches against seven reference databases (KEGG KOfam, FOAM, PHROGs, Pfam-A, dbCAN, CAMPER, and METABOLIC), curates putative auxiliary gene functions using AMG, APG, and AReG-specific rules (and AMG-weights for AMGs specifically, which measure how “virus-like” and “metabolic-like” an annotation is), and verifies that AVGs are virus-encoded. The *de-novo* module (right) embeds each protein with ESM-2, contextualizes the embeddings with a finetuned genome language model CheckAMG-PST, and infers viral and AVG-like probabilities by nearest-neighbor search against labeled training proteins. Predictions from the two modules are aggregated per gene into three reported classes: annotation-supported only, *de-novo*-only, and *de-novo* + annotation-supported. Either module can be run independently. Shapes denote input (rounded rectangles), intermediate data (white rectangles), processes (filled rectangles), and outputs (highlighted rectangles).

When invoked using the *end-to-end* workflow, the two modules run independently on the same input sequences, and their outputs are evaluated for each gene to assess concordance. This yields final AVG predictions that fall into one of three evidence classes: (i) AVGs supported by annotation alone, (ii) AVGs supported by both annotation and CheckAMG-PST embedding similarity, and (iii) *de-novo*-only AVGs that lack any functional annotation match in CheckAMG’s curated reference database (Figure 1). Either module can also be run on its own. The *annotate* module requires no GPUs and processed 48,660 proteins encoded by 3,175 scaffolds in 1.6 hours using 10 CPUs (or 0.6 hours when using 30 CPUs). It provides full AMG, APG, and AReG predictions with interpretable annotation-level evidence, making it best suited for datasets in which “known” AVGs are expected. In contrast, the *de-novo* module likewise requires no GPUs, with the same 48,660-protein input completing in 5.5 hours using 10 CPUs (2.7 hours using 30). However, the *de-novo* module is substantially accelerated using a GPU, as a run with 1,031,362 proteins encoded on 47,987 scaffolds completes in 12 hours using a single NVIDIA H200 GPU (and one CPU for overhead). The *de-novo* module is best suited to discovering candidate AVGs whose function cannot be inferred from sequence homology to existing reference proteins with labeled functions in the CheckAMG database. Together, these modules give CheckAMG a single framework that is both interpretable and capable of discovery, which neither annotation-only nor model-only approaches provide on their own.

We next benchmarked each component of CheckAMG in turn. For the *annotate* module, we evaluated the viral-origin classifier against held-out test datasets spanning near-pure viral to host- and MGE-dominated sequence compositions, and compared its AMG calls against the widely used tools DRAM-V^27^ and VIBRANT^28^. For the *de-novo* module, we evaluated the viral and AVG classifier against the same held-out test datasets and tested their ability to generalize to proteins and functions withheld entirely from model training.

### Is the functional protein of interest in a virus or is it host-derived?

The first step in identifying AVGs in metagenomic assemblies is assessing whether the functional gene of interest is present on a virus or a cellular genomic fragment, since a gene that is not of viral origin is not an AVG regardless of its function. However, predicting this for individual genes is challenging, even when they occur on scaffolds classified as overall viral by virus sequence identification tools^22^. This is because viral-versus-cellular genome boundaries can be blurred by imprecise prophage boundary predictions^22^, variously degraded prophage genome states^23^, nested mobile genetic elements (MGEs)^24^, and, more rarely, chimeric assembly^25^. As a step towards resolving these challenges, CheckAMG *annotate* includes a viral origin confidence classifier that evaluates the “viralness” of each protein in a scaffold via a Light Gradient Boosting Machine (LightGBM) model^35^ trained and tested to separate viral from non-viral proteins using annotation-derived genomic context.

To evaluate the “viralness” of proteins, the classifier integrates multiple complementary lines of evidence. One set of features (gene-level statistics; Methods) assesses the protein itself, quantitatively scoring how often the protein function is encoded by viruses broadly, captured by V- and V_L_-scores^33^ resulting from KEGG^36^, Pfam^37^, and PHROG^38^ HMM searches. Another set of features assesses the genomic context of the protein under consideration by checking whether it is surrounded by other viral or non-viral proteins. This signal is quantitatively captured by scaffold-level and local window averages of V_L_-scores (evaluated below). A third set of features records the distance to the nearest high-V-score (highly “virus-like”) annotation in the left and right flanks, as well as the distance to the nearest annotation associated with a nonviral mobile genetic element (MGE). Together, the lines of evidence captured by these features give CheckAMG *annotate* a quantitative metric to assess whether a functional gene of interest is genuinely viral or encoded within a non-viral genomic region, whether cellular, a putative nonviral MGE, or ambiguous, such as the boundary of a degraded provirus.

The classifier was trained and evaluated on 16,825,811 proteins encoded by 1,501,338 scaffolds drawn from two complementary ground-truth sources: (i) the fragmented viral and non-viral dataset used to train the virus finding software, geNomad^39^, and (ii) a set of bacterial and archaeal sequences from proGenomes v3^40^, a strictly curated database of consistently annotated prokaryotic genomes. Critically for the latter, we annotated MGE and prophage regions within these cellular genomes using the pangenome-based resource proMGE^24^. proMGE defines MGE boundaries using the presence of 68 recombinase/endonuclease subfamilies sourced from transposons, conjugative elements, integrons, phages, and nonmobile cellular subfamilies, together with pangenome-wide maps of accessory regions identified through comparative genomics. However, since the MGE boundaries defined by proMGE are, by design, overlapping and extend viral regions to “upper bounds” past chromosomal regions^24^, we developed a complementary “strict viral region” identifying algorithm to refine these boundaries. This strict algorithm is not enabled by default; its primary role is to refine ground-truth labels in the training and test data (Methods; Extended Data Figure 1).

The resulting training set was 46.9% host-chromosomal, 13.9% MGE, and 39.3% viral at the protein level. This class balance was chosen to provide reliable per-protein labels across all three source types rather than to mimic any single real-world sample composition, which we instead evaluate directly using the ten realistic-composition test sets below. Forty features, including per-protein and scaffold-level V- and V_L_-scores, distances to flanking high-V-score genes, and MGE-flank identity, were computed per protein from CheckAMG *annotate* outputs (Methods). Feature importance in the trained model was dominated by scaffold- and window-level average V- and V_L_-scores, with the scaffold-wide V- and V_L_-scores of Pfam, KEGG, and PHROG accounting for six of the seven most-used split features; per-protein V- and V_L_-scores and distances to flanking viral genes contributed secondarily (Supplemental Table 1).

We next tested the trained classifier under varied conditions across ten held-out datasets. These ten datasets were randomly sampled to span a broad range of user-encounterable situations, ranging from near-pure viral genomes to host- or MGE-dominated metagenomes, to a dataset of integrated proviruses where viral and non-viral genes were co-located on the same scaffold (Extended Data Figure 2A; Supplemental Table 2). Area Under the Precision-Recall Curve (AUPRC) ranged from 0.891 to 0.996 across eight of the ten sets and fell only on the two most challenging compositions, MGE-enriched (0.627) and near-all-host (0.726; Supplemental Table 2; Extended Data Figure 2B–C). The MGE-enriched dataset reflects a particularly challenging case that we deliberately included because gene flow between MGEs and viruses may limit how sharply a per-protein classifier can distinguish between them. Although classifier performance was lowest on this dataset, it retained useful predictive power, with an AUPRC exceeding the random-guessing baseline of 0.073, the share of viral proteins in that set. Moreover, this MGE-enriched composition represents an unusually extreme scenario that is unlikely to reflect typical virus-enriched or mixed-community metagenomic datasets, for which the relative abundance of MGEs is expected to be substantially lower than in this hypothetical test case^24,41^.

Because a single probability cutoff forces a tradeoff between missing true AVGs and admitting false ones, we next calibrated operating thresholds for practical use. Two probability thresholds partition the continuous classifier output into high, medium, and low-confidence tiers (thresholds selected as described in Methods). At these thresholds, the high-confidence tier reached a precision of 0.953-0.997 (the fraction of calls that are correct) and recall of 0.439-0.676 (the fraction of true-labeled AVGs recovered) across all ten held-out test datasets except the MGE-enriched and near-all-host sets (Supplemental Table 2; Extended Data Figures 2C, 3). The medium-confidence tier had a precision of 0.696-0.989 on the same eight datasets, with correspondingly higher recall. Notably, both tiers maintained precision above 0.90 even on the integrated provirus test set (high: 0.955; medium: 0.907), which was designed to represent a challenging case in which viral sequences are embedded within host chromosomes with both discrete and ambiguous boundaries. Model output probabilities tracked the true viral fraction, with the greatest calibration drift occurring in compositions most different from the training distribution (Extended Data Figure 2D). Together, these results show that the LightGBM classifier assigns reliable viral origin confidence to individual proteins across a wide range of metagenomic sequence compositions, although we recommend applying CheckAMG to viral scaffolds or vMAGs identified with high confidence by a reliable upstream tool to avoid permissive misuse on unscreened assemblies.

### Does the virus-encoded functional protein of interest encode an *auxiliary* function?

Once a functional protein of interest is assessed as likely viral, the next challenge is to evaluate whether its function most plausibly impacts host metabolism (AMG), physiology (APG), or gene regulation (AReG). To achieve this, CheckAMG *annotate* employs three separate lists of putative metabolic, physiological, and regulatory functions derived from seven pre-existing profile-HMM databases^36–38,42–45^ for making initial classifications (Supplemental Tables 3–5). After putative AMG, APG, and AReG assignments have been made, the next step is to distinguish auxiliary from non-auxiliary functions. This distinction matters because functional annotations are generally assigned within a cellular context and therefore do not by themselves indicate that a function is auxiliary to the virus. The same biochemical activity can serve different biological roles depending on context: a function that contributes to cellular metabolism, physiology, or gene regulation when encoded by a host may instead perform a core viral function when encoded by a virus. Core viral functions may likewise build the viral particle, facilitate genome replication or packaging, lyse the cell, or have other biochemical activities necessary to complete the viral infection cycle. In contrast, an auxiliary function acts on host metabolism, physiology, or gene expression to support that infection cycle indirectly. Thus, annotated functions that appear to be involved in host metabolism, physiology, or gene regulation require contextual curation to determine whether they represent an auxiliary versus core viral function^1^.

This contextual distinction is particularly important for AMGs, because databases of protein functions used to assign metabolic annotations encompass broad and overlapping biological processes^46,47^ and were not designed to distinguish host metabolic functions from viral auxiliary metabolism. To address this problem, CheckAMG *annotate* computes an “AMG weight” for AMG predictions that combines the metabolic specificity of an annotation (its metabolic ratio, derived from KEGG BRITE, FOAM, GO, and EC ontological classifications; Methods) with its viral rarity (a penalty derived from its V_L_-score^33^). AMG weights range from 0 to 1, with higher values indicating functions that are both strongly associated with metabolism and relatively uncommon among viruses broadly, whereas lower values indicate functions that are weakly associated with metabolism, relatively common among viruses, or both (Figure 2A). Accordingly, annotations with high V_L_-scores, indicating that they are broadly common across virus genomes, or low metabolic ratios, indicating broad or nonspecific metabolic associations, receive lower AMG weights even when they fall within a metabolic ontology.

**Figure 2.**
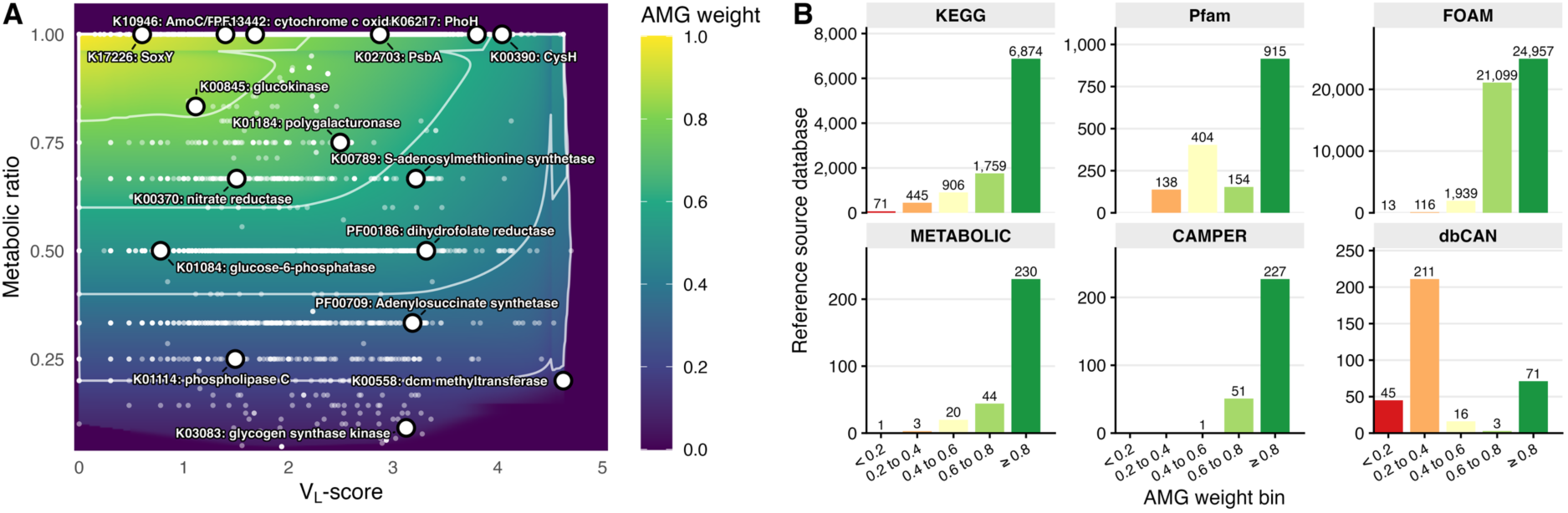
AMG weights balance metabolic specificity and viral rarity. **(A)** Relationship between metabolic ratio (the number of metabolism-like pathways to which an annotation maps divided by the total number of mapped pathways), V_L_-score (the gene’s “viralness”), and AMG weight for a subset of KEGG KO and Pfam profiles in the CheckAMG database. By default, CheckAMG *annotate* treats annotations with AMG weights of at least 0.6 as putative AMGs, which excludes annotations that are common in viral genomes (high V_L_-scores) and/or have broad or non-metabolic associations (low metabolic ratios). Annotations without V_L_-scores are not shown because their AMG weight equals their metabolic ratio. White contour lines mark AMG weights of 0.8, 0.6, 0.4, and 0.2. **(B)** Distribution of AMG weights across CheckAMG reference annotations, faceted by source database (KEGG, Pfam, FOAM, METABOLIC, CAMPER, dbCAN). PHROG annotations were not assigned AMG weights because they lack ontological mappings comparable to what the other databases have.

By default, CheckAMG *annotate* treats only annotations with an AMG weight of at least 0.6 as candidate AMGs, as this is a point at which candidate annotations become predominantly metabolic and rarely viral in the reference distribution (Figure 2A). This threshold was defined manually and is user-configurable. Applied to CheckAMG’s curated AMG reference database, the weights distribute clearly across the HMM databases queried (Figure 2B). At the 0.6 cutoff, KEGG contributes 8,633 KOs and Pfam contributes 1,069 families, with an additional 46,056 families from FOAM, and others from METABOLIC (274), CAMPER (278), and dbCAN (74) (Figure 2B). Most KEGG entries cluster at high weights (at least 0.8), consistent with their derivation from the narrow “Metabolism” BRITE hierarchy^36,47^, whereas Pfam entries span a broader range, reflecting the greater functional diversity of Pfam families^37^. CAZyme annotations from dbCAN, however, mostly fall below the 0.6 threshold. This distribution results from its abundance of glycosyltransferases and glycoside-hydrolases, which by design were considered as “non-metabolic” functions when calculating metabolic ratios, as viruses commonly encode these enzymes for core viral functions such as host cell entry^1^ (Methods). This weight distribution defines which annotations CheckAMG can emit at default settings, and provides an annotation-level scoring system for what constitutes a likely AMG. As shown below, we use this same system as an evaluative metric to compare the AMG calls of different tools with a common framework.

Although distinguishing auxiliary from non-auxiliary functions is equally important for APGs and AReGs, we did not develop a metric analogous to the AMG weight for these classifications. This is due to two limitations: (i) comprehensive, curated, and ontologically structured databases of physiological and regulatory functions are less developed than those available for metabolic functions, and (ii) fewer guidelines have been established for distinguishing core from auxiliary viral functions specifically for APGs and AReGs, which have only recently been distinguished from AMGs. Consequently, we cannot quantify the breadth of physiological or regulatory functions and pathways associated with an annotation using a standardized ontological framework, as is available for metabolic functions (and is a key component of the AMG weight). Instead, CheckAMG *annotate* uses curated lists of physiological and regulatory functions, together with APG- and AReG-specific rules, to reduce potential misclassification. This provides a qualitative approach to contextualizing these predictions in the absence of a comparable quantitative metric (Methods, Supplemental Tables 6, 7).

### CheckAMG’s AMG calls are more often viral than DRAM-V’s and more numerous than VIBRANT’s

To evaluate how differences in viral origin classification, functional annotation, and AMG curation affect AMG predictions, we benchmarked CheckAMG *annotate* against the two most widely used automated AMG tools, DRAM-V^27^ and VIBRANT^28^. We restricted the comparison to AMGs, since neither tool reports APGs or AReGs, and to the annotate module, since neither has a counterpart to the de-novo module. The three tools were run on mixed-community metagenomes, viromes sequenced from the same samples, and whole viral genomes from human gut, aquatic, and soil ecosystems, and received identical protein inputs so that differences reflect annotation and curation rather than ORF calling (Methods).

Because the three tools report AMG calls at different levels of confidence in viral origin, we compared them at matched, cumulative confidence levels (Methods). DRAM-V ranks viral context with its own rule-based auxiliary score (unrelated to the AMG weights and V-scores above), from 1, the most confident, to 5, the least, and its default output includes scores of at most 3, so DRAM-V (aux 3) and DRAM-V (default) are equivalent. Comparisons of confidence levels below match DRAM-V (aux 1) to CheckAMG (high) and VIBRANT (high), and comparisons of the output each tool returns by default use DRAM-V (default). At each tool’s most confident level, CheckAMG (high) and DRAM-V (aux 1) emitted similar numbers of AMG calls, 3,221 and 3,380, and VIBRANT (high) fewer than half as many, 1,270, whereas DRAM-V emitted 47,308 at its default setting (Figure 3A–B). Of the CheckAMG (high) calls, 57.6% (1,856) were also called by at least one other tool at some tier, but only 10.1% (325) at DRAM-V (aux 1) and 6.9% (221) at VIBRANT (high). The gap widened at more permissive settings, with 170,161 calls for CheckAMG (low), 12,677 for VIBRANT (low), and 171,833 for DRAM-V (aux 4, allow T), reflecting how differently each tool treats weaker viral evidence (Extended Data Figure 4).

**Figure 3.**
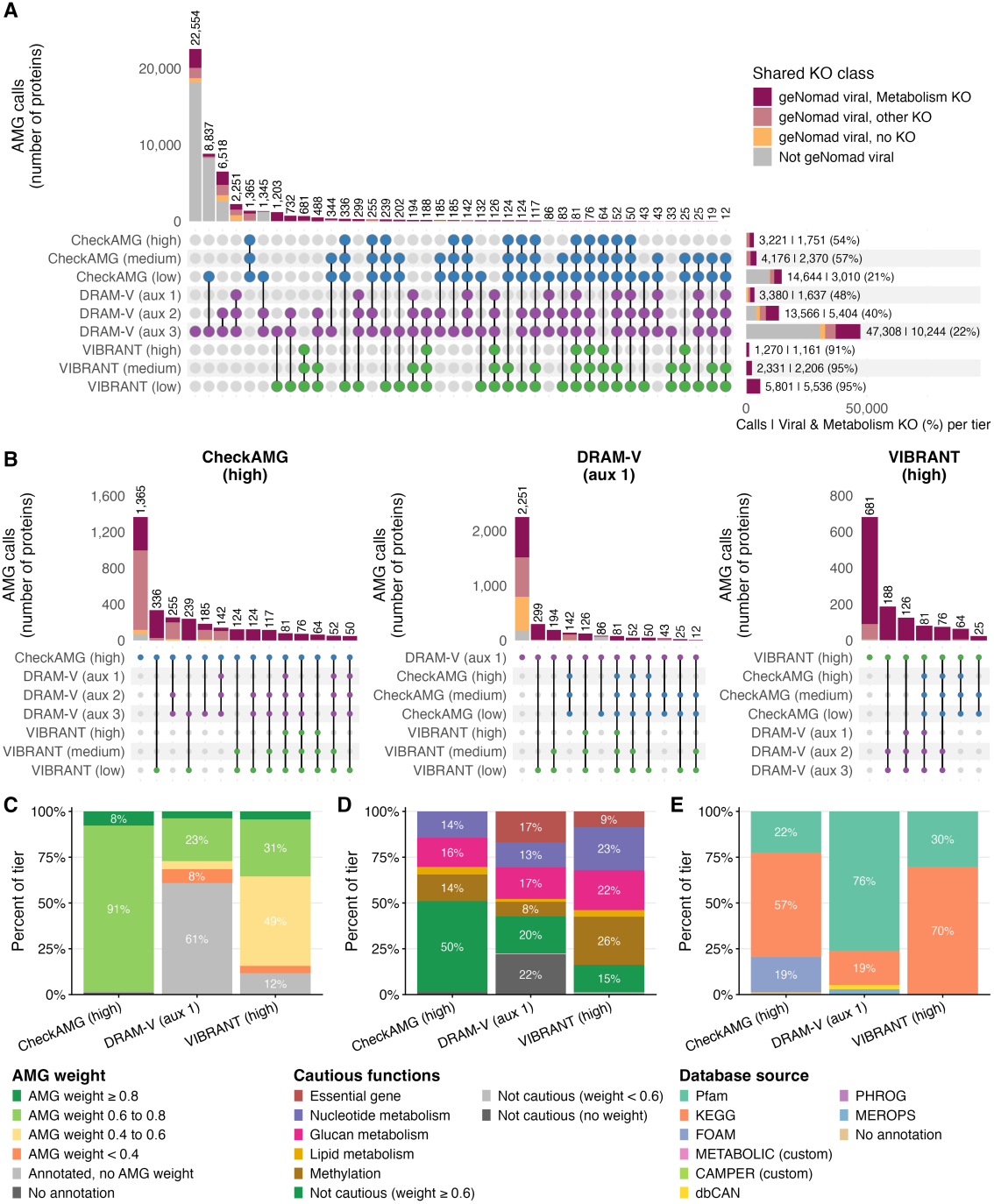
CheckAMG, DRAM-V, and VIBRANT produce mostly distinct AMG predictions. **(A)** Overlap of AMG calls across the confidence tiers of each tool. Rows are the tiers and columns the calls shared by exactly the tiers marked with connected dots. All tiers are cumulative, each containing the calls of the stricter tiers of the same tool. DRAM-V ranks its calls with a rule-based auxiliary score, on which 1 (aux 1) is the strongest support and 3 (aux 3) is the weakest still emitted under default settings; thus DRAM-V (aux 3) and DRAM-V (default) are equivalent. CheckAMG (high) are the AMG calls that CheckAMG emits under default settings. VIBRANT does not separate calls by viral confidence on a per-gene basis like CheckAMG and DRAM-V, so here we use VIBRANT (high) as its matched tier. Dots are colored by tool, CheckAMG blue, DRAM-V purple, VIBRANT green. Intersection bars are split by whether separate evaluations of the same proteins also identified them as being encoded by a virus (geNomad viral) and encoding a function considered metabolic by its KofamScan best hit (Metabolism KO; Methods), serving as a separate classification for evaluating AMG calls on common ground (Shared KO class). Only genes called by at least one tool at CheckAMG (high), DRAM-V (default) or VIBRANT (high) are shown, horizontal bars give the calls per tier among those genes, and the labels to the right of them show the number of these calls and just those that are geNomad viral with a metabolic KO. For the full, unfiltered set of AMG calls at all tiers, including lower-confidence calls outside these tiers, see Extended Data Figure 4. **(B)** The intersections of (A) separated by each tool and restricted to the AMG calls made at its most confident tier, CheckAMG (high), DRAM-V (aux 1) or VIBRANT (high), showing which of those calls the other tools recover at any tier. Bars and dots are colored as in (A), and intersections below 12 calls are omitted for simplicity. **(C)** AMG weights of the calls at the tiers of (B), assigned by CheckAMG during annotation, with unmatched calls labeled “Annotated, no AMG weight.“ Calls with multiple annotations take the one with the highest mapped AMG weight. **(D)** Share of the calls in (C) carrying an annotation in a CheckAMG “cautious function” category, with unflagged calls split by AMG weight and multi-category calls assigned the category least common in their tier. **(E)** Database source of the representing annotation for the calls in (C), the highest-weight annotation or, where none carries a weight, the strongest hit.

To compare the AMG calls themselves on common ground, we evaluated them on two axes independent of CheckAMG’s own definitions. First, we used geNomad^39^ to independently evaluate the viral origin of every AMG call. Among these, 3,153 of 3,221 CheckAMG (high) calls (97.9%) were geNomad viral, against 3,135 of 3,380 DRAM-V (aux 1) calls (92.8%), 1,263 of 1,270 VIBRANT (high) calls (99.4%), and only 16,746 of the 47,308 DRAM-V (default) calls (35.4%) (Figure 3A–B, Extended Data Figure 4, Supplemental Tables 8 and 9). The same order held at the medium levels, with 86.8% for CheckAMG (5,843 of 6,728), 70.0% for DRAM-V (aux 2) (9,490 of 13,566), and 99.7% for VIBRANT (3,708 of 3,720), while CheckAMG (low), a set 3.6 times larger than DRAM-V (aux 3), fell to 5.2% (8,879 of 170,161). Second, we reannotated every call with KofamScan^48^ and classed each by whether its best KO fell under the KEGG “Metabolism” category (Metabolism KO, Methods). This criterion is not inherent to CheckAMG, although alone it is neither necessary nor sufficient to confirm a putative AMG as likely^1^. Among geNomad-viral calls, the share with a Metabolism KO was similar for CheckAMG and DRAM-V at every matched level, 55.5% against 52.2% at the most confident (1,751 of 3,153 and 1,637 of 3,135). VIBRANT stayed near 92% throughout (1,161 of 1,263 at high), since it draws its AMG calls only from annotations in the KEGG metabolism category, with separate cutoffs for its HMM-based hits^28^.

A Metabolism KO also misses functions that KEGG places outside its Metabolism category. Of the 1,402 geNomad-viral CheckAMG (high) calls without a strict Metabolism KO, 997 had a KO in the KEGG “Unclassified: metabolism” or “Membrane transport” categories, against 316 of 1,498 for DRAM-V (aux 1) (Supplemental Table 9). The Shared KO class in Figure 3A–B therefore understates the metabolic functions of CheckAMG’s calls more than those of DRAM-V’s. We therefore also compared how “auxiliary like” the calls of each tool were using CheckAMG’s AMG weight, which was possible for protein families in both CheckAMG’s reference AMG database and that of DRAM-V or VIBRANT (54% of DRAM-V references and 85% of VIBRANT references, Extended Data Figure 5). At the AMG weight minimum of 0.6 defined above for “likely” AMGs, 98.7% of CheckAMG (high) calls mapped to likely AMGs (3,178 of 3,221) (Figure 3C). The remaining 1.3% of CheckAMG (high) calls had a highest-scoring annotation from a protein family without a functional label, such as in PHROGs, but also carried a FOAM, KEGG, or Pfam annotation with a weight of at least 0.6 (Supplemental Tables 8, 9). For DRAM-V, 27.1% of aux 1 calls (915 of 3,380) and 37.4% of default calls (17,694 of 47,308) mapped to likely AMGs (Extended Data Figure 6B, Supplemental Tables 8, 9), as did 35.4% of VIBRANT (high) calls (450 of 1,270). Because CheckAMG annotate pre-filters AMG candidates at a weight of at least 0.6 (Methods), its near-100% share largely reflects that design choice. The lower shares for DRAM-V and VIBRANT, which do not filter by AMG weight, reflect how each tool defines putative AMGs and are not, by themselves, evidence of prediction error.

We next compared the tools against a separate, intentionally stringent set of criteria, CheckAMG’s list of cautious functions that may warrant additional scrutiny when classified as AMGs^1^ (essential viral genes, nucleotide metabolism, glucan metabolism, lipid metabolism, and DNA methylation; Supplemental Table 7). CheckAMG filters only the essential viral gene category by default, so the other categories offer a common comparison, and DRAM-V and VIBRANT calls carry a flag only if their annotation maps to the list. Of the CheckAMG (high) calls, 49.0% carried at least one cautious flag, compared to 57.3% for DRAM-V (aux 1) and 83.7% for VIBRANT (high), the latter driven substantially by methyltransferase annotations (Figure 3D). At its default setting DRAM-V’s share was 39.2%, and among geNomad-viral calls it was 55.4% against 49.1% for CheckAMG (high) (Extended Data Figure 6C, Figure 3 Source Data).

Overall, CheckAMG’s levels offer a usable trade between confidence and reach that the other two tools do not (Extended Data Figure 6A). Almost all of CheckAMG’s default calls are viral (97.9%), its medium level roughly doubles the call count while keeping 86.8% viral, and its low level is an explicit recall net, at 5.2% viral across 170,161 calls. DRAM-V’s viral share instead falls from 92.8% at aux 1 to 70.0% at aux 2 and 35.4% at its default. VIBRANT holds the highest viral share at every level by calling AMGs only within scaffolds it has already judged viral, but recovers the fewest calls, and its near-constant metabolic share is definitional rather than evidential. CheckAMG is therefore the only one of the three tools whose default output is both comparable in size to DRAM-V’s most confident set and nearly as viral as VIBRANT’s.

### Reference databases, viral-origin criteria, and curation rules each constrain cross-tool agreement

We next asked why the tools disagree on individual calls. For every AMG call made at a tool’s default or most confident tier, we classified why each of the other tools did not make the same call (Extended Data Figure 7A, Supplemental Results), which identified three sources of divergence. The first is which reference databases each tool searches. DRAM-V’s AMG calls rest mostly on Pfam, with 2,568 of its 3,380 aux 1 calls (76.0%) and 26,823 of its 47,308 default calls (56.7%) represented by a Pfam annotation, and VIBRANT’s rest on KEGG, with 886 of its 1,270 high calls (69.8%) represented by a KEGG annotation. The 3,221 CheckAMG (high) calls drew mainly on KEGG (1,839), Pfam (723), and FOAM (614), and FOAM is absent from the databases used by DRAM-V and VIBRANT (Figure 3E, Extended Data Figure 5C). However, this affected relatively few calls. Only 130 of the CheckAMG (high) calls (4.0%) were missed by DRAM-V and 119 (3.7%) by VIBRANT because their best hit came from a database the other tool does not search (Extended Data Figure 7A).

The second source of divergence is how each tool decides that an AMG call is viral rather than host-encoded. DRAM-V uses annotations from an independent tool, VirSorter2^49^, in a rule-based decision tree that scores how confidently each call is viral^27^. VIBRANT accepts metabolic genes on any scaffold it identifies as viral, whereas CheckAMG scores the viral origin of each gene as a continuous probability with the classifier described above. As a result, DRAM-V passed 204 of the 3,221 CheckAMG (high) calls (6.3%) only at its lower-stringency settings (auxiliary score 4 or the T flag), and VIBRANT called 1,053 (32.7%) only at its medium or low tiers, which reflect genome quality rather than per-gene viral evidence. Nearly all of these genes were geNomad viral, 193 of the 204 (94.6%) and 1,050 of the 1,053 (99.7%) (Extended Data Figure 7A–B). The third, and largest, source of divergence is which annotated functions each tool admits as an AMG, as defined by its AMG reference list. These lists share few entries across tools. CheckAMG’s curated AMG reference at weight at least 0.6 contains 8,633 KEGG KOs and 1,069 Pfam families. VIBRANT’s contains 2,826 KOs and no Pfam families, and DRAM-V’s only 34 KOs and 261 Pfam families (Extended Data Figure 5A). Only 14 KOs are shared across all three tools and 22 between DRAM-V and CheckAMG, and 963 of CheckAMG’s 1,069 Pfam families are absent from DRAM-V’s reference (Extended Data Figure 5B). This sets a hard ceiling on cross-tool agreement, since most functional identifiers that CheckAMG can use to support an AMG call are not curated as candidate AMGs by either competing tool, so no parameter change or relaxation of a tool’s criteria can recover them.

Differences in what each tool admits as an AMG were most evident at the level of individual calls. Most calls made by only one tool were annotated with the same database entry by the other tool, which then declined to call them as AMGs (Extended Data Figure 7A, Supplemental Results). DRAM-V declined about half of the CheckAMG (high) calls this way, 1,531 of 3,221 (47.5%), as did VIBRANT, 1,578 (49.0%). CheckAMG declined an even larger share of the other tools’ calls, 2,412 of the 3,380 DRAM-V (aux 1) calls (71.4%) and 991 of the 1,270 VIBRANT (high) calls (78.0%). AMG weights distinguished these two groups of calls. Most of the DRAM-V and VIBRANT calls that CheckAMG did not make had a weight below 0.6 or no weight in its reference, 1,960 of 2,412 (81.3%) and 765 of 991 (77.2%). In contrast, nearly all of the CheckAMG calls missing from the other tools had a weight of at least 0.6, 1,527 of 1,531 (99.7%) and 1,571 of 1,578 (99.6%). DRAM-V and VIBRANT therefore admit many functions that CheckAMG weights as unlikely AMGs, yet they leave out many that CheckAMG weights as likely.

Taken together, these analyses identify three distinct sources of divergence among AMG prediction tools: the reference databases each tool searches, its viral-origin criteria, and which annotated functions its curation rules admit as AMGs, with the last accounting for most disagreement. CheckAMG can recognize functional families that are absent from the DRAM-V or VIBRANT reference sets, applies annotation-level curation that favors functions with stronger metabolic specificity and lower viral prevalence, and identifies proteins that the other tools call only at lower-stringency settings despite evidence supporting their viral origin. These results support the value of integrating functional annotation, viral-origin confidence, and genomic context when identifying candidate AMGs, while also showing that differences among tools reflect both their reference databases and the criteria used to interpret viral-associated functions.

### A protein-based genome language model for annotation-independent AVG discovery

The results above depend on reference-database-dependent functional annotation, which cannot detect AVGs whose function has no match in a reference database. To identify AVGs independently of reference protein databases, CheckAMG includes a *de-novo* module built on a finetuned Protein Set Transformer (PST)^34^, a protein-based genome language model that takes ESM-2^31^ embeddings of all protein translations from predicted open reading frames on a scaffold and produces context-aware per-protein embeddings (CheckAMG-PST; Figure 1; Supplemental Figure 2).

CheckAMG-PST was finetuned from the pre-trained PST-TL-P large model checkpoint with two triplet-loss objectives. The first jointly classifies proteins by viral origin and AVG-like function within a four-class embedding space: non-viral/non-AVG, viral/non-AVG, non-viral/AVG, viral/AVG. The second incorporates a local genomic context objective to improve discrimination between the viral/non-AVG (common) and viral/AVG (rare) proteins by contrasting multiple overlapping contextual views of the same labeled protein. Importantly, the “AVG” label denotes a curated metabolic, physiological, or regulatory function that is not considered core-like, derived from CheckAMG annotate predictions without any filtering by viral origin confidence (Methods), regardless of whether the protein itself is viral or non-viral. This labeling scheme provides both viral/AVG and non-viral/AVG class examples so the model learns AVG-like function independently of viral origin. Each contextual view is anchored on the labeled AVG (or non-AVG) protein of interest but contains a different set of neighboring proteins, allowing the model to learn which upstream and downstream contextual information is most informative for distinguishing AVG-like proteins from similar non-AVGs. Because the contextual views are defined relative to the protein of interest rather than requiring complete genomic context, the model can also make use of the limited neighborhood information available on short or fragmented scaffolds. At inference, CheckAMG-PST emits two complementary per-protein probabilities: a viral probability and an AVG-like probability, calculated by distance-weighted voting over the *k* nearest training proteins in the learned embedding space (Methods). Their product is the final AVG probability, capturing the joint probability that a protein is both virus-encoded and has an AVG-like function, and therefore is an AVG.

Both training labels have defined authorities. The viral-origin label comes from the curated genome provenance of the geNomad training and benchmarking set, in which each fragment is labeled from the genome it was cut from rather than from any tool’s prediction, while the AVG-like label comes from CheckAMG annotate’s own curated function calls, assigned without reference to viral origin or genomic context (Methods). CheckAMG-PST was evaluated on the same ten test datasets used for the LightGBM classifier. For the viral classifier, AUROC and AUPRC were near-perfect across the seven most balanced-to-virus-enriched splits (AUPRC 0.981-0.998), and dropped only on the three hardest splits, MGE-enriched (AUPRC = 0.715), near-all-host (AUPRC = 0.937), and host-enriched (AUPRC = 0.975; Figure 4A–B). For the AVG-like classifier, we evaluated it as a soft “AND” logic gate against the viral label (i.e., a protein is only classified as an AVG if it is also classified as viral), and AUPRC ranged from 0.501 to 0.763 across all splits, with the PR curves separating cleanly at the chosen thresholds (Figure 4D–E). Both probabilities were conservatively calibrated on host- and MGE-heavy splits (Figure 4C, F). These results show that CheckAMG-PST learns an embedding space in which viral and AVG signal can be retrieved by nearest-neighbor search, and that combining its viral and AVG-like probabilities as a soft AND gate produces conservative AVG calls that stay reliable as the input composition shifts away from the training distribution.

**Figure 4.**
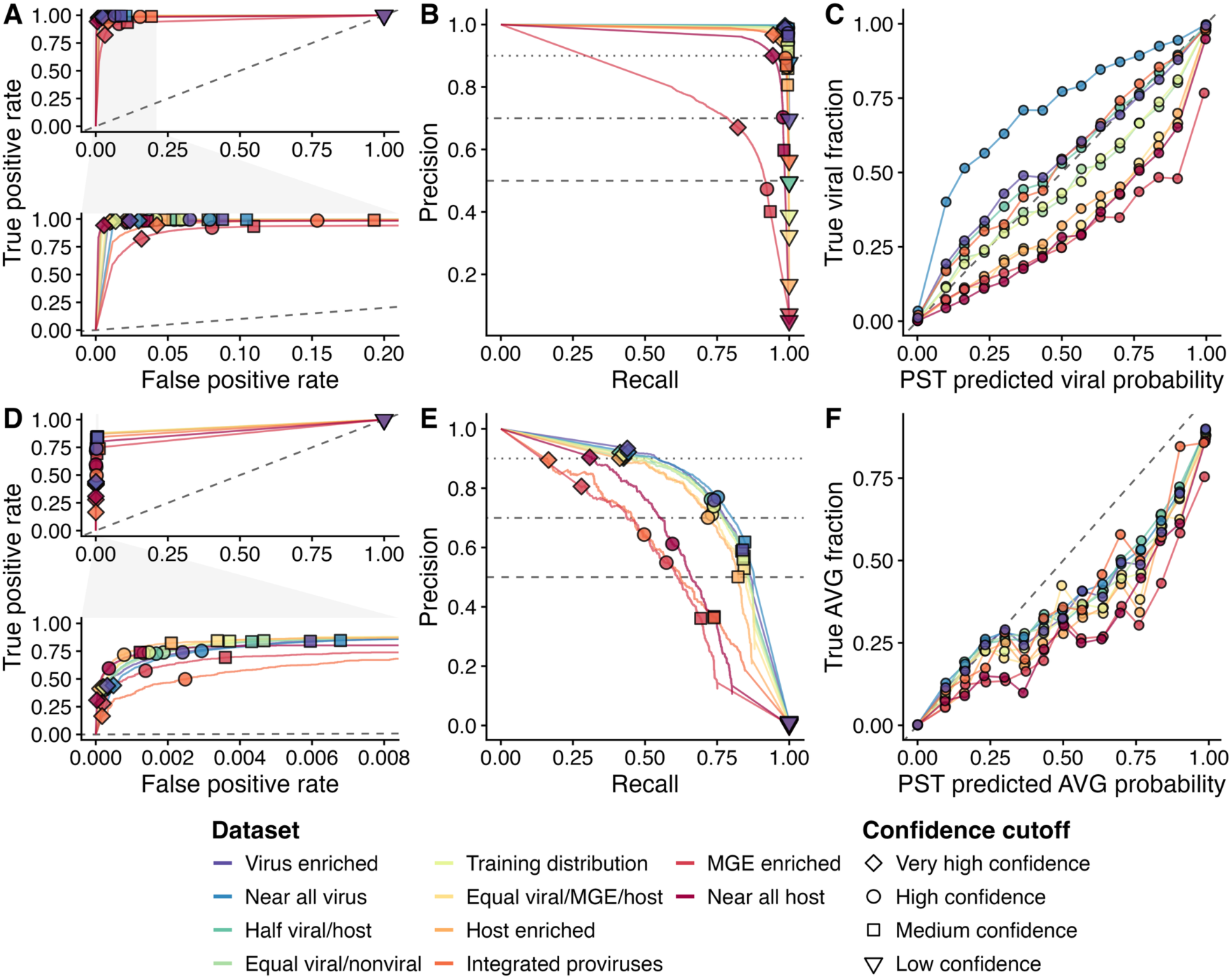
Performance of CheckAMG-PST viral and AVG classifiers across test sets spanning a range of viral, host, and MGE compositions. Markers denote operating thresholds for very high (diamond), high (circle), medium (square), and low (inverted triangle) confidence. **(A)** Receiver operating characteristic (ROC) curves for the viral classifier. The dashed diagonal indicates random performance, and the inset spans the operating range of high- and medium-confidence calls. **(B)** Precision–recall (PR) curves for the viral classifier. Horizontal lines mark precision of 0.90, 0.70, and 0.50. **(C)** Calibration of the viral classifier (15 bins per dataset; dashed diagonal is perfect calibration). **(D)** PR curves for the AVG classifier. **(E)** ROC curves for AVG classifier. **(F)** Calibration of the AVG classifier.

### CheckAMG-PST viral classification is more sensitive than the sequence similarity-based LightGBM classifier

To compare the sequence similarity-dependent (LightGBM) and the sequence similarity-independent (CheckAMG-PST) classifiers, we compared their per-protein viral probabilities across the test datasets (Extended Data Figure 8A). The two agreed strongly overall (Pearson r = 0.889, Spearman rho = 0.837), most closely on balanced and virus-enriched splits and least on near-all-host. At the medium-confidence threshold, CheckAMG-PST called 850,618 proteins viral (precision = 0.942, recall = 0.996) and the LightGBM called 835,300 (precision = 0.906, recall = 0.941; Extended Data Figure 8B). Of these, 774,761 proteins (91.1%) were called viral by both classifiers (precision = 0.975, recall = 0.940). Calls unique to CheckAMG-PST (75,857) retained majority-correct precision of 0.600, whereas calls unique to the LightGBM (60,539) were overwhelmingly incorrect (precision = 0.019), because CheckAMG-PST can recognize viral proteins by embedding-space similarity independent of annotation. CheckAMG-PST was also robust to scaffold length. LightGBM probability and accuracy rose steeply with gene count (Spearman rho 0.37; logistic slope 0.41 per log gene), whereas CheckAMG-PST was essentially length-invariant (rho 0.10; slope -0.005; Extended Data Figure 8C–G). In short, CheckAMG-PST viral classification is more sensitive than the LightGBM and more robust to scaffold length, while the two largely agree on confident calls. This supports running both modules together by default, preferring CheckAMG-PST calls, while retaining the interpretable LightGBM classifier when GPUs are unavailable or CPU scalability is limited.

### Application of CheckAMG to one million uncultivated soil and human-gut viral genomes

Applied at scale, CheckAMG recovered hundreds of thousands of AVGs, including many that homology-based methods cannot detect. We ran the *end-to-end* workflow on soil and human-gut viral genomes from the Meta-virus resource (MetaVR)^19^. MetaVR is a collection of viral genomes that were computationally identified and quality-filtered from public shotgun metagenomes, viromes, metatranscriptomes, and isolate genomes in a standard manner, based on best-practices developed in the field of viral genomics. After we performed additional quality filtering to further reduce the risk of analyzing nonviral sequences^50^, 767,946 soil and 238,034 human-gut viral genomes were retained as inputs for this analysis (Methods). The *annotate* module identified 174,200 AVGs in soil viral genomes at medium or high viral origin confidence (40,129 AMGs, 21,958 APGs, and 112,113 AReGs) and 177,614 in human-gut genomes (20,118 AMGs, 28,695 APGs, and 128,801 AReGs; Figure 5A; Supplemental Table 10). The *de-novo* module identified 253,379 AVGs in soil and 230,786 in gut genomes (484,165 total at medium confidence or higher; Figure 5A). Of these, 189,385 were not called by CheckAMG *annotate*, with 109,107 (57.6%) lacking any functional prediction of any kind (Figure 5B), encoded by 94,060 viral genomes. In the following sections, we further explore whether putative functions could be assigned to these 109,107 “sequence-similarity invisible” AVGs, and thus determine whether these are false positives or actual likely AVGs that could not be detected with sequence homology searching.

**Figure 5.**
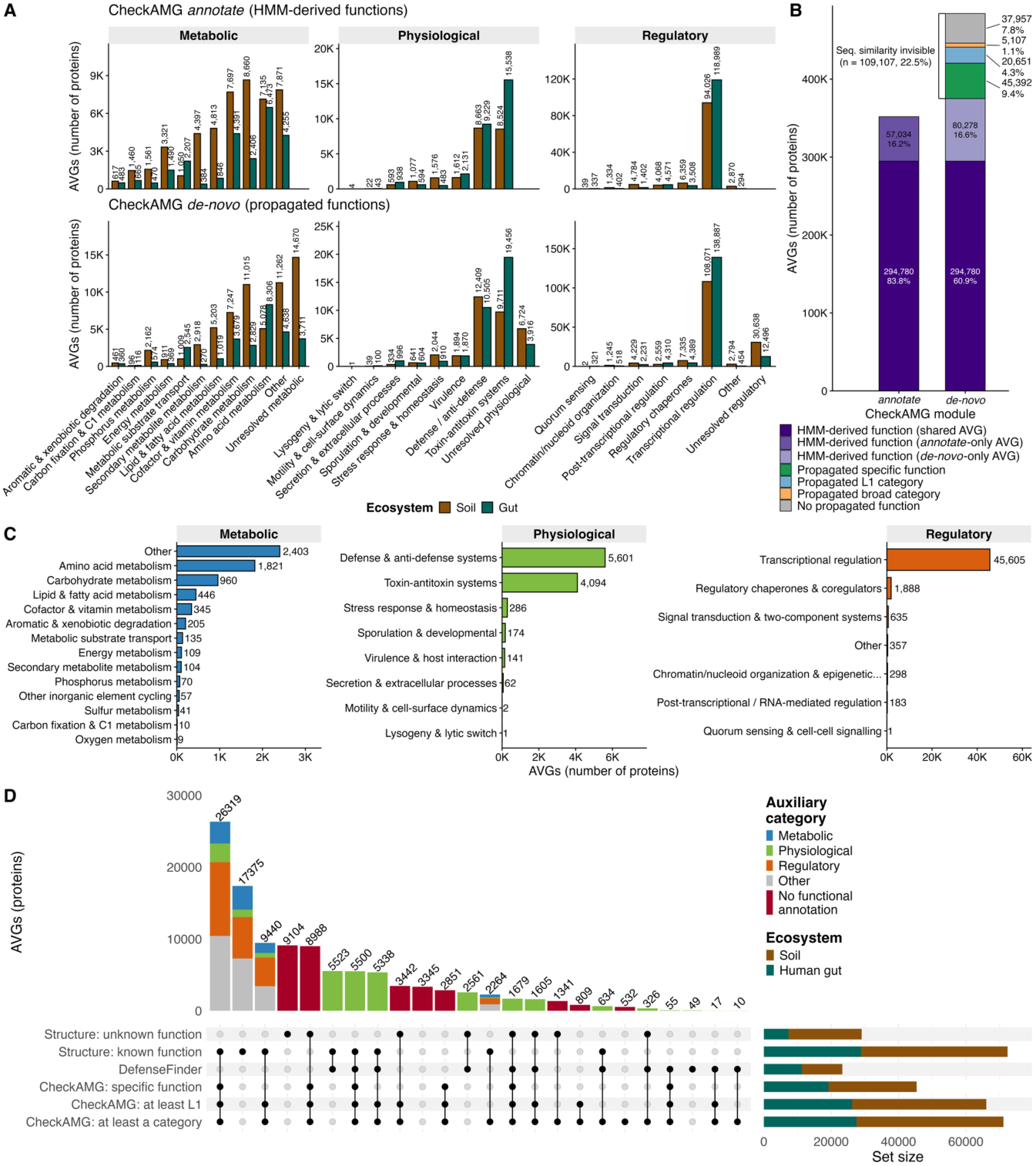
Auxiliary metabolic, physiological, and regulatory genes encoded by globally distributed human-gut and soil viruses. **(A)** Number of AMGs, APGs, and AReGs identified by the *annotate* and *de-novo* modules in each biome (medium confidence or higher), by CheckAMG’s “L1” function category. The functions for AVGs identified by the *de-novo* module were assigned by nearest-neighbor propagation in CheckAMG-PST embedding space (Methods). AVGs without propagated functions and only a broad metabolic, physiological, or regulatory category assignment fall under the “unclassified” functions. An AVG can be assigned to > 1 category, and each category an AVG was assigned to is counted. **(B**) The number and proportion of AVGs identified by each CheckAMG module. Proteins that were assigned homology-based functions by CheckAMG annotate are categorized as “HMM-derived function.” Proteins with HMM-derived functions called as AVGs by both modules are labeled as “shared AVG”, while proteins with HMM-derived functions flagged as an AVG by only one module are also shown (*annotate*-only and *de-novo*-only). Proteins called as AVGs by the *de-novo* module only that lack any HMM-derived function are labeled as “sequence-similarity invisible” and are further categorized by the level of function able to be assigned via nearest-neighbor propagation. **(C)** Auxiliary functions propagated to sequence-similarity invisible AVGs reported in (B), reported at the L1 function category level and faceted by broad metabolic, physiological, or regulatory classifications. **(D)** The 109,107 *de-novo*-only AVGs with no *annotate* HMM hit in any reference database (sequence-similarity invisible AVGs). Columns represent the number of proteins with intersecting functional classifications provided by annotations to DefenseFinder/AntiDefenseFinder reference, alignments against protein structures, and propagation of CheckAMG-based annotations using sequence- and embedding-based clustering. Structural matches are split by whether the matched structure has a known or an unknown function label. Set-size bars are colored by the ecosystem from which the genome encoding the sequence originated.

### Protein and embedding clustering assigns functions to most sequence-similarity invisible AVGs

Since 109,107 proteins classified as AVGs by the *de-novo* module carried no corresponding functional annotation from the *annotate* module (Figure 5B), we next evaluated whether functions can be accurately transferred from labeled proteins in the CheckAMG training dataset using either protein sequence-based or CheckAMG-PST embedding-based clustering, or both. Functional labels were propagated in two tiers: from the majority label of their protein sequence cluster where the cluster contained a labeled protein (the protein cluster tier), and otherwise from the nearest labeled training protein in CheckAMG-PST embedding space (the embedding tier), requiring each approach to meet a minimum 90% precision target when validated on held-out proteins (n = 41,957; Methods).

The protein cluster tier on its own labeled 89.4% of held out proteins at 94.4% precision, and the embedding tier alone labeled 90.0% at 86.5% precision (Extended Data Figure 9A; Supplemental Tables 11, 12). Protein clustering was therefore the more precise rule where it applied, and at specific depth it could reach 24,320 of the 41,957 held-out proteins against 9,714 for the embedding tier (Extended Data Figure 9B). Its reach nonetheless fell with sequence identity to the labeled set. Among the 468 held-out proteins below 30% identity to their nearest training protein, the cluster tier assigned no specific function at all, because at the 50% identity threshold deployed for specific names none of these proteins shared a cluster with a labeled training protein (Methods). The 30% threshold used for the two category levels did reach 84 of the 468 (17.9%) at 95.2% precision for L1 categories and 97.6% for broad categories (Supplemental Table 11). The embedding tier reaches every protein before its calibrated precision gate is applied (Methods), and after the gate it assigned a specific name to 48 of the 468 at 87.5% precision, an L1 category to 57 at 94.7%, and a broad category to 125 at 84.0% (Supplemental Tables 11, 12). Combining them, so that the embedding tier reaches only the proteins sequence clustering could not, labeled 92.2% of the same held-out proteins at 93.7% precision, at the same 90% precision target (Extended Data Figure 9A). The label-transfer accuracy was not a product of the fine-tuned CheckAMG-PST model itself. At matched sample size, the unmodified ESM-2 backbone transferred labels more accurately than CheckAMG-PST at every level of the functional hierarchy (Extended Data Figure 9C; Supplemental Results). Yet, overall, the two-tier approach labels more proteins than either alone, at a precision matching the more precise of the two.

Therefore, we applied the two-tier approach to propagate functions to the sequence similarity-invisible AVGs. This assigned some level of function to 71,150 (65.2%) of the 109,107, with 45,392 (41.6%) receiving a specific function, 20,651 (18.9%) a high-level function (“L1” classifications, as defined by CheckAMG *annotate*), and 5,107 (4.7%) only a broad metabolic (14.9% of this group), physiological (15.1%), or regulatory (70.0%) category. The remaining 37,957 (34.8%) received no label (Figure 5B-C; Supplemental Table 10; Supplemental Results; Extended Data Figure 9D). Of the labeled proteins, 26,544 (24.3% of the 109,107) took their label from the protein sequence cluster tier and 44,606 (40.9%) from the embedding tier, with the embedding tier supplying 70.1% of the specific function names and the cluster tier 62.6% of the L1 categories (Extended Data Figure 9E; Supplemental Tables 11, 12). Overall, our deep-learning classifier recovered over 100,000 auxiliary genes invisible to sequence homology search, underscoring how much AVG diversity remains beyond the reach of annotation-only tools.

### CheckAMG-PST discovers families of novel AVGs

We next asked whether the sequence-similarity invisible AVGs identified by CheckAMG *de-novo* are readily identifiable by other annotation approaches. We first searched predicted protein structures for every sequence-similarity invisible AVG against public databases including structures with viral and nonviral origins. The structural alignments matched 72,393 proteins (66.4%) to a structure with a specific or named function, and another 29,046 (26.6%) to a confident structural match whose target had an unknown function (Figure 5D). The remaining 7,668 (7.0%) did not have a structural match at all. After grouping the named matches by functional theme into metabolic, physiological, and regulatory categories, 32,109 (44.4%) were regulatory, 9,090 (12.6%) metabolic, and 5,516 (7.6%) physiological, with the remaining 25,678 (35.5%) falling outside these three categories (Supplemental Table 13).

The named matches spanned 5,458 distinct function names, of which the ten most common accounted for 22.0%, and helix-turn-helix transcriptional regulators were the most frequent, led by the XRE family (3,878 proteins), with toxin-antitoxin systems and nucleases dominating the physiological matches. Methyltransferases were the largest single class within the metabolic theme (2,288 of 9,090), a class CheckAMG flags as warranting additional scrutiny in an AMG call and one that a structural match alone cannot resolve into a host-metabolic or a genome-modifying activity (Supplemental Results). These are the functional classes expected of genes that act on the host during infection rather than of core viral genes, which further supports the *de-novo* only AVGs as likely genuine AVGs rather than false positive predictions. Furthermore, because a substantial portion of AVGs were assigned defense and anti-defense functions (Supplemental Table 10; Figure 5A), we also searched the sequence-similarity invisible AVGs against models in the DefenseFinder and AntiDefenseFinder database^51^. We identified 23,297 proteins (21.4% of the 109,107) with matches, of which 14,778 passed the database-provided confidence thresholds, while the remaining 8,519 were detectable only with a more permissive cutoff (Figure 5D).

Together, DefenseFinder and alignments to structures with known functions could identify 78,695 of the 109,107 sequence-similarity invisible AVGs (72.1%; Figure 5D). The remaining 30,412 (27.9%) could not be assigned any putative function with DefenseFinder or structural alignments. Among these remainders, the two-tier functional propagation approach described above assigned at least a broad metabolic, physiological, or regulatory classification to 59.1%, an L1-level function to 52.9%, and a specific function to 38.9%. The remaining 12,449 (40.9%) had no propagated function at any level (Figure 5 Source Data). In other words, 12,449 *de-novo*-predicted AVG proteins could not be assigned a function of any kind by structural alignment to a large and diverse set of references, by matching to DefenseFinder references, or by our dual protein sequence- and embedding-based clustering approach.

Protein sequence clustering was not effective at resolving broad families for the 30,412 AVGs without matched functions. Proteins spanned 13,419 distinct clusters at 30% identity and 80% coverage, and 8,740 (28.7%) fell in a cluster that was a singleton across the full 1,500,646-protein set, so they had no sequence relative at that threshold. Clustering CheckAMG-PST embeddings grouped those 13,419 clusters into 106 families, of which 50 also contained proteins that DefenseFinder or structural alignment identified and held 30,312 of the 30,412, while 56 contained no identified member and held the remaining 100 (Figure 5 Source Data). Most of these proteins were therefore divergent members of families whose other members carry an independently identified function, rather than isolated sequences. Considering that a specific function could not be obtained for most of them by sequence-based, structure-based or embedding-based approaches, while a broad auxiliary category could be assigned to the majority, we suggest that the auxiliary genes lacking specific functional annotations found only by CheckAMG *de-novo* represent genuinely uncharacterized auxiliary functions, and are a source of auxiliary viral “dark matter” that remains to be described.

### CheckAMG reveals soil- and human gut-specific AVG functions

To resolve broadly coherent, ecologically interpretable groups of related AVGs, we clustered the CheckAMG-PST embeddings of all identified AVGs using HDBSCAN density-based clustering with a minimum cluster size of 300 protein embeddings. This yielded 173 clusters, most of which showed at least some skew toward one biome and a minority that were strongly skewed towards either soil or the human gut (Figure 6A). Several of the biome-skewed AVG functions match the ecology of their environment. For instance, soil-specific clusters were dominated by host transcription and cell-cycle regulators (for example, transcription factor WhiB, GapR, and the chaperonin GroES), consistent with the sporulating, slow-growing Actinomycetota and Bacillota prevalent in soil^52–56^, whereas gut-specific clusters were dominated by toxin–antitoxin systems, single-stranded-DNA-binding proteins, and abortive-infection/anti-defense proteins, consistent with the intense host-phage defense arms race of the gut^57–59^ (Figure 6A).

**Figure 6.**
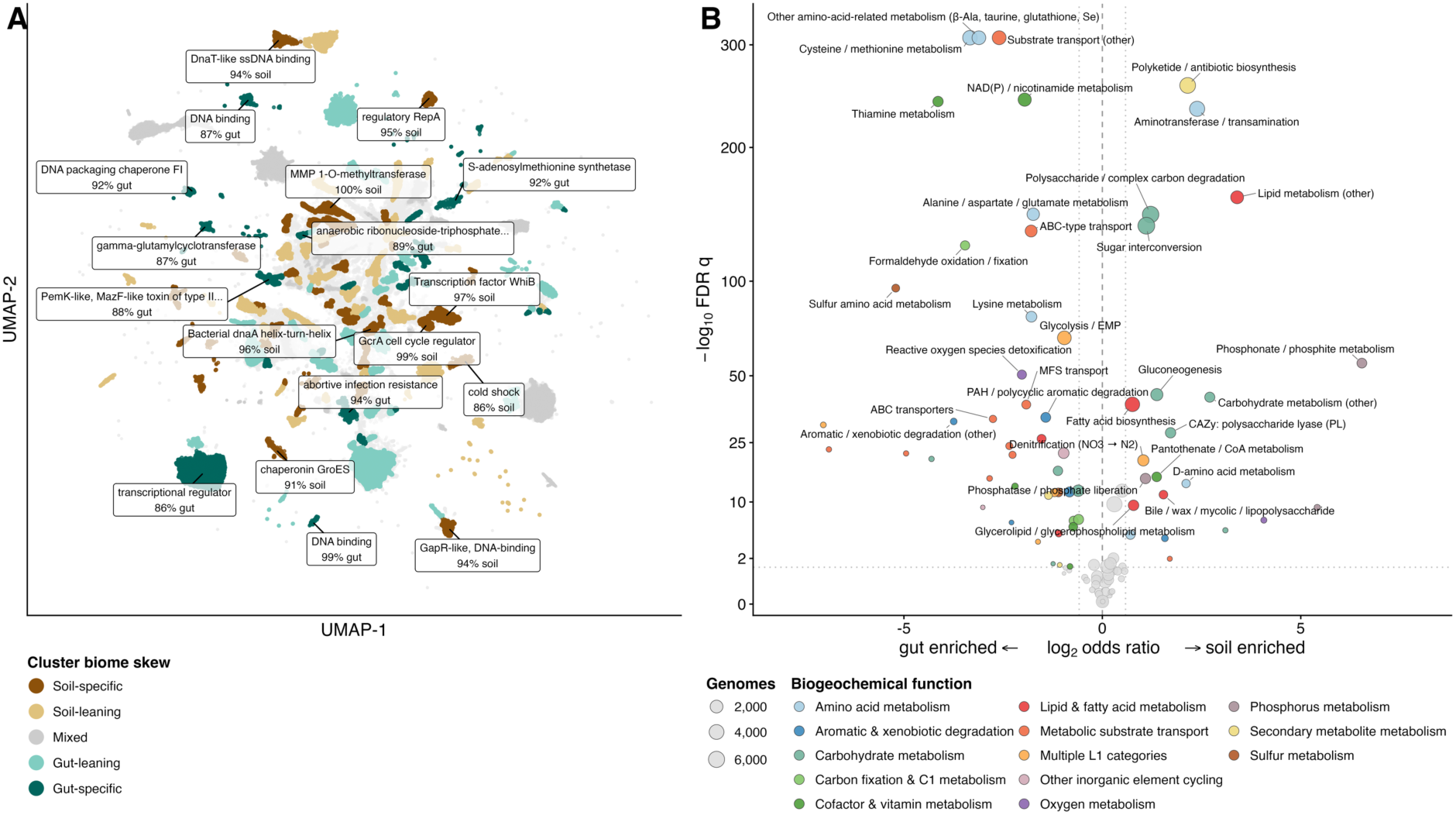
CheckAMG reveals soil- and human gut-enriched auxiliary viral gene functions. **(A)** UMAP of the CheckAMG-PST embeddings calculated for all soil and human gut AVGs identified by CheckAMG. Points are colored by how skewed protein embedding clusters are towards containing proteins from a single biome of origin (see Methods; either soil- or gut-“specific,” “leaning,” or “mixed”). Representative functions of biome-skewed clusters are labeled. **(B)** Auxiliary metabolic functions significantly enriched in soil versus gut viral genomes. Points represent distinct AMG functions either assigned by CheckAMG *annotate* or propagated to sequence-similarity invisible AVGs identified by CheckAMG *de-novo*. Sizes of points are proportional to the number of viral genomes encoding the function and are colored by the L1 biogeochemical category the function is assigned to in the CheckAMG reference. Points to the left and right of the thin dashed lines represent AMG functions enriched in either human gut (left) or soil (right) environments (*q* < 0.05, Cochran-Mantel-Haenszel test stratified by total gene count, Benjamini-Hochberg correction), in the context that they were analyzed in this study. For simplicity, only the 16 enriched functions with the lowest *q* in each biome are labeled. Gray points correspond to non-enriched functions.

We also examined which metabolic functions were biome-enriched and identified 22 soil-enriched and 42 gut-enriched functions, while 34 functions showed no biome specific difference (Cochran-Mantel-Haenszel test stratified by total gene count, *q* < 0.05 Benjamini-Hochberg corrected, odds ratio > 1.5 or < 0.667; Supplemental Table 14). Enriched functions largely tracked the known biochemistry of each environment (Figure 6B; full results in Supplemental Results). Soil-enriched AMGs included phosphonate and polyphosphate metabolism, carbohydrate-active enzymes for complex-polysaccharide degradation, polyketide synthesis, and denitrification, consistent with soil nutrient cycling, the breakdown of plant-derived organic matter, and secondary metabolism prevalent in soils^60–63^. Gut-enriched AMGs tracked the gut nutritional environment, including thiamine, sulfur-amino-acid, formaldehyde, and alanine/aspartate/glutamate metabolism. At the level of predicted host phylum, associations again matched established biology, such as iron reduction in soil Desulfobacterota_B^64^ and organosulfur metabolism in human gut Bacillota^65,66^ (Extended Data Figure 10). Because auxiliary functions are expected to reflect the metabolism of their hosts and environments, this concordance provides independent support that the AVGs CheckAMG identifies are biologically coherent rather than artifacts of annotation.

## Discussion

CheckAMG resolves two separate limitations that have constrained the study of virus-encoded auxiliary genes. By combining a genome-context classifier of per-protein viral origin with an auxiliary-function-specific weighting scheme, CheckAMG distinguishes AVGs from both non-viral contamination and core viral function using criteria that are explicit, tool-independent, and can be applied after the fact to any candidate annotation. By pairing this interpretable annotation-based path with CheckAMG-PST, a finetuned protein-genome language model, CheckAMG additionally identifies auxiliary genes that share no detectable sequence homology with any reference database, a class of AVG that homology-based tools cannot access in principle rather than merely in practice.

The scale of disagreement we observed between CheckAMG, DRAM-V, and VIBRANT explains why reported AMG catalogs have varied so widely across previous studies. Most disagreement did not reflect differences in the ability to detect sequence homology. VIBRANT’s curated AMG reference simply lacks many identifiers that CheckAMG treats as high-confidence AMGs, so no adjustment of its stringency could recover them without updating VIBRANT’s reference AMG database. Unlike CheckAMG and DRAM-V, VIBRANT also does not perform any curation of AMG functional annotations. DRAM-V’s reference AMG database is still limited compared to CheckAMG, and its default setting pools auxiliary scores 1 to 3, which apply different rule-based standards of viral evidence, so most of its calls are on sequences that the external tool used for evaluation, geNomad^39^, does not classify as viral. These are differences in curation and reference composition, not in detection sensitivity, and they mean AMG counts reported by different tools are not directly comparable even when applied to identical data.

The AMG weight framework introduced here, which scores any candidate annotation on a continuous scale from its metabolic specificity and its rarity in viral genomes, offers a quantitative way to evaluate AMG calls from each tool, although CheckAMG also applies it as its own default filter. It can therefore be applied retrospectively to reassess AMG catalogs already reported in the literature. Two caveats attach to this. The framework was developed for CheckAMG and neither DRAM-V nor VIBRANT carries a comparable score, so comparing tools on AMG weight asks how each tool’s calls rate under CheckAMG’s definition of AMG-likeness rather than under its own. It is computed from the annotation rather than from the tool that produced it, which is what allows it to be applied to any tool’s output, and the cautious-function set used above offers a separate comparison that does not depend on it. Furthermore, because it is designed to be updated as new studies reveal, characterize, and experimentally validate AVGs, both the AMG weight framework and CheckAMG can continue to evolve alongside community-established guidelines for studying AVGs.

Applied at scale, CheckAMG recovered auxiliary functions that track the known biochemistry of each biome studied. Soil-enriched functions included phosphonate and polyphosphate metabolism, carbohydrate-active enzymes for complex-polysaccharide degradation, polyketide synthesis, and denitrification, consistent with nutrient cycling and the breakdown of plant-derived organic matter in soils, while gut-enriched functions tracked the gut nutritional environment, including thiamine, sulfur-amino-acid, and one-carbon metabolism. At the host level, predicted associations such as iron reduction in Desulfobacterota_B and complex carbon degradation in Thermoplasmatota similarly matched established biology. That computationally predicted AVGs recapitulate independently known host and environmental biology, rather than appearing as annotation artifacts, is itself a form of validation that is difficult to obtain any other way for genes whose auxiliary role is inferred rather than experimentally confirmed. This concordance also extends the reach of AVG ecology beyond the handful of biomes and gene families characterized in detail so far, complementing recent surveys of AMGs across the global ocean^21^ and across specific biogeochemical functions such as methane^67,68^, sulfur^68–71^, and soil carbon cycling^61,72,73^, by placing soil and human-gut viromes on the same curated, quantitative footing for the first time.

That our deep-learning classifier in the CheckAMG *de-novo* module (CheckAMG-PST) recovered auxiliary genes entirely without any annotation support (including non-auxiliary or general viral functions), at a scale exceeding 100,000 proteins, shows how much AVG diversity is invisible to homology search even when that search is exhaustive. Cross-referencing this set against defense-system profiles^51^ and predicted protein structure recovered a putative function for 78,695 of the 109,107 sequence-similarity invisible genes (72.1%), while the remaining 30,412 (27.9%) could not be explained by any external evidence and instead fell into 106 embedding-based protein families, 56 of which contained no member of known function, consistent with genuinely uncharacterized auxiliary function rather than misclassification. This pattern mirrors a broader trend in which protein and genome language models are beginning to recover functional signal that sequence homology alone misses, as recently shown for anti-phage defense systems^32^, structure-informed phage gene annotation^74^, and genome-context-aware protein annotation more generally^75^. CheckAMG extends this approach to a harder, compound problem, in which a gene must first be established as both virus-encoded and auxiliary before its function is even considered, and suggests that language-model-based discovery will be similarly productive for the physiological and regulatory categories of viral dark matter that CheckAMG also identifies.

Several limitations qualify these conclusions. The viral-origin labels used throughout the benchmark come from geNomad, and CheckAMG’s own classifier was trained in part on geNomad’s training data, so the two are not fully independent. Moreover, functional labels propagated to sequence-similarity invisible AVGs from CheckAMG-PST embedding space are reliable at the level of broad auxiliary category but considerably less so for exact function names, so specific functional claims for individual sequence-similarity invisible genes should be read as descriptive rather than confirmed. However, the unmodified ESM-2 backbone transferred labels more accurately than CheckAMG-PST. CheckAMG-PST’s contribution therefore lies in the AVG call itself rather than in label transfer. Moreover, AMG weight scoring, which underlies both the default curation stringency and the cross-tool comparison, is not implemented for APGs and AReGs, reflecting the comparative novelty of these categories and the absence of equivalently rich ontological resources for physiological and regulatory function, a gap that future versions of CheckAMG and the wider field will need to address. CheckAMG-PST’s embedding space also inherits whatever biases are present in its PST backbone (including underlying ESM-2 models that PST builds off of), the PST pretraining corpus, and the labeled training proteins used for fine-tuning, so its performance on lineages, functions, or environments poorly represented in any of these stages remains untested. Lastly, our large-scale application was limited to uncultivated soil and human-gut viral genomes from a single (albeit large) database. While the biome-specific patterns recovered here are internally consistent and ecologically plausible, extending this analysis to viruses of marine, freshwater, and other host-associated systems and more comprehensive databases will be needed to establish how general these patterns are.

Overall, CheckAMG turns AVG identification from a largely qualitative judgment into a quantitative framework. CheckAMG *de-novo*’s finetuned protein-based genome language model further extends that framework to genes that no homology-based method, including CheckAMG’s own *annotate* module, can reach. As a result, CheckAMG enables the discovery of auxiliary protein families with no functionally labeled counterpart in existing databases. CheckAMG thus provides a reproducible, scalable framework for identifying the genes through which viruses reprogram host biology.

## Methods

### CheckAMG *annotate* pipeline, databases, model training, and evaluation

#### Sequence filtering, ORF prediction, and translation

The annotation module of CheckAMG is implemented as a Snakemake workflow with a command-line interface that can accept input sequences in either nucleotide or amino-acid FASTA format. For nucleotide inputs, sequences (scaffolds) are checked for indicators of circularity by checking for direct terminal repeats (DTRs), inverted terminal repeats (ITRs), and circularly permeated sequence, using a *k*-mer matching approach as employed by the tool CheckV^22^. Scaffolds are then filtered by length to match the user-specified minimum, or the default 5,000 bp (Figure 1). Scaffolds passing the length filter undergo open reading frame (ORF) prediction with pyrodigal^76,77^, as implemented by the python package metapyrodigal (github.com/cody-mar10/metapyrodigal) for efficient parallelization, with metagenomic models enabled as well as additional virus models (from prodigal-GV^39^) to select the optimal genetic code for ORFs when translating into amino acid sequences (proteins). The proteins resulting from ORF prediction, or the input user proteins (for amino-acid inputs) are then filtered so that all scaffolds encode a user-specific minimum number of ORFs, or the default minimum of 4 (Figure 1). All proteins encoded by a scaffold that does not meet the minimum number of proteins are discarded from further analysis.

#### Functional annotation of query sequences

Filtered protein sequences are then annotated for putative functions by searching for sequence homology to reference profile hidden Markov models (HMMs)^78^ (Figure 1). As of CheckAMG v1.1, the reference profile HMM databases include KEGG KOfam^36^ version 2025-12-01, Functional Ontology Assignments for Metagenomes (FOAM)^43^ version 2023-01-10, Prokaryotic Virus Remote Homologous Groups database (PHROGs)^38^ v4, Pfam-A^37^ v38, CAZymes from dbCAN^42^ v14, Curated Annotations for Microbial (Poly)phenol Enzymes and Reactions (CAMPER)^45^ v1.0.0, and the custom set of prokaryotic metabolism HMMs from METABOLIC^44^ v4.0. These databases were chosen to balance curated functions from confident references (KOfam, Pfam-A) with increased sensitivity for functions encoded by environmental and uncultivated microbes and their viruses (FOAM, PHROG), as well as additional metabolic functions that can be overlooked by the other databases (dbCAN, CAMPER, METABOLIC). Since PHROGs do not provide HMM-formatted profiles of protein families, profile HMMs were built from multiple sequence alignment FASTA files of PHROG protein families using the “Builder” function from the pyhmmer Plan7 python library^79^ with default settings. The HMM-formatted PHROGs are included in the CheckAMG database.

To ensure that functions are not being inferred from profiles representing small individual domains rather than full proteins, all profiles are searched unconditionally, but hits to profiles whose database-defined threshold falls below 60 bits (default, but configurable by users) are excluded from functional annotation during per-hit assessment (using full-sequence thresholds when available for KEGG, FOAM, CAMPER, and METABOLIC profiles, domain-level thresholds otherwise, and the gathering threshold for Pfam). Profile HMMs without database-defined thresholds (PHROGs, dbCAN, and a portion of profiles in the other databases) are not subject to this exclusion (Supplemental Figure 1).

Protein sequences are searched for homology to the referenced filtered profiles with HMMsearch^78^, as implemented by the python package pyhmmer^79^. A broad HMMsearch is conducted for all query proteins against each filtered reference database with an initial, permissive maximum E-value of 1e-2. The resulting hits are then filtered in a three-stage process (Supplemental Figure 1). In the first stage, hits are filtered so that all hits have a bit-score of at least 60, an alignment coverage of at least 40% of the query sequence, and the same minimum coverage for the aligned profile HMM. In the second stage, the bitscores of the hits passing the first stage are compared to the database-defined bitscore thresholds for detection for the profile HMM in the hit (gathering threshold for Pfam). Hits with a bitscores greater than or equal to the database-defined threshold are accepted. Hits with bitscores below the database-defined threshold are subject to the third filtering stage, except for Pfam profiles that publish a gathering cutoff, which is enforced without relaxation. The third stage is a database-specific, adaptive strategy, inspired by the “anvi-run-kegg-kofams” method employed by Anvi’o^80^ to improve sensitivity for divergent sequences encoded by uncultivated microorganisms while retaining precision^81^. In this stage, hits that have a maximum E-value of 1e-5 and a bitscore that is at least 50% of the database-defined threshold are accepted, while hits with an E-value greater than 1e-5 or a bitscore less than 50% of the database-defined threshold are discarded. Profiles that publish only a domain-level threshold are evaluated per aligned domain against that threshold, each domain subject to its own coverage requirement and to the same two-part rule, and the hit is retained when any single domain passes. On this domain path neither the full-hit coverage test nor the 60-bit floor applies, so a retained hit can score below 60 bits. Hits to profiles without database-defined bitscore thresholds are not subject to filtering by the second- and third-stage (Supplemental Figure 1).

Therefore, all HMMsearch hits passing these filters have a bitscore of at least 60, an alignment coverage of at least 40% of the query sequence, and the same minimum coverage for the aligned profile HMM, and a maximum E-value of 1e-2 (for hits to profiles without database-defined thresholds), and hits to profiles with database-defined thresholds either have a bitscore that meets or surpasses that threshold and a maximum E-value of 1e-2, or a bitscore that is at least 50% of that threshold and a maximum E-value of 1e-5. Since hits to profiles with thresholds below 60 bits are excluded from annotation, all hits to profiles with thresholds that reach the annotation path therefore have a minimum bitscore of 30 when the third filtering stage is applied.

The HMMsearch results feed two independent downstream paths (Supplemental Figure 1). For functional annotation, only hits passing the quality filters above are retained, and the single best hit per reference database per protein is selected, as done by the metagenome annotation pipeline Metacerberus^82^; the lowest E-value is used as the primary tiebreaker, followed by the highest bitscore. In parallel, a separate V-score table is produced from all hits at E-value ≤ 1e-5, regardless of other hit metrics, retaining the best hit per protein–database combination independently across databases. The V-score table is used exclusively for V- and V_L_-score assignment for downstream viral origin confidence predictions and is not used for functional annotation.

#### Candidate AVG reference annotations

The functional annotation curation module of CheckAMG *annotate* addresses two complementary problems: (1) determining whether a given HMM-derived functional annotation is likely to represent a metabolic, physiological, or regulatory function rather than a non-specific or broadly-functional domain, and (2) determining whether that function is likely to be auxiliary (i.e., rarely found in viral genomes under normal circumstances) rather than a conserved viral core function. To accomplish this, CheckAMG constructs a suite of curated reference tables prior to analysis, using ontological classifications and keyword-matching strategies applied across all reference HMM databases. These tables define candidate sets for each AVG category (AMG, APG, AReG), quantify the metabolic specificity of AMG annotations, and flag annotations associated with known non-auxiliary viral functions. The construction of all reference tables is fully documented and publicly available on GitHub for reproducibility (github.com/AnantharamanLab/CheckAMG/blob/main).

An AMG candidate table was assembled from multiple database sources. From KEGG, all KOs classified under the “Metabolism” A-level BRITE hierarchy category, as well as those under the “Unclassified: metabolism” B-level category and relevant “Brite Hierarchies” B-level sub-categories, were selected as initial candidates. However, several B-level categories were excluded regardless of their A-level classification, including “Protein families: genetic information processing,” “Signal transduction,” “Information processing in viruses,” “Viral protein families,” “RNA family,” “Translation,” “Replication and repair,” “Nucleotide metabolism,” “Transcription,” “Chromosome,” and “Glycan biosynthesis and metabolism.” Additional C-level pathway categories that describe predominantly non-metabolic or structurally specific functions (e.g., “Transcription factors,” “Secretion system,” “Glycosyltransferases,” “Peptidoglycan biosynthesis,” “Bacterial motility proteins,” “Two-component system”) were also excluded; a complete list is provided in Supplemental Table 15. An additional filtering step was applied to KOs in the “Enzymes with EC numbers” C-level category only, using keyword matching against annotation descriptions to remove profiles indicative of non-metabolic functions (e.g., “lysozyme,” “glycosyltransferase,” “capsid,” “DNA polymerase,” “replication;” see Supplemental Table 16 for the complete keyword list). From FOAM, all HMMs not classified under the L1 category “21_Cellular response to stress” or the L2 category “Polysaccharide biosynthesis and assembly of cell surface structure” were included. All HMMs from the METABOLIC and CAMPER databases were included as AMG candidates, as both were designed to catalog metabolically-relevant protein families. All CAZyme HMMs from dbCAN were also included, reflecting the relevance of carbohydrate-active enzymes to host carbohydrate metabolism and nutrient cycling.

Additional Pfam-based AMG candidates were identified by mapping Pfam HMMs to Gene Ontology (GO) terms^46^ using pfam2go annotations (https://current.geneontology.org/ontology/external2go/pfam2go)^83^. A curated set of metabolic-like GO terms was constructed by taking all descendants of the biological process (BP) root term GO:0008152 (metabolic process), then subtracting descendants of subtrees that capture predominantly non-metabolic processes, including macromolecule metabolic process (GO:0043170), DNA metabolic process (GO:0006259), macromolecule biosynthetic process (GO:0009059), nucleobase-containing compound metabolic process (GO:0006139), lipopolysaccharide metabolic process (GO:0008653), tetrahydrofolate metabolic process (GO:0046653), methylation (GO:0032259), and viral process (GO:0016032). A small number of terms within excluded subtrees that are clearly relevant to metabolism (e.g., glycolytic process [GO:0006096], polysaccharide metabolic process [GO:0005976], and coenzyme A metabolic process [GO:0015936]) were rescued and retained. Selected molecular function (MF) subtrees were also included, covering major enzyme classes (oxidoreductase, transferase, hydrolase, lyase, isomerase, and ligase activities), transmembrane transporter activity, and relevant binding activities, with specific MF subtrees excluded (e.g., nucleic acid binding [GO:0003676], nucleotide binding [GO:0000166], glycosyltransferase activity [GO:0016757], and hydrolase activity acting on glycosyl bonds [GO:0016798]). Complete lists of included and excluded GO subtrees, as well as all rescued terms, are provided in Supplemental Table 17.

Pfam HMMs mapped to at least one GO term in the curated metabolic set were then identified as AMG candidates. Pfam HMMs whose names contained the word “domain” were excluded from this GO-based set to avoid including profiles that represent small individual protein domains rather than complete functional units. The same keyword-matching approach used to filter KEGG enzyme annotations was applied a second time to remove Pfam candidates with descriptions suggestive of non-metabolic functions despite metabolic GO mappings. Finally, all Pfam HMMs present in the DRAM-V AMG database^27^ were included as AMG candidates. The final AMG candidate table was formed by taking the union of all candidates from these sources.

APG and AReG candidate tables were constructed using keyword-matching against the combined HMM description table. For APG candidates, all HMMs with descriptions matching a set of physiology-related terms were selected using case-insensitive substring or whole-word matching. Keywords covered functional categories including heat shock and cold shock responses, oxidative stress, sporulation, stringent response, biofilm formation, motility and flagellar assembly, virulence, toxin-antitoxin systems, lysogeny, bacterial secretion systems, and anti-defense/anti-CRISPR systems. A small set of case-sensitive terms (e.g., “Acr” for anti-CRISPR proteins, “Abi” for abortive infection systems) were also matched using whole-word exact case matching. For AReG candidates, a similar approach was used with regulatory keywords, including “regulation,” “regulator,” “repressor,” “activator,” “transcription,” “sigma,” “two-component,” “quorum,” “autoinducer,” “DNA-binding,” “RNA-binding,” “helix-turn-helix,” “nucleoid-associated,” “antitermination,” and related terms. Complete keyword lists for both APG and AReG tables are provided in Supplemental Table 6.

Because the three AVG categories are not mutually exclusive in terms of keyword coverage, overlaps between the AMG, APG, and AReG candidate tables were identified and resolved to ensure each HMM profile was assigned to at most one category. All HMMs appearing in both the APG and AReG candidate tables were retained in the AReG table only, as upon inspection these annotations were more consistent with regulatory rather than physiological functions. HMMs appearing in both the AReG and AMG candidate tables were similarly retained in the AReG table in most cases, with the exception of a small number of annotations describing proteins that serve primarily as regulatory subunits or accessory components of metabolic enzyme complexes (e.g., methane monooxygenase regulatory protein B [MmoB], photosystem II oxygen-evolving enhancer proteins [PsbO, PsbP, PsbQ], ATP phosphoribosyltransferase regulatory subunit [HisZ], and two-component flavin-dependent monooxygenases), which were retained in the AMG table. HMMs appearing in both the AMG and APG candidate tables were retained in the APG table in most cases, with the exception of one profile (K04035). After resolving all overlaps, the AMG, APG, and AReG tables each contain mutually exclusive sets of HMM profiles. The final AMG, APG, and AReG tables are provided in Supplemental Tables 3, 4, and 5 respectively.

#### Calculation of metabolic ratios and AMG weights

To quantify how metabolic-like each AMG candidate annotation is, a metabolic ratio was calculated for each profile in the AMG table. The metabolic ratio is defined as the proportion of a profile’s known ontological pathway classifications that are considered metabolic, and ranges from 0 to 1. The denominator is the total number of unique ontological pathway categories a profile is mapped to; the numerator is the number of those categories that are classified as metabolic. This design means that profiles associated exclusively with metabolic processes receive a ratio of 1, while profiles that are also classified under non-metabolic pathways receive lower values. The specific ontological source used to calculate the metabolic ratio differed by database, and where multiple sources provided a ratio for the same HMM, the most conservative (lowest) value was retained.

For KEGG KOs, the metabolic ratio was calculated from the KEGG BRITE hierarchy. All C-level (pathway/sub-category) entries were classified as metabolic or non-metabolic based on their B-level parent category and C-level name, using the same exclusion criteria applied during AMG table construction (see above and Supplemental Table 18). The metabolic ratio for each KO was then defined as the number of unique metabolic C-level entries divided by the total number of unique C-level entries in which that KO appears. For KOs in the “Transporters” functional category, an additional sub-classification was applied to distinguish transporters of metabolically relevant substrates (e.g., specific ions, amino acids, organic acids, and small metabolites; see Supplemental Table 19) from transporters of non-specific or structural substrates; only the former were counted in the metabolic numerator.

For FOAM HMMs, because each FOAM profile can map to one or more KEGG KOs (via KO identifiers encoded in the FOAM HMM headers), metabolic ratios were first calculated at the level of individual KOs using the FOAM L1/L2 ontological hierarchy, applying the same metabolic/non-metabolic classification logic described above (with “07_Nucleic acid metabolism” and “21_Cellular response to stress” excluded from L1, and “Glycosyltransferase,” “Glycoside hydrolase,” and “Polysaccharide biosynthesis and assembly of cell surface structure” excluded from L2, along with transporter sub-classification at L1). The metabolic ratio for each KO was the number of unique metabolic L2 categories divided by the total. FOAM HMM-level ratios were then derived by summing the metabolic and total path counts across all KOs mapped to that profile before dividing, such that multi-KO FOAM profiles integrate evidence from all associated functions.

For Pfam HMMs, metabolic ratios were calculated from GO annotations via pfam2go. The metabolic ratio for each Pfam was defined as the count of its GO annotations falling within the curated metabolic GO set (described above) divided by the total count of in-scope GO annotations, where “in-scope” excludes very broad GO terms describing general cellular anatomy (specifically, descendants of “membrane” [GO:0016020] and “cellular anatomical structure” [GO:0110165]) that would otherwise inflate the denominator without contributing meaningful functional information. For dbCAN (CAZyme) HMMs, metabolic ratios were calculated analogously by mapping EC numbers extracted from CAZyme descriptions to GO terms via ec2go, then applying the same metabolic GO set and in-scope denominator. For TIGRFAM HMMs present in the METABOLIC database, metabolic ratios were computed using GO annotations from the NCBI PGAP^84^ HMM table (ftp.ncbi.nlm.nih.gov/hmm/current/hmm_PGAP.tsv) via the same GO-based approach. METABOLIC and CAMPER HMMs not scored by any of the above routes (i.e., those not assignable to KEGG KOs and lacking GO annotations) were assigned a metabolic ratio of 1.0, reflecting the design intent of both databases to include only metabolically-relevant protein families. When multiple scoring routes provided a metabolic ratio for the same HMM, the lowest (most conservative) value was retained. Only profiles with at least one metabolic pathway mapping (metabolic ratio > 0) were retained in the scored AMG table.

To additionally account for how commonly a given function is encoded by viruses, and therefore how likely it is to represent a truly auxiliary rather than a conserved viral function, V_L_-scores from the V-Score database^33^ were incorporated into the scoring. V_L_-scores (the log_10_-transformed number of viral hits for a given protein family) are available in the V-Score database for KEGG KOs and Pfam HMMs. Because V_L_-scores are not provided directly for FOAM profiles, they were propagated by averaging the V_L_-scores of all KEGG KOs mapped to each FOAM profile via its KO assignments; FOAM profiles with at least one KO V_L_-score received a propagated V_L_-score in this way. Profiles from METABOLIC, CAMPER, and dbCAN that could not be mapped to KEGG KOs with V_L_-scores did not receive a V_L_-score and were not adjusted.

The final AMG weight for each profile was then calculated as:

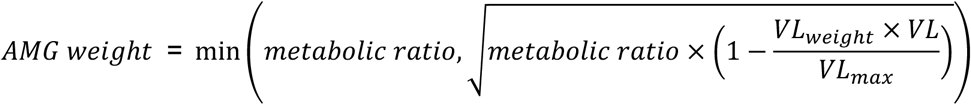

Where *VL* is the V_L_-score assigned to the HMM profile and *VL_max_* is the maximum V_L_-score observed across all profiles in the AMG table (4.63). The term 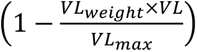 acts as a penalty, with *VL_weigℎt_* (0.75) controlling the weight of the penalty. Profiles with low V_L_-scores (rarely found in viruses) receive a penalty near zero and their AMG weight approaches the metabolic ratio. Profiles with high V_L_-scores (commonly found in viruses) receive a strong penalty and their AMG weight is substantially reduced. To obtain the final AMG weight, the geometric mean of the metabolic ratio and this penalty is taken, then capping the result at the metabolic ratio, to ensure that the V_L_-score adjustment can only decrease the AMG weight and never increase it. For profiles without a V_L_-score, the AMG weight is set equal to the metabolic ratio. The AMG weight is thus a single score ranging from 0 to 1 that integrates both the metabolic specificity and the viral rarity of an annotated function: values near 1 indicate that a function is predominantly metabolic and rarely encoded by viruses (consistent with a genuine AMG), while values near 0 indicate a function that is either predominantly non-metabolic, highly common in viral genomes, or both (Figure 2; Supplemental Table 3).

Metabolic ratios and AMG weights are calculated only for profiles in the AMG table, and not for APG or AReG profiles in CheckAMG v1.1. This is due to the relative novelty of APGs and AReGs as distinct AVG categories compared to AMGs, as well as the more complex and context-dependent nature of physiological and regulatory functions, which do not map as cleanly to existing metabolic ontologies. This limitation can be addressed in future versions of CheckAMG as the field develops richer ontological resources for these functional categories.

#### Filtering non-auxiliary functions from AVGs

Separate filter tables were constructed for each AVG type to catalog HMM profiles whose annotations are indicative of known essential or non-auxiliary viral functions. For AMGs, the AMG filter table organizes profiles into five categories using a combination of database ontological classifications and keyword matching: (1) “essential” viral functions, including structural proteins, DNA replication machinery, lysis enzymes, and cell entry/adhesion proteins; (2) nucleotide metabolism (identified from the KEGG “Nucleotide metabolism” B-level and related C-level categories, the FOAM “07_Nucleic acid metabolism” L1 category, PHROG profiles classified as “DNA, RNA and nucleotide metabolism,” and a set of nucleotide-related keywords); (3) glycan biosynthesis and modification (glycosyltransferase [GT] and glycoside hydrolase [GH] CAZyme families from dbCAN, FOAM profiles in the “Glycosyltransferase,” “Glycoside hydrolase,” and “Polysaccharide biosynthesis and assembly of cell surface structure” L2 categories, KEGG KOs in the “Glycan biosynthesis and metabolism” B-level and “Glycosyltransferases” C-level categories, and matching keywords); (4) lipopolysaccharide and phospholipid metabolism (KEGG KOs in the “Lipopolysaccharide biosynthesis,” “Glycerophospholipid metabolism,” and “Ether lipid metabolism” C-level pathways, and matching keywords); and (5) methylation-related functions (keyword-matched). For each category, HMMs from other databases with descriptions matching those from KEGG, FOAM, or PHROG reference profiles were identified via normalized description matching (case- and punctuation-insensitive, after stripping EC numbers and gene abbreviations) and added to the respective filter category. By default, only the “essential” filter category is applied as a hard removal step; the remaining four categories (nucleotide metabolism, glycan, lipid, methylation) are recorded for each prediction but are not applied unless the user activates them via filter preset options (e.g., “filter_glucan,” “filter_nucleotide”), enabling more stringent curation when warranted by the biology of the input dataset.

For APGs, a filter table was constructed using only the “essential” viral function category (structural, packaging, and surface proteins: capsid, portal, tail, head, sheath, terminase, and immunoglobulin-related terms), as the other AMG filter categories (nucleotide metabolism, glycan, lipid) are less applicable to physiological gene curation. For AReGs, the “essential” filter additionally excluded DNA replication, nuclease activity, recombination, transposition, and methyltransferase-related functions that are not directly involved in gene regulation. Importantly, annotations that also matched regulatory terms (“regulation,” “regulatory,” or “regulator”) were rescued from the AReG essential filter, to prevent the inadvertent removal of genuine regulatory genes with replication-adjacent descriptions. A full list of AMGs, APGs, and AReG filters are provided in Supplemental Table 7.

Using these reference tables, CheckAMG *annotate* applies a multi-step curation pipeline to all AVG predictions (Supplemental Figure 1), following established best practices for AVG identification in viral ecology^1,26^. All parameters are configurable by the user; the following describe default behavior. The pipeline operates in four sequential stages applied separately to AMG, APG, and AReG candidates.

In the first stage (AMGs only), AMG candidates are pre-filtered by AMG weight. The AMG reference table is subset to retain only profiles whose AMG weight meets or exceeds the minimum threshold (default: 0.6), producing a set of “weight-passing” AMG IDs. A gene is then classified as an AMG candidate only if at least one of its HMM annotations matches a weight-passing AMG profile.

In the second stage (all AVG types), “AVG array” detection is performed on the full annotation table before any keyword-based filtering is applied, so that array membership is determined using the same candidate pool across all three AVG types. A gene is considered part of an AVG array if it belongs to a contiguous run of consecutive genes (within the same scaffold) each annotated as any AVG type (AMG, APG, or AReG). By default, at least three consecutive AVGs in a row are required to be flagged as an AVG array, and all AVGs in a flagged array are excluded from final AVG predictions (though this minimum array length is configurable), as contiguous runs of AVG-like annotations are more consistent with host genomic contamination^27^, MGE cargo gene clusters^21^, or misclassified stretches of viral genes^22^ than with genuine auxiliary functions.

In the third stage (all AVG types), keyword/flag-based filtering is applied independently to each AVG type using the respective filter table. For each AVG type, a gene is first confirmed as a candidate by requiring at least one HMM annotation (among the database top hits) matching a profile in the corresponding AVG reference table (the weight-passing AMG table, the APG table, or the AReG table). Each candidate is then checked against the filter table: by default, if any of its HMM annotations (database top hits) match a profile flagged in the “essential” category for that AVG type, the gene is removed. The remaining filter categories (nucleotide metabolism, glycan, lipid, and methylation for AMGs; see above) are not applied by default but can be activated via user-specified presets for extra-strict curation.

Two additional reference annotation tables were constructed for feature engineering (described below) and for user reference. A viral hallmark gene table was built by selecting all HMMs from the combined HMM description table whose annotation descriptions contained the terms “virion structur*,” “capsid,” “portal,” “tail,” or “terminase” (case-insensitive), following a previously established framework for viral hallmark gene definition^85^. A mobility/MGE gene table was constructed by filtering the V-Score database for profiles whose descriptions matched terms associated with mobile genetic elements, including transposases, recombinases, resolvases, relaxases, DDE-domain proteins, plasmid partitioning and stability genes, and bacterial conjugation transfer proteins (Tra*) (Supplemental Table 20), supplemented by plasmid marker genes from the geNomad database^39^.

#### Training and test dataset construction

A ground-truth dataset for downstream models and classifiers was assembled from two complementary sources. The first source was the training and benchmarking dataset utilized by geNomad^39^, which comprises artificially fragmented sequences derived from curated bacterial, archaeal, eukaryotic, plasmid, and viral genomes (chromosomal sequences 2–637 kb, plasmid sequences 2–607 kb, viral sequence lengths 2–422 kb), supplemented with mock provirus sequences (5–30 kb) constructed by embedding phage genomes into prokaryotic chromosomes at defined positions. Sequences included in the geNomad authors’ quality-control exclusion list were removed. Additionally, fragmented sequences whose parent genomes overlap with the phage genomes used to construct the mock proviruses were excluded to prevent data leakage between the fragmented and mock-provirus subsets. The geNomad dataset thus provides clean, whole-sequence or region-level ground-truth labels: sequences of viral origin (viral genomes or proviral regions within mock proviruses) are labeled viral (True Positive), and all others (bacterial, archaeal, eukaryotic, plasmid sequences, or host-chromosome flanks of mock proviruses) are labeled non-viral (True Negative). Ground-truth labels were assigned consistent with the geNomad authors’ assignments (“sequence_weights.tsv” for non-proviruses and “mock_prophages.tsv” for mock proviruses from zenodo.org/records/8049246).

The second source was a dataset of bacterial and archaeal sequences derived from proGenomes v3^40^ with MGE and prophage boundaries annotated using proMGE^24^. From the proGenomes representative scaffold set, four categories of sequences were prepared: (1) chromosome-only sequences, from which all proMGE-annotated MGE and prophage regions were computationally excised; (2) mixed-chromosome sequences, retaining scaffolds that contain annotated MGE or prophage regions alongside host chromosomal sequence; (3) standalone MGE sequences, consisting only of non-phage MGE regions (transposons and integrative conjugative elements, minimum length 1,000 bp); and (4) standalone virus sequences, consisting only of prophage regions. Ground-truth labels were then assigned to each protein by overlapping its genomic coordinates with the proMGE region definitions (“mges.txt” from promge.embl.de), with additional label modification to ensure overlapping, non-viral regions were not labeled as positives (see *Ambiguous viral region identification and ground-truth label assignments*).

CheckAMG *annotate* v1.1 with database version 1.1 was run on all sequences from both dataset sources (using minimum scaffold length and minimum ORF count of 1 to permit small standalone MGEs). To partition the combined dataset into training and test sets while preventing data leakage, an all-vs-all protein similarity search was first performed using MMseqs2^86^ with parameters “--start-sens 4 --sens-steps 3 -s 7.5 --min-seq-id 0.0” on all proteins from both sources in one combined input. Average amino acid identity (AAI) between each pair of genomes/scaffolds was calculated from the MMseqs2 results using the reciprocal best hits from protein alignments. Pairs of genomes/scaffolds were considered similar if they shared at least 40% AAI and either at least 16 shared proteins or at least 20% of the proteins of both genomes were shared. Similar genomes were then clustered using the Markov Cluster Algorithm (MCL)^87^ with an inflation parameter of 2.0, producing genome clusters in which members share substantial sequence-level similarity.

The dataset was then split into training (90%) and test (10%) sets with three constraints to prevent leakage: (1) all proteins encoded on the same scaffold were moved together as a single unit, so that no scaffold was split across training and test; (2) all scaffolds from the same genome moved together as a unit; and (3) all scaffolds in the same MCL cluster moved together, preventing highly similar sequences from appearing in both training and test. Class balance within the training set was enforced by downsampling the dominant class. This splitting resulted in a training dataset with 13,999,957 proteins encoded by 1,196,497 scaffolds, of which 6,563,453 proteins (46.88%) and 582,564 scaffolds (48.69%) were from host chromosome regions, 1,941,190 proteins (13.87%) and 95,001 scaffolds (7.94%) were from non-viral MGE regions, and 5,495,314 proteins (39.25%) and 544,679 scaffolds (45.52%) were from viral regions (scaffold percentages slightly exceed 100% because some scaffolds contain multiple region types, such as integrated proviruses alongside chromosomal sequence).

Test sets were also constructed in multiple compositions designed to approximate different real-world scenarios, including near-all-virus (88.7% viral, 3.6% MGE, 7.8% host), virus-enriched (70.2% viral, 12.5% MGE, 17.3% host), half viral/host (50.0% viral, 4.6% MGE, 45.4% host), equal viral/non-viral (50.0% viral, 20.8% MGE, 29.2% host), training-distribution (39.3% viral, 13.9% MGE, 46.9% host), equal viral/MGE/host (33.3% viral, 33.3% MGE, 33.4% host), host-enriched (17.0% viral, 12.5% MGE, 70.6% host), near-all-host (5.5% viral, 5.0% MGE, 89.5% host), and MGE-enriched (10.3% viral, 75.0% MGE, 14.7% host). An additional test dataset composed solely of proteins encoded on scaffolds with integrated proviruses was also constructed (57.5% viral, 0.4% MGE, 42.1% host). Overlap between different test compositions was permitted but no scaffold or protein was shared between the training set and any test set. The training dataset and all test datasets are provided as Supplemental Data 1 and 2, and summaries of their protein- and scaffold-level compositions by sequence type and dataset source are provided in Extended Data Figure 2A and its Source Data, and per-dataset classifier performance in Supplemental Table 2.

#### Ambiguous viral region identification and ground-truth label assignments

A challenge in constructing the training dataset was that one of the two source databases intentionally reports MGE boundaries as upper limits that encompass nested MGE types and surrounding host chromosomal sequence^24^. Directly labeling proteins from this dataset as viral without further curation would therefore introduce noisy ground-truth labels. To address this, we implemented an annotation-based viral region identification algorithm in CheckAMG *annotate* that was applied during training data construction to flag proteins in genomically “ambiguous” regions to refine viral/non-viral ground-truth labels in the proGenomes-sourced training and test data. Importantly, this strict algorithm operates exclusively on annotation-derived V- and V_L_-scores and is not applied as a default filter in the CheckAMG *annotate* pipeline for user data; its role is restricted to training data quality control and optional stringent curation for the CheckAMG *annotate* workflow. The algorithm was used solely to relabel proGenomes regions with weak annotation-based support for viral origin as nonviral, rather than for dataset-wide provirus prediction or identification of new viral sequences.

The algorithm identifies putative viral regions on each scaffold in five sequential steps, illustrated step-by-step on an example scaffold in Extended Data Figure 1:

In step 1, seed genes are identified as those with a sliding-window average V_L_-score (calculated across KEGG, Pfam, or PHROG annotations within a fixed nucleotide window; see feature description below) at or above a minimum threshold (default: 3.0). Consecutive seed genes, allowing gaps of up to three non-seed genes, are grouped into candidate viral regions, retaining only those containing at least one seed gene.

In step 2, each candidate region is refined to a core by identifying its leftmost and rightmost genes that pass a V-score gate: a gene passes if it has at least one V-score annotation and all present V-scores meet or exceed the minimum V-score threshold (default: 10.0).

In step 3, boundaries of each refined core are extended outward (“walked back”) while genes remain bridge-eligible (defined as having a window-average V_L_-score within a small margin below the step 1 threshold), subject to a hard limit of 20 genes per side, and requiring that the extended shoulder contain at least one V-score-passing anchor gene.

In step 4, boundaries are optionally snapped outward to nearby strong-V genes (all three V-scores present and all at or above the threshold) within 10 genes, provided the intervening path contains no more than one V-score-failing gene and includes at least one bridge-eligible gene.

In step 5, adjacent refined regions are merged if the gap between them is at most 10 genes and the gap is predominantly bridge-eligible (at least 50% bridge genes, maximum one consecutive non-bridge gene, and at most one V-score-failing gene). Regions with fewer than five genes after refinement are discarded before merging.

A protein is considered to fall outside an ambiguous region only if it is in a final merged region passing all five steps (step5_in_merged_region = True) and is not itself in a genomic position flagged as non-viral by the ground-truth region labels; proteins not meeting both criteria within proMGE-derived scaffolds are excluded from training labels. For proteins in both the geNomad-derived and proMGE-derived sequences, only proteins for which the ambiguous viral region identification algorithm agreed with the proMGE viral label (i.e., step5_in_merged_region = True for strict viral proteins) were retained with a viral label; all others in ambiguous regions were labeled as nonviral. Examples of ambiguous viral regions identified with this algorithm are shown in Extended Data Figure 1.

#### LightGBM model training, evaluation, and implementation in the CheckAMG annotate genome context module

A binary LightGBM classifier^35,88^ (gradient-boosted decision tree ensemble; LightGBM python library v4.6.0) was trained to predict viral (positive) versus non-viral (negative) protein origin. LightGBM was chosen for its scalability to large datasets, its native handling of missing feature values (common when a protein lacks an annotation in one or more databases), and its ability to integrate both local per-protein annotation signals and broader genomic neighborhood context as independent tabular features without requiring nucleotide sequence encoding.

All features used as LightGBM inputs were derived from the annotation-based V-scores and V_L_-scores assigned to each protein by KEGG, Pfam, and PHROG HMM searches conducted by CheckAMG *annotate*, together with scaffold-level and local genomic context computed across all annotated proteins on a scaffold. Features fall into four groups. The first group consists of per-protein annotation scores: the V-score and V_L_-score of the best-matching HMM for each of KEGG, Pfam, and PHROG (6 features). The second group captures scaffold-level and window-level aggregate scores: for each database, the mean V-score and mean V_L_-score of all proteins on the scaffold (6 features), and the mean V_L_-score of proteins within a fixed nucleotide window centered on each gene, calculated independently for KEGG, Pfam, and PHROG (3 features; default window ±5,000 bp, computed only when annotated proteins cover at least 10% of the window length). The third group captures the distance from each protein to the nearest flanking gene with a high viral V-score: for each database, the nucleotide distance to the nearest gene to the left and right whose V-score meets or exceeds the minimum threshold (default: 10.0), yielding left and right distances per database (6 features). The fourth group captures the proximity and annotation character of the nearest flanking MGE-annotated gene: for each database, the nucleotide distance to the nearest left and right gene whose annotation matches the MGE reference table (6 features), plus the V-score and V_L_-score of those nearest left and right MGE genes (12 features). A single additional binary feature indicates whether the scaffold was predicted to be circular (1 feature). Together these 40 features allow the model to make per-protein viral origin assessments that integrate both the local annotation signal of the protein itself and the genomic neighborhood context.

Before conducting a hyperparameter search of the LightGBM model, a stratified 20% hold-out validation split was carved from the training data using scaffold-grouped splitting, ensuring no scaffold was shared between the 80% fit set and the 20% validation set. Per-protein sample weights were assigned to address the imbalance between the three source categories: viral proteins (positive class) received a weight of 1.0, host-derived proteins (negative class) received a weight of 1.0, and MGE-derived proteins (negative class) received a weight of 3.0. This weighting scheme penalizes misclassification of non-viral MGE proteins more heavily than host or viral proteins, reflecting that viruses themselves are MGEs and very diverse, often sharing characteristics with non-viral MGEs.

Hyperparameter optimization was performed on the 80% fit set using Bayesian search (“BayesSearchCV”) from the python package scikit-optimize^89^ v0.10.2 over 40 candidate configurations evaluated with 5-fold “StratifiedGroupKFold” cross-validation (grouped by scaffold to prevent leakage across folds) from the python package scikit-learn^90^ v1.7.2, optimizing for average precision. The search space covered 11 hyperparameters: number of leaves (16-256), maximum tree depth (unconstrained to 12), learning rate (0.01-0.20, log-uniform), number of estimators (300-1,200), row subsampling fraction (0.5-1.0), column subsampling fraction (0.4-1.0), row subsampling frequency (0-10 iterations), L2 regularization (10^-3^-5.0, log-uniform), L1 regularization (10^-3^-2.0, log-uniform), minimum gain to split (0-1.0), and maximum histogram bins (63-255). The best configuration identified (cross-validated average precision = 0.961) was: number of leaves = 256, maximum depth = 12, learning rate = 0.179, number of estimators = 641, row-subsampling fraction = 0.974, column-subsampling fraction = 0.803, row-subsampling frequency = 7, L2 regularization = 0.047, L1 regularization = 2.0, minimum gain to split = 0.730, maximum histogram bins = 254. After hyperparameter selection, the best configuration was used to train a final model on the full 80% fit set, with early stopping monitored on the 20% validation set (patience: 200 rounds, metric: AUCPR). The relative importance of the 40 features considered during training of the final model was calculated using the native feature importance scores provided by the LightGBM python library (Supplemental Table 1).

The continuous probability output of the trained model was converted to three discrete “viral origin confidence” tiers (high, medium, and low) by selecting two probability thresholds. Thresholds were chosen by scanning a grid of candidate values (step size 0.001) and selecting, for each tier, the threshold that simultaneously achieved the precision and recall targets across the greatest number of test datasets. Precision was assessed using the Wilson lower bound (95% CI) to account for uncertainty in smaller datasets; a dataset was considered to pass only if the Wilson lower bound of precision met the target. For the high-confidence tier, targets were a recall ≥ 0.25 and Wilson lower-bound precision ≥ 0.95, corresponding to a False-Discovery Rate (FDR) of < 5%; for the medium-confidence tier, Wilson lower-bound precision ≥ 0.90 (FDR < 10%) and recall ≥ 0.50. These precision/FDR cutoffs follow the precedent for high-confidence and slightly more permissive viral classification established by the virus identification software, geNomad^39^. Additionally, a hard per-dataset precision floor of 0.95 (high) and 0.90 (medium) was enforced on the integrated-provirus test dataset (“test_provirus”): candidate thresholds that failed to meet this floor on that dataset were ineligible for selection regardless of their behavior on other datasets. This constraint ensures that the advertised confidence tiers hold on the most challenging compositional case the classifier encounters in practice. Among candidate thresholds passing the same number of datasets, the one achieving the highest mean recall across passing datasets was preferred; ties were broken by selecting the lower threshold. The final selected thresholds were 0.910 for high confidence and 0.454 for medium confidence, with the low tier encompassing all predictions (threshold 0.000).

The final model was evaluated on all test datasets using AUPRC, AUROC, precision, recall, False-Discovery Rate (FDR), F1 score, Matthews correlation coefficient (MCC), and balanced accuracy, reported separately for each of the three confidence tiers. Probability calibration was assessed using expected calibration error (ECE) with 15 equal-width bins. Performance was summarized across the full range of test dataset compositions described above, allowing assessment of model behavior from near-homogeneous viral inputs to near-homogeneous host or MGE inputs and integrated-provirus scenarios.

In the CheckAMG *annotate* pipeline, the trained LightGBM model is applied to every input protein after HMM annotation and feature computation (Supplemental Figure 1). Each protein receives a continuous viral-origin probability and an associated confidence tier (high, medium, or low), which is propagated to downstream AVG curation. The continuous probability is also reported per protein in the pipeline output, enabling users to apply alternative thresholds when the biology of their input dataset warrants it.

### Comparing CheckAMG *annotate*, DRAM-V, and VIBRANT

AMG detection by CheckAMG was benchmarked against two existing tools, DRAM-V^27^ and VIBRANT^28^, representing the most widely adopted automated AMG identification approaches in the field. Since CheckAMG identifies APGs and AReGs in addition to AMGs, only AMG predictions were considered for these comparisons. The comparison addressed two questions: which functions each tool is able to annotate as an AMG, and how each tool establishes that an AMG call is virus-encoded.

#### Source datasets and benchmark labeling

Benchmark datasets were selected to represent the range of input data types that these tools are designed to process and to span multiple ecosystems. Three input data types (mixed-community metagenomes, viromes, and complete viral genomes) were used across three ecosystems (human gut, aquatic, and soil), yielding nine dataset classes in total. Metagenomes and viromes, sequenced in pairs from the same biological samples, were drawn from previously published datasets, including human fecal samples^91^, aquatic (freshwater) samples from Lake Mendota, WI, USA^92^, and soil microcosm samples^93^. The three largest assemblies from each environment were selected, and all were sequenced and assembled by the JGI with consistent workflows and parameters. Complete viral genomes were obtained by randomly sampling 1,000 high-quality or complete genomes per ecosystem from IMG/VR v4^94^, selecting genomes with soil, marine, and human digestive system (large intestine) ecosystem classifications.

Because metagenomes and viromes contain non-viral sequences, and even complete viral genomes may contain host sequence contamination, an independent viral classification was required to establish ground-truth viral labels for each protein. The virus sequence prediction tool geNomad^39^ was chosen as the viral-origin labeler because it is the field’s currently most widely adopted viral sequence identification tool, is well documented, and has well-characterized performance across diverse sequence types. However, CheckAMG is not fully independent of geNomad, because part of the dataset used to train and evaluate CheckAMG annotate’s viral-origin classifier came from geNomad’s own training data, with additional filtering and label refinements as described above. All benchmark sequences were processed with geNomad v1.11.0 with score calibration enabled and the “--relaxed” flag used to retain predictions across the full score range for downstream filtering. A protein was labeled viral (True Positive) if it was encoded on a sequence predicted to be viral by geNomad with an FDR at or below 0.1, or if it fell within a provirus region predicted by geNomad on a mixed chromosome sequence and that region met the same FDR threshold. Proteins encoded on sequences with no geNomad viral prediction meeting the threshold, or on sequences shorter than 5,000 bp, were labeled non-viral (True Negative). The 5,000 bp minimum sequence length was enforced uniformly across all tools and datasets to remove very short scaffolds where ORF calling and viral identification are unreliable, and because this threshold exceeds the most permissive default used by any of the three benchmarked tools (DRAM-V default: 2,500 bp; VIBRANT default: 1,000 bp; CheckAMG default: 5,000 bp).

#### Tool inputs and execution

To ensure that performance differences among tools reflect their annotation and curation strategies rather than differences in ORF calling or protein translation, DRAM-V was run first on the VirSorter2-identified contigs of all benchmark sequences, and the translated proteins produced by DRAM-V (generated using Prodigal^77^ v2.6.3) were then used as inputs to both VIBRANT and CheckAMG, since both accept amino acid FASTA inputs. DRAM-V v1.5.0 was run, with MMseqs2 13.45111 used both to build and to search its Pfam database, with reference databases KEGG KOfam v113 (accessed 2025-01-27), UniRef90 (2024-06, accessed 2025-01-27), Pfam-A v37.0 (2024-05, accessed 2025-01-27), dbCAN v11 (accessed 2025-01-28), RefSeq v227 viral (2025-01-06, accessed 2025-01-28), MEROPS v12.5 (accessed 2025-01-28), and VOGDB v227 (2024-11-29, accessed 2025-01-28). DRAM-V was run using “--max_auxiliary_score 4” to ensure all candidate AMG predictions up to that auxiliary score threshold were included in outputs, which excludes genes with auxiliary scores of 5 from final AMG predictions. VirSorter2^49^ v2.2 was run on each dataset prior to DRAM-V to generate the required annotation files, using default settings. VIBRANT v1.2.1 was run with databases KEGG KOfam v91 (2019-08-10; accessed 2024-02-01), Pfam-A v32.0 (2018-10; accessed 2024-02-01), and VOGDB v94 (2023-08-22; accessed 2024-02-01). CheckAMG annotate v1.1 was run with the CheckAMG annotate database v1.1 (described above). All tools were executed via Snakemake^95^ v7.32.4 workflows to ensure reproducible and parallelized execution across datasets. The raw AMG prediction output tables from each tool are provided in Supplemental Data 3.

#### Viral prediction confidence mapping across tools

Because the three tools report viral origin evidence using different schemes, AMG predictions from each tool were organized into a set of comparable confidence levels for analysis. For CheckAMG and VIBRANT, confidence levels are cumulative: each higher-confidence level is a strict subset of the one below it. For DRAM-V, four stringency configurations were evaluated, each representing a different combination of auxiliary score cutoff and flag allowance. These configurations are not rank-ordered confidence levels, but the default configuration is divided into three cumulative levels by auxiliary score, matching the CheckAMG and VIBRANT tiers.

For CheckAMG, a gene’s AMG prediction confidence level was assigned based on its per-protein viral origin confidence tier output by the LightGBM classifier: CheckAMG (high) comprises genes classified at high viral origin confidence only; CheckAMG (medium) comprises genes classified at high or medium viral origin confidence; CheckAMG (low) comprises all AMG-predicted genes regardless of viral origin confidence tier (high, medium, or low). This cumulative structure means that CheckAMG (medium) is a superset of CheckAMG (high), and CheckAMG (low) is a superset of CheckAMG (medium).

For VIBRANT, confidence levels are similarly cumulative, derived from VIBRANT’s genome quality designations: VIBRANT (high) comprises AMG predictions from complete circular or high quality draft genomes only; VIBRANT (medium) comprises predictions from complete circular, high quality draft, or medium quality draft genomes; VIBRANT (low) comprises all VIBRANT AMG predictions regardless of genome quality, including low quality draft genomes. Again, VIBRANT (medium) is a superset of VIBRANT (high), and VIBRANT (low) is a superset of VIBRANT (medium).

For DRAM-V, four stringency configurations were defined, each represented as a Boolean AMG-prediction flag per gene. DRAM-V (default), run as DRAM-V distill with --remove_transposons, required an auxiliary score below 4 and the M flag, and excluded the A, V, P and T flags. DRAM-V (allow T), the distill command-line default, was the same but allowed the T flag. DRAM-V (aux 4) was the default with auxiliary scores of 4 admitted, and DRAM-V (aux 4, allow T) additionally allowed the T flag. DRAM-V (default) is contained in each of the other three configurations, and DRAM-V (allow T) and DRAM-V (aux 4) are both contained in DRAM-V (aux 4, allow T), but neither of those two contains the other. Within DRAM-V (default), three cumulative levels were defined by auxiliary score, DRAM-V (aux 1), (aux 2) and (aux 3), holding its calls at auxiliary score 1, at most 2 and at most 3, so DRAM-V (aux 3) is equivalent to DRAM-V (default). Matched levels pair the first, second and third level of each tool, CheckAMG (high) with DRAM-V (aux 1) and VIBRANT (high), CheckAMG (medium) with DRAM-V (aux 2) and VIBRANT (medium), and CheckAMG (low) with DRAM-V (aux 3) and VIBRANT (low). All four configurations are shown in figures for completeness. In DRAM-V v1.5.0, the B flag is assigned to any three consecutive genes carrying the M flag and raises their auxiliary score to 4, and the T flag is assigned to every gene on a scaffold on which any gene has a Pfam hit to one of 17 transposase families (mag_annotator/annotate_vgfs.py).

#### AMG prediction definitions across tools

For CheckAMG, a gene was considered an AMG prediction if it was present in the final curated metabolic gene output (Protein Classification = ‘metabolic’), with the additional requirement that at least one of its HMM annotations matched a profile in the CheckAMG AMG reference table with AMG weight ≥ 0.6 (the default minimum AMG weight applied during the benchmark run). For VIBRANT, a gene was considered an AMG prediction if it appeared in VIBRANT’s AMG individuals output table with a non-null AMG KO annotation (i.e., VIBRANT assigned a KEGG KO to the gene). Each VIBRANT-predicted AMG gene carries exactly one AMG KO. For DRAM-V, a gene was considered an AMG prediction under each of the four stringency configurations as defined in the confidence tier section above. Under the default configuration (auxiliary score < 4, M flag, no A/V/P/T flags), genes with auxiliary scores of 1, 2, or 3 that carry the M flag and lack A, V, P, and T flags are included. The ‘--max_auxiliary_score 4’ flag was used during DRAM-V execution to retain all genes up to auxiliary score 4 in the output, enabling post-hoc application of all four stringency configurations. Since each tool enforces its own default minimum input sequence length, the combined set of AMG predictions was filtered to only contain AMG calls on input scaffolds at least 5 kb in length (the maximum default minimum length across the three tools). The combined, length-filtered AMG predictions were used for downstream comparisons, and the final annotations of the genes called at CheckAMG (high), DRAM-V (default) or VIBRANT (high) are provided in Supplemental Table 8. The geNomad-viral DRAM-V (default) calls with an AMG weight of at least 0.6 that CheckAMG makes at no tier are listed in Supplemental Table 21.

#### Cross-tool functional comparison

To compare AMG calls on a basis independent of any tool’s own criteria, every protein called as an AMG at any tier by any of the three tools was annotated once with KofamScan v1.3.0 and HMMER v3.4 against the KOfam profile database. Two assignment rules were applied. Under the primary rule, a protein took the KO of its hit with the lowest E-value among those with a maximum E-value of 1e-5, with ties broken by the highest score and then by KO. Under the sensitivity rule, a protein took the hit reaching the KofamScan adaptive, family-specific threshold with the highest ratio of score to threshold, with ties broken by the lowest E-value and then by KO, and was counted in a KO class when any of its hits reaching the threshold was in that class. A KO was counted as a “Metabolism KO” when it fell under the A-level “Metabolism” category (09100) of the KEGG BRITE hierarchy, which places 6,490 of 28,008 KOs in that set, and KOs under the “Unclassified: metabolism” (09191) or “Membrane transport” (09131) categories that lie on no Metabolism pathway map were labeled Class II. Each call was then assigned one of four shared KO classes by combining its KO with the geNomad viral call for its encoding sequence, as geNomad viral with a Metabolism KO, geNomad viral with another KO, geNomad viral with no KO, or not geNomad viral. Results in the main text use the primary rule, and counts under the sensitivity rule are reported in Source Data.

#### AMG-weight and cautious-function composition analysis

To compare AMG call quality on a common basis, each tool’s final annotations per gene were looked up in CheckAMG’s scored AMG reference table (AMGs.tsv) to retrieve an AMG weight per annotation. Because DRAM-V can produce multiple Pfam hits per gene, and CheckAMG likewise produces one best-hit annotation per database per gene, each tool can produce multiple annotations per gene, each potentially yielding a different AMG weight. Rather than forcing a single representative annotation, the per-gene AMG weight was taken as the maximum AMG weight across all of a tool’s final passing annotations for that gene (multi-hit-safe operation). A gene with no passing annotations in CheckAMG’s AMG reference table was labeled “annotated, no AMG weight”; a gene with no annotation from CheckAMG’s output at all was labeled “no annotation.” In parallel, CheckAMG’s AMG filter table was used to assign cautious-function flags (essential, nucleotide, glucan, lipid, methyl) per annotation, and flags were OR-aggregated across all passing annotations for a given gene. When a gene matched more than one cautious-function category, the category least common within that gene’s tier was assigned, which is the rule the figure and its legend use. A call’s AMG weight and database source come from its highest-weight annotation, or from its strongest hit where no annotation carries a weight. An EC number that DRAM-V reports as the gene_id of a call is attributed to KEGG, the database of the KOfam hit that supplied it. For genes in the “none” category, a three-way split was applied based on the per-gene best AMG weight: “Not cautious (weight ≥ 0.6),” “Not cautious (weight < 0.6),” or “Not cautious (no weight).” This analysis was applied at the most confident tiers (Figures 3C, 3D) and across all tiers (Extended Data Figures 6B, 6C).

#### Cross-tool disagreement classification and genomic-context validation

To assess whether disagreements in AMG calls between tools were attributable to differences in homology detection sensitivity or to differences in curation and scoring criteria, the full annotation outputs from all HMM searches (for CheckAMG and VIBRANT) and from all DRAM-V database searches were collected for each sample. Annotations were aggregated per gene per tool as sets of hit identifiers and source databases. For the purposes of disagreement classification, a ‘shared database hit’ was defined as a hit identifier present in both the source tool’s final annotations and the other tool’s raw annotation set (regardless of whether the other tool flagged it as an AMG). These per-gene annotation sets are provided in Supplemental Data 4.

For every gene called as an AMG at default/high by one tool, each other tool’s response was classified into one of six mutually exclusive categories, applied in the following precedence order: (a) called only at a lower-stringency setting by the other tool (i.e., the other tool did call the gene as AMG, but only at a lower stringency configuration); (b) agrees at default/high (pass-through, reported separately); (c) source tool’s best-hit database is not queried by the other tool; (d) a shared-database hit identifier exists between the two tools, but the other tool did not flag it as an AMG; (e) the other tool has a hit in a shared database but returned a different top hit; (f) the other tool produced no annotation at all for this gene. For DRAM-V, the same classification was also applied to the calls at DRAM-V (aux 1) and (aux 2). Each source-tier gene set was counted per gene, and the number of genes not on a geNomad-viral region is reported first. An orthogonal ‘gene lacks any viral evidence’ flag was computed separately for each source-tier gene set and reported as an independent count (a gene was considered to lack viral evidence if it was not on a geNomad-viral region, not in a CheckAMG strict viral region, and had no flanking high-V-score viral gene on either side). This flag is not part of the six-category per-comparison breakdown.

To assess which tool is more likely correct in tier-conflict scenarios, each conflicted gene was cross-referenced against four genomic-context signals: (i) geNomad viral classification at FDR ≤ 0.1; (ii) inclusion inside a CheckAMG strict viral region; (iii) the presence of a gene with V-score ≥ 10 within 10 kb on at least one side, and on both sides; and (iv) the character of the nearest MGE-annotated flank (virus-like: nearest MGE V-score ≥ 10; bacterial-like: V-score < 10). A flank whose nearest MGE-annotated gene has no defined V-score counts as neither virus-like nor bacterial-like. The V-score threshold of ≥ 10 matches the minimum viral-V-score gate used in CheckAMG’s ambiguous-region algorithm. Eight pre-specified tier-conflict scenarios were evaluated (Extended Data Figure 7B), defined as genes present in one tool’s high/default tier but only cleared by another tool at a weaker stringency configuration (e.g., genes in CheckAMG high that are also in DRAM-V allow T but not DRAM-V default).

### CheckAMG *de-novo* pipeline, model training, and evaluation

#### CheckAMG-PST finetuning

The *de-novo* module adapts the Protein Set Transformer (PST)^34^ protein-based genome language model for joint viral-origin and AVG prediction. PST uses a Set Transformer architecture that takes all ESM-2 protein embeddings from a scaffold and applies attention across the scaffold to produce context-aware per-protein embeddings. For CheckAMG, we only used the contextualized protein embeddings, dropping the PST decoder so that PST’s aggregated genome-level embeddings are never computed. The PST model was finetuned from a pre-trained PST-TL-P large checkpoint with two triplet-loss objectives: a joint objective that classifies each protein simultaneously by viral origin and AVG-likeness within a single four-class embedding space (defined below), and a local-context objective that gives the model finer resolution for AVG discrimination specifically. Here, AVG-like refers to proteins similar to training proteins with curated metabolic, physiological, or regulatory functions, defined independently of viral origin or genome-context curation. The local-context loss uses a strategy in which, for each labeled AVG, multiple overlapping views of the surrounding neighborhood are used instead of the full scaffold, countering the signal dilution that occurs when a rare AVG is encoded within a long, mostly non-AVG-encoding scaffold. Scaffolds whose protein count does not exceed the context size are excluded from this loss, as local and global views would be identical and provide no useful gradient. Throughout, we refer to the finetuned model as CheckAMG-PST to distinguish it from the original PST.

#### Training and test dataset preparation for CheckAMG-PST

The same training and test protein datasets built for the LightGBM classifier (the same 90%/10% split described above) were relabeled for CheckAMG-PST training along two binary dimensions, viral origin and AVG-like. The viral label is identical to the LightGBM label. The AVG label was assigned from CheckAMG *annotate* v1.1 predictions (metabolic, physiological, or regulatory genes) without filtering by viral origin or genomic context, so the model could learn to separate auxiliary-like from non-auxiliary-like functions independently of viral origin. Because this label is drawn directly from CheckAMG annotate’s own calls rather than an independently curated ground truth, the annotate module’s own criteria for AVG-ness effectively define what CheckAMG-PST could learn to recognize as an AVG (Discussion). The two labels define four mutually exclusive classes: non-viral and non-AVG (class 0), viral and non-AVG (class 1), non-viral and AVG (class 2), and viral and AVG (class 3). Two preprocessing steps preceded embedding. First, scaffolds encoding < 5 proteins were removed. Second, scaffolds encoding > 2,048 proteins were split into non-overlapping chunks of at most 2,048 proteins, which is the maximum PST input size per scaffold. Chunks from the same parent scaffold were assigned new scaffold identifiers, and protein identifiers within split chunks were renamed accordingly.

#### ESM-2 embedding, graph-formatting, and validation split

Each protein FASTA (training chunks and test datasets) was embedded using ESM-2 (esm2_t30_150M)^31^ via the PST library^34^ v1.5.4. Proteins longer than 20,000 amino acids were manually fragmented into non-overlapping segments of at most 20,000 amino acids for embedding, and the resulting fragment embeddings were averaged back into a single per-protein representation. Embeddings were computed using the UW-Madison Center for High-Throughput Computing^96^ and converted to PST graph format with per-protein embeddings, protein and scaffold identifiers, strand orientation, and the ground-truth labels. A validation set was defined within the training data for monitoring training loss during PST fine-tuning. To prevent similar genomes from appearing in both the training and validation fractions, scaffold-level clustering was performed using the protein embeddings (FAISS^97^ v1.13.2 IVF-flat index; 15 nearest neighbors by cosine similarity; Leiden community detection^98^ at resolution = 0.75 in igraph^99^ v1.0.0) and assigning singleton-cluster scaffolds to the validation set, yielding a validation fraction near the 10% target. The labeled training graph with the validation mask and the test datasets are provided as Supplemental Data 5.

#### CheckAMG-PST hyperparameter search and model selection

Fine-tuning was performed in iterative rounds on CUDA GPUs at the UW-Madison Center for High-Throughput Computing (CHTC)^96^, all initialized from PST-TL-P large with batch size 32, learning rate 1e-4, triplet loss margin 1.0 and attention heads fixed at the architecture supplied with PST-TL-P large. Every anchor-positive pair in a minibatch contributes to the loss, but negatives were mined for them in chunks of at most 10,000 pairs at a time to bound memory. Each triplet was weighted by the inverse frequency of its anchor’s class, and the AVG view loss was left unweighted. The AVG triplet loss used a fixed local context of 7 total bidirectional neighboring proteins, substantially smaller than the 25-protein chunk size used for PST’s own genome-context computation, prioritizing tight local resolution around each labeled AVG over broader scaffold context. Models were trained for at most 20 epochs with early stopping on validation loss (patience of 5 epochs, minimum improvement of 0.01). Two positive-mining and two negative-mining strategies were tested, plus an additional restriction of negatives to the most-opposite class, described below.

For positive mining, the “all” strategy uses all same-label pairs as candidate positives, while the “esm_easy” strategy introduced for CheckAMG-PST filters each anchor’s positive to its nearest same-class protein in the input ESM-2 embedding space.

For negative mining, two complementary strategies were tested. “Semihard” negatives are farther from the anchor than its positive but still within the margin, and “hard” negatives are closer to the anchor than its positive. Independently, negatives were either unrestricted or restricted to the most-opposite class present for each anchor. Under the restricted option, the hard or semihard candidate set is computed first, then narrowed to the highest-priority opposite class containing at least one of those candidates. Priority runs by flipping the viral label first (keeping the AVG label), then the AVG label (keeping the viral label), then both, so a viral AVG anchor prefers non-viral AVG negatives, then viral non-AVG, then non-viral non-AVG. If no negative satisfies the hard or semihard condition, the same priority is applied to all differently labeled proteins as a fallback.

Together, this yielded eight configurations (2 positive × 2 semihard/hard × 2 restricted/unrestricted). For each run, overall and AVG-specific triplet losses were logged per epoch with the PyTorch Lightning^100^ CSVLogger, and the best-epoch checkpoint per run was retained. The final CheckAMG-PST checkpoint was selected as the run with the lowest overall validation triplet loss across the sweep. This was the model variant combining the “all” positive mining strategy, “semihard” negative mining, and no opposite-class negative restriction at a local context size of 7, whose best epoch (epoch 13) reached an overall validation loss of 0.235.

#### CheckAMG-PST model evaluation

Candidate checkpoints were evaluated on the same test panel used for the LightGBM classifier plus the integrated-provirus split. Test embeddings were concatenated into one file, per-protein viral and AVG-like probabilities were computed by CheckAMG *de-novo* inference against each candidate’s CheckAMG-PST-embedded training proteins, and per-split predictions were decoded from the concatenated test FASTA. Three per-protein scores were evaluated: the viral-origin probability against the viral label, the AVG-like probability, and the effective AVG probability (their product) against the conjunctive viral-AND-AVG label, which acts as a soft AND gate requiring both lines of evidence to agree before a protein is scored as a candidate AVG. For each score and dataset, PR and ROC curves, AUPRC, and AUROC were computed, and three thresholds per score were selected to define very high, high, medium, and low tiers, choosing for each tier the smallest threshold meeting a per-tier precision floor (0.90, 0.70, 0.50) on every eligible split, or the threshold maximizing worst-case precision when no floor was attainable. The MGE-enriched split was excluded from the viral-score floor, and the MGE-enriched, near-all-host, and integrated-provirus splits from the effective-AVG floor, because their PR curves lay well below the others and would otherwise dominate threshold selection. For the final CheckAMG-PST checkpoint included in the CheckAMG de-novo database v1.1 used here, the viral classifier thresholds were ≥ 0.622 (very high), ≥ 0.051 (high), ≥ 0.001 (medium), and ≥ 0 (low); the raw AVG-like thresholds were ≥ 0.890 (very high), ≥ 0.201 (high), ≥ 0.034 (medium), and ≥ 0 (low); and the effective-AVG thresholds were ≥ 0.956 (very high), ≥ 0.512 (high), ≥ 0.096 (medium), and ≥ 0 (low). Summary results for all test datasets and evaluation frameworks are provided in Supplemental Table 22, and the final CheckAMG-PST model checkpoint in Supplemental Data 6.

#### CheckAMG de-novo inference workflow

The CheckAMG *de-novo* inference pipeline is implemented as a Snakemake workflow with a CLI that accepts either nucleotide or protein FASTA input (or pre-computed ESM-2 or CheckAMG-PST embeddings) and produces per-protein AVG-like and viral-like probability scores (Supplemental Figure 2). When nucleotide input is provided, ORF prediction and translation are first performed using pyrodigal-GV^39,76,77^ (as in the CheckAMG *annotate* module). Input proteins are then filtered by a minimum of 5 proteins per scaffold (by default), a maximum of 2,048 proteins per scaffold (scaffolds exceeding this limit are split), and a maximum protein length of 20,000 amino acids (proteins exceeding this limit are fragmented for embedding, then their fragment embeddings are averaged). Filtered proteins are then embedded using ESM-2 (esm2_t30_150M) and converted to graph format as described above, producing per-protein ESM-2 embeddings stored alongside scaffold grouping metadata. The CheckAMG-PST model then processes these graph-formatted embeddings in batches to produce context-aware CheckAMG-PST embeddings.

Inference is performed by nearest-neighbor search: query CheckAMG-PST embeddings are compared against the pre-embedded training data using a FAISS IVF-flat index, retrieving the *k* nearest training proteins. We incrementally tested multiple values of *k* until we reached an optimal value of *k* = 20 that balanced cross-validation precision and model inference runtime to set a default *k* value for the CheckAMG *de-novo* model inference workflow. Predicted class probabilities for each query protein are computed by distance-weighted voting among the *k* neighbors, where weights are computed via the PyTorch function “softmax” over negative distances scaled by the standard deviation of finite distances. The resulting four-class probability vector (classes 0–3 as defined above) is decomposed into two independent binary probabilities: the viral-origin probability (sum of class 1 and class 3 probabilities) and the AVG-like probability (sum of class 2 and class 3 probabilities). Both probabilities are reported per protein in the final output (Supplemental Figure 2).

#### Comparing CheckAMG-PST and LightGBM viral origin predictions

Per-protein viral probabilities from both classifiers were collected for all test datasets and compared by Pearson and Spearman correlation, per dataset and combined. The AVG prediction sets were intersected to identify proteins called viral by both, by CheckAMG-PST only, or by the LightGBM only; agreement was summarized with weighted Cohen’s kappa, and binary calls (medium confidence or higher) compared for set encompassment and for precision and recall against the ground-truth viral label. Sensitivity to scaffold length was assessed by relating each model’s per-protein probability and correctness to log gene count per scaffold (Spearman correlation and a logistic regression with a model-by-length interaction).

### CheckAMG application to MetaVR soil and human-gut viral genomes

#### Data sources and quality control

Soil and human-gut viral genomes were obtained from the Meta-virus resource (MetaVR)^19^ by exporting all uncultivated viral genome (UViG) metadata records (as of May 2026) labeled “Environmental; Terrestrial; Soil” (n = 2,054,095 UViGs; 1,284,646 vOTUs; https://www.meta-virome.org/Uvigs?hieco=Environmental&hicat=Terrestrial&hitype=Soil&pageSize=20) and “Host-associated; Mammals (Human); Digestive system” (n = 499,416 UViGs; 229,824 vOTUs; https://www.meta-virome.org/Uvigs?hieco=Host-associated&hicat=Mammals%3A%20Human&hitype=Digestive%20system&pageSize=20). The corresponding nucleotide sequences were extracted from the MetaVR sequences (IMGVR5_UViG.fna).

Quality filtering followed the criteria of Graham et al.^50^: (1) linear scaffolds ≥ 1 kb with CheckV^22^ Complete, High-quality, or Medium-quality completeness, or scaffolds ≥ 1 kb with direct or inverted terminal repeats; (2) linear scaffolds ≥ 10 kb with geNomad^39^ virus score > 0.8 and at least one viral hallmark gene; and (3) linear scaffolds > 5 kb and < 10 kb with geNomad virus score > 0.9 and at least one viral hallmark gene. The MetaVR metadata does not include columns for the “geNomad virus marker” and “marker enrichment” terms used by Graham et al., so those were not applied; otherwise the criteria are unchanged. Because MetaVR includes proviruses (which Graham et al. excluded), an additional rule was added to retain proviruses ≥ 10 kb that were CheckV Complete, High-quality, or Medium-quality with CheckV contamination < 5%, geNomad virus score > 0.8, and at least one viral hallmark gene. After filtering, 767,946 soil UViGs and 238,034 human-gut UViGs remained.

#### AVG prediction

The filtered UViG nucleotide sequences were extracted from IMGVR5_UViG.fna. CheckAMG *annotate* v1.1 with database v1.1 (described above) was then run on each dataset with the options --min-len 1 --min-orf 1 since inputs were already quality-filtered. For the *de-novo* module, the filtered single-scaffold protein FASTAs produced by CheckAMG *annotate*’s pyrodigal-GV step were used directly as input. CheckAMG *de-novo* v1.1 was run on GPU nodes provided by the UW–Madison Center for High-Throughput Computing^96^ using the final selected CheckAMG-PST checkpoint, the corresponding precomputed FAISS IVF-flat index of training-protein CheckAMG-PST embeddings, and the matching training-protein label file. Protein sequences (in amino-acid FASTA format) for all AVGs predicted by CheckAMG *annotate* and *de-novo* are provided in Supplemental Data 7.

#### Sequence-similarity invisible AVGs and auxiliary category assignment

Among *de-novo*-only AVGs (AVGs identified by the *de-novo* module but not by *annotate*), those with no candidate functional annotation from *annotate* in any of the seven reference databases are referred to as sequence-similarity invisible. A functional label was propagated to each of these proteins in two tiers. All labeled training proteins and all applied AVGs were clustered together in one MMseqs2^86^ (v18.8cc5c) universe of 1,500,646 sequences at each of four sequence-identity thresholds (cluster, --min-seq-id 0.3, 0.5, 0.7 and 0.9, -c 0.8 --cov-mode 0 -s 7.5). The first tier assigned the majority label of the labeled training proteins in a protein’s cluster, where that cluster held at least one labeled protein and the vote was not tied. The second tier, used only where the first could not reach, propagated the label of the protein’s single nearest labeled training protein in CheckAMG-PST embedding space. At each of three hierarchical levels, from most to least specific (a specific function name, the L1 category assigned to that function name in the CheckAMG reference classification table (see Supplemental Tables 23, 24, and 25 for AMG, APG, and AReG category assignments, respectively), or a broader metabolic, physiological, or regulatory category), the cluster majority was taken first, at the identity threshold selected for that level on the calibration split (50% for specific function names and 30% for both category levels), and otherwise the nearest neighbor’s label was assigned only if the calibrated probability of that label being correct, given the embedding distance to the neighbor, met a 90% realized-precision target (calibration described below). Each protein was labeled at the deepest level meeting one of these two criteria. Proteins with insufficient evidence at every level were left without a label rather than being assigned an unreliable one.

#### Cross-validation of auxiliary category assignments

CheckAMG-PST’s ability to assign a broad auxiliary category (metabolic, physiological, or regulatory) was tested on 133,744 distinct proteins withheld entirely from training, using a single-nearest-neighbor rule (Supplemental Table 26; Extended Data Figure 9). For each withheld test protein, the category assigned to it was that of its single nearest labeled training neighbor in CheckAMG-PST embedding space. Four comparisons were made. First, accuracy on the true category labels (0.895) was compared against accuracy after the training labels were randomly reassigned (a chance-level control; 0.515) and against always predicting the single most common category (a naive baseline; 0.668). Second, accuracy was compared between test proteins whose nearest training match came from a bacterial genome (0.920) versus a viral genome (0.890). Third, the fraction of confident predictions (nearest-neighbor distance within the threshold defined below) was compared between true auxiliary genes (7.24%) and a negative-control set of non-auxiliary structural genes (0.0035%). Fourth, the classification accuracy of the finetuned CheckAMG-PST model (0.896) was compared with that of an unmodified ESM-2 model (esm2_t30_150M) that received none of CheckAMG-PST’s additional training (0.988), at a matched sample size. As a stress test of how much of this performance rests on close sequence relatives, the raw nearest-neighbor rule, without its calibration gate, was re-run with same-cluster neighbors masked, so that each protein took its nearest surviving neighbor outside its own MMseqs2^86^ cluster (41,903 of the 41,957 reporting proteins; Supplemental Table 27). Broad-category accuracy fell from 88.6% to 86.0%, L1 accuracy from 57.8% to 49.3%, and specific-function accuracy from 37.4% to 26.1%. This validation strategy follows that used for GeneCLR-DF^32^, a model that also combines ESM-2 protein embeddings with a genome-context transformer, though for a different task (predicting antiphage defense systems in bacterial genomes).

#### Calibration of functional-label assignment depth

Each protein included for functional propagation was assigned a label at the deepest level of a three-level hierarchy (a specific function name, an “L1” function category from CheckAMG *annotate* classifications, or a broad metabolic, physiological, or regulatory category) if the calibrated probability of the label being correct met a chosen precision target (90%). Proteins with insufficient evidence at every level received no propagated functional label. A probability of correctness was calibrated separately for each level, for the second tier only, since the first tier is a majority vote rather than a scored prediction. The evaluation set was partitioned three ways: a fitting split of 55,942 proteins used to train the correctness model, a calibration split of 41,957 proteins used to set the admission thresholds, and a reporting split of 41,957 proteins, held out from both, on which all reported coverage and realized precision were measured. For each level, the admission threshold was set by inverting the precision-coverage relationship on the calibration split, restricted to the proteins the first tier could not reach at that level. Candidate labels were ranked by calibrated probability, and the threshold was taken as the lowest probability at which cumulative precision still met the target. Realized precision was then measured on the reporting split, where the two-tier rule labeled 92.2% of proteins at 93.7% precision at the depth assigned. Within that, the cluster tier reached 0.950 precision for specific function names, 0.946 for L1 categories and 0.994 for broad categories, and the embedding tier 0.895, 0.916 and 0.887, each against the 0.90 target.

#### AVG protein embedding clustering and “biome-skew” calculation

CheckAMG-PST embeddings for all identified AVGs (both *annotate* and *de-novo*, including sequence-similarity invisible proteins) were normalized to unit length and projected to two dimensions with UMAP (n_neighbors = 30, min_dist = 0.1, cosine metric), then clustered on the resulting coordinates with HDBSCAN (min_cluster_size = 300, min_samples = 30). Each cluster’s soil fraction was the proportion of its member proteins that originated from a soil viral genome. Clusters were classified as soil-specific (soil fraction at least 0.85), soil-leaning (0.65 to 0.85), mixed (0.35 to 0.65), gut-leaning (0.15 to 0.35), or gut-specific (at most 0.15).

#### Biome- and host-enrichment of AVG functions

Functions were ranked by how often they occur among CheckAMG *annotate*-derived (reference-annotated) AMG proteins and split into three equal-sized groups. Only functions in the most common third were tested, because propagated labels for rarer functions are markedly less precise (micro precision in the rare tercile is 0.054 at the specific level, see Supplemental Results). For each auxiliary function (at specific or L1 depth) that fell in this common-frequency group and was carried by at least 20 AVG proteins, the number of soil and gut AVG proteins carrying that function was compared against the total number of AVG proteins in each biome using a two-sided Fisher’s exact test, with Benjamini-Hochberg (BH) correction across functions, calling a function enriched at *q* < 0.05 together with an odds ratio > 1.5 or < 0.667. This analysis includes both reference-annotated and propagated functional labels. Propagated labels are included only at specific or L1 depth and only for functions in this common-frequency group. A protein counts toward every function category it carries, whether reference-annotated or propagated, so proteins with more than one relevant function contribute to more than one test. Multi-membership is handled identically regardless of whether the function came from reference annotation or propagation, so that the two label sources remain comparable. This test is reported in the Supplemental Results as a concordance check between the two label sources rather than as a separate enrichment result.

For testing enrichment of AMG categories between biomes, a genome-level test was applied separately at L1 and L2 category resolution (Supplemental Table 14). Here the unit is the genome rather than the AMG protein. For each category, the number of genomes in each biome encoding at least one gene in that category was compared against the total number of *annotate*-screened genomes in that biome (767,786 soil and 238,034 human gut), so that a category present in many copies within one genome counts once. The soil total is 160 fewer than the 767,946 genomes retained by quality filtering, because those 160 genomes were not screened by CheckAMG annotate. Reference-annotated genes contribute at whatever categories CheckAMG annotate assigns them (see Supplemental Table 23 for AMG category assignments for all CheckAMG reference AMGs). Propagated genes contribute only where a specific function name was assigned, which is then expanded through the reference mapping to every (L1, L2) parent that function belongs to. Genes labeled only to L1 depth have no basis for an L2 assignment and were excluded from the L2 test rather than imputed. Each category was tested exactly once at each level, with L2 categories belonging to more than one L1 grouped as “Multiple L1 categories” for visualization only. Categories carried by fewer than 20 genomes in total were not tested.

Gut viral genomes in this collection are larger than soil genomes, a median of 18 predicted genes against 13, so a genome-level presence test is confounded by genome size: a genome with more genes has more opportunity to carry any given category. Categories were therefore tested with a Cochran-Mantel-Haenszel test stratified by quintiles of total gene count, which compares biomes within strata of comparable size and conditions on nothing downstream of the outcome. A stratified test was preferred to a regression adjusted for the number of labeled genes, because that covariate is itself partly determined by the outcome, and total gene count was preferred to genome length as the stratifying variable, because presence-absence scoring makes the opportunity scale with gene number while the recorded length of a provirus is that of its host genome. The estimator was chosen on the design of the test rather than on the result it produced, and the unadjusted two-sided Fisher exact result is reported alongside every adjusted one in Supplemental Table 14. Categories were called enriched at *q* < 0.05 (BH-adjusted, applied separately within each level) together with an absolute log_2_ odds ratio > 0.585 (odds ratio > 1.5 or < 0.667). Categories present in only one biome, for which the odds ratio is undefined, received a Haldane-Anscombe continuity correction of one half per cell for the effect-size estimate only, and their *P*-values were unaffected.

Each AMG category was also tested for over-representation within a particular predicted host phylum (from the iPHoP^101^ host predictions in the MetaVR metadata for the UViGs) relative to the rest of the same biome, using the same genome-level two-sided Fisher’s exact test at the resolution of L2-level categories assigned by CheckAMG *annotate* (Extended Data Figure 10; Supplemental Table 23). CheckAMG *annotate*-annotated genes contributed at their curated L2 categories, whereas propagated genes contributed only where a specific function name was assigned, mapped through the reference to its L2 parent categories. A host phylum was tested only if at least 300 (soil) or 150 (human gut) genomes were assigned to it, a category had to be present in at least 10 genomes within that biome, and at least 3 genomes in the tested phylum had to carry it. BH correction was applied separately within each biome, and a category was called enriched at *q* < 0.05 together with an odds ratio > 1.5. No continuity correction was applied to these tests, so categories absent from the remainder of the biome have an undefined (infinite) odds ratio.

#### Defense/anti-defense searches and structural alignments of sequence-similarity invisible AVGs

To assess whether the set of 109,107 sequence-similarity invisible AVGs (defined above) could be assigned putative functions that CheckAMG did not detect, they were searched using two independent approaches.

First, the sequence-similarity invisible AVGs were directly searched with pyhmmer against the full database of 1,332 DefenseFinder and AntiDefenseFinder profile HMM models (v3.1.0)^51^. The resulting hits were evaluated separately at each model’s standard confidence threshold (curated and provided in the DefenseFinder database v3.1.0) and at a more permissive threshold (E ≤ 1e-3).

Second, Foldseek (v10.941cd33) in ProstT5 structure-aware sequence mode^102^ (ProstT5 prostt5-f16 weights) was used to search every sequence-similarity invisible protein against structures in PDB100^103^ (2024-01-01 release), BFVD^104^ (Big Fantastic Virus Database, v2023_02_v2), and AFDB50^105^ (AlphaFold DB, UniProt50-clustered), with parameters -e 0.01 -s 9.5 --max-seqs 300. We considered a confident structural match to be one with an alignment E-value ≤ 1e-5 and at least 50% query coverage. Foldseek was run with a custom nine-column output layout (query, target, target description, fident, alnlen, evalue, bits, qcov, tcov), so E-value and bitscore are columns six and seven rather than their standard tabular positions. Structural matches were considered to have “known” (predicted) functions if the matched target’s description stated a specific molecular function or named family. Matches to targets that contained descriptions indicative of a hypothetical, uncharacterized, or DUF (domain of unknown function) entry were labeled as matches to structures with “unknown” functions.

A match counted as named only when the target description, or for BFVD targets the UniProt protein name of the target accession, was longer than three characters and matched none of the uncharacterized or assembly-artifact patterns, and when the best such match for that protein, ranked first on carrying a specific rather than a domain-level name and then on bitscore, carried a specific function name. Of the 72,393 proteins with a named structural match, 64,935 were named from AFDB50, 6,979 from BFVD, and 479 from PDB. Each named match was also assigned a functional theme by keyword rules applied to the match description, so a theme reports what the matched structure is called rather than the auxiliary class CheckAMG assigns. Names can therefore fall in a theme that their biological role does not match, as for peptidase S24, the self-cleavage domain of LexA-family repressors, and for methyltransferase names, which group with metabolic functions whether the enzyme acts on a host metabolite or on nucleic acid.

The 109,107 sequence-similarity invisible AVGs were clustered into embedding-based protein families using Leiden community detection (objective function: modularity) on a cosine k-nearest-neighbor graph built from their CheckAMG-PST embeddings (L2-normalized; FAISS HNSW inner-product index, M = 32, efConstruction = 200, efSearch = 64; 15 nearest neighbors per protein, self excluded), yielding 291 families. A family was considered “novel” only if none of its members were recovered by DefenseFinder, AntiDefenseFinder, or structural homology searches (56 families, 100 proteins). We refer to these 100 proteins as novel AVGs. This is a stricter, family-level definition than the protein-level count of 30,412 AVGs unresolved by any of these approaches reported in the Results, because most families containing an unresolved protein also contain at least one identified member.

#### Use of large language models

Large language models were used under the authors’ direction to assist with debugging, organizing code and the software repository, editing code for testing and plotting, and checking the reproducibility of analyses and the agreement of reported values and cross-references with analysis outputs. They were also used to check grammar and spelling and to make minor wording refinements to the manuscript text. Study design, methods, analyses, interpretation, and conclusions were determined by the authors, and all AI-assisted code, figures, and text were reviewed and validated against the underlying data by the authors. The authors take full responsibility for the content of this work.

## Data availability

The CheckAMG reference database (the AMG, APG, and AReG HMM reference tables and associated filter and category tables) is available on Zenodo using the DOI 10.5281/zenodo.21776005. Supplemental data needed to reproduce the analyses reported here, including the datasets used to train and evaluate the LightGBM classifier (amino acid sequences and per-protein annotations, labels, and features), the raw outputs and compiled annotations from the CheckAMG, DRAM-V, and VIBRANT benchmark runs, and the datasets used to train and test the CheckAMG-PST model (amino acid sequences, per-protein labels, protein embeddings) are available on Zenodo using the DOI 10.5281/zenodo.24025068. Source data for the main and extended data figures are provided with this manuscript.

## Code availability

CheckAMG is available on GitHub at github.com/AnantharamanLab/CheckAMG, distributed under the GPL-3.0 license. Scripts and code notebooks used to perform the analyses here and to generate the figures and tables are available on GitHub at the CheckAMG repository.

## Acknowledgments

We thank the University of Wisconsin-Madison Center for High Throughput Computing (CHTC) for computational resources and assistance in their use. JCK was supported by a National Science Foundation Graduate Research Fellowship under grant no. 2137424. CM was supported by a University of Wisconsin-Madison SciMed Graduate Research Scholars fellowship and by a National Science Foundation Graduate Research Fellowship. JMW was supported by the Grayce B. Kerr Early Career Fellowship. This research was supported by a National Institute of General Medical Sciences of the National Institutes of Health award (R35GM143024, KA) and a National Science Foundation Advances in Biological Informatics (ABI) award (2149505, BB, MU, and MBS). K.A. also acknowledges support by Human Frontier Science Program research grant RGP018/2024.

## Author contributions

J.C.K. and K.A. conceived the study. J.C.K. designed and implemented CheckAMG, built the reference databases and curation framework, trained and evaluated the LightGBM and CheckAMG-PST models, performed the benchmarking and large-scale analyses, generated the figures, wrote the initial manuscript, and revised the final manuscript with input from all authors. C.M. implemented the code for CheckAMG-PST finetuning, advised on model training, and led CheckAMG-PST model evaluation. J.M.W., B.B., M.U., and M.B.S. provided conceptual input on auxiliary viral gene curation and manuscript structure. J.M.W. and M.U. tested the software. K.A. supervised the project and acquired funding. All authors reviewed and edited the manuscript and approved the final version.

## Extended Data Figures

**Extended Data Figure 1.**
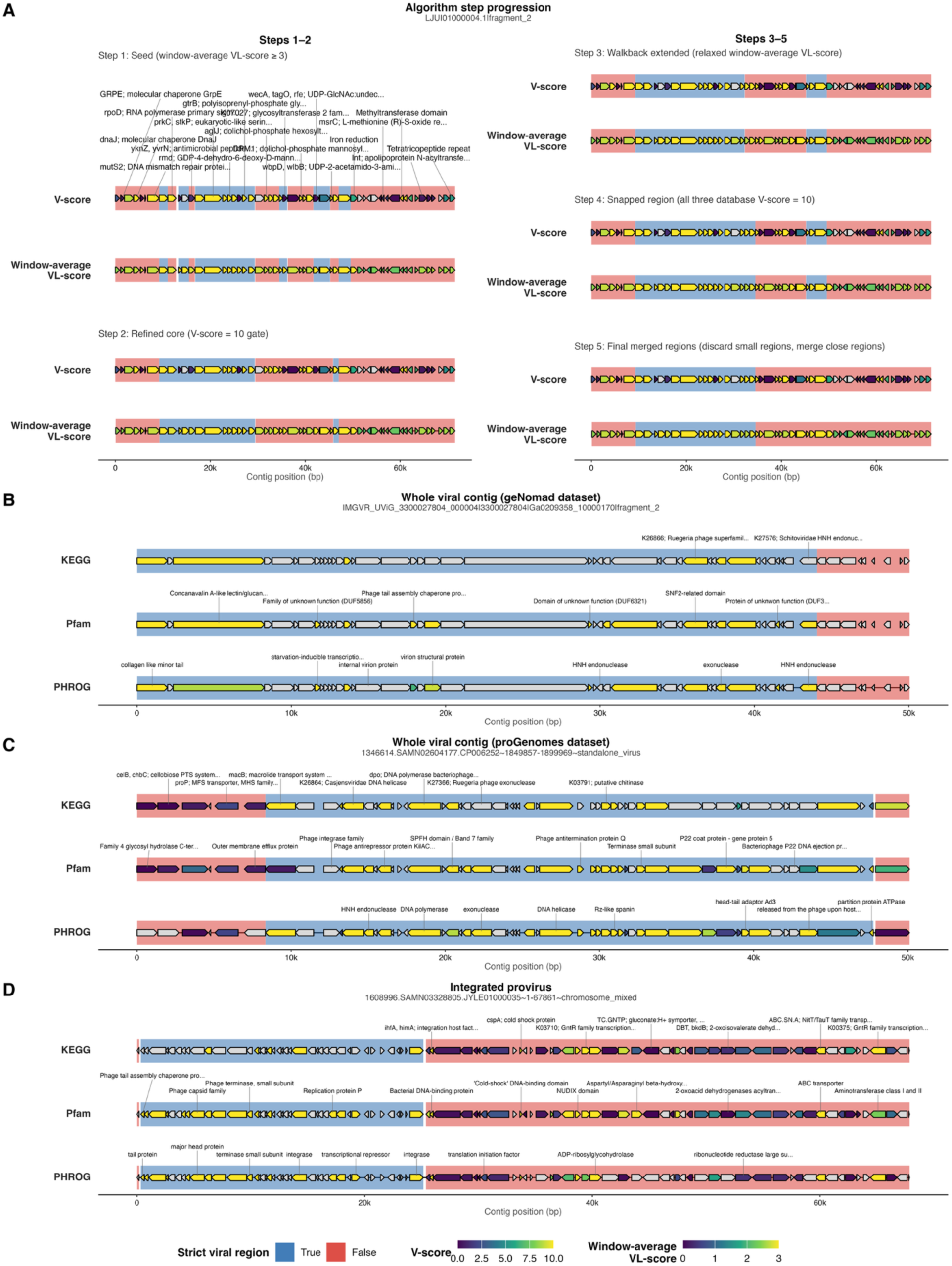
Strict viral regions identified with the V- and V_L_-score-based algorithm in CheckAMG *annotate*. Strict viral regions are highlighted in blue and ambiguous regions in red. **(A)** Step-by-step progression of the algorithm on a single scaffold, showing the gene track, per-gene V-score (colored by the maximum of KEGG, Pfam, and PHROG), and the sliding-window-average V_L_-score. Each step refines the previous call: Step 1 seeds genes by window-average V_L_-score; Step 2 refines to a V-score-passing core; Step 3 walks boundaries back through bridge-eligible genes; Step 4 optionally snaps boundaries to a nearby strong-V anchor gene; Step 5 discards short regions and merges close ones. **(B)** Example viral scaffold from the geNomad-sourced data. **(C)** Example viral scaffold from the proGenomes/proMGE-sourced data, where the algorithm reproduces the source viral label. **(D)** Example integrated provirus in a proGenomes/proMGE scaffold, where the algorithm refines the proMGE-reported boundaries.

**Extended Data Figure 2.**
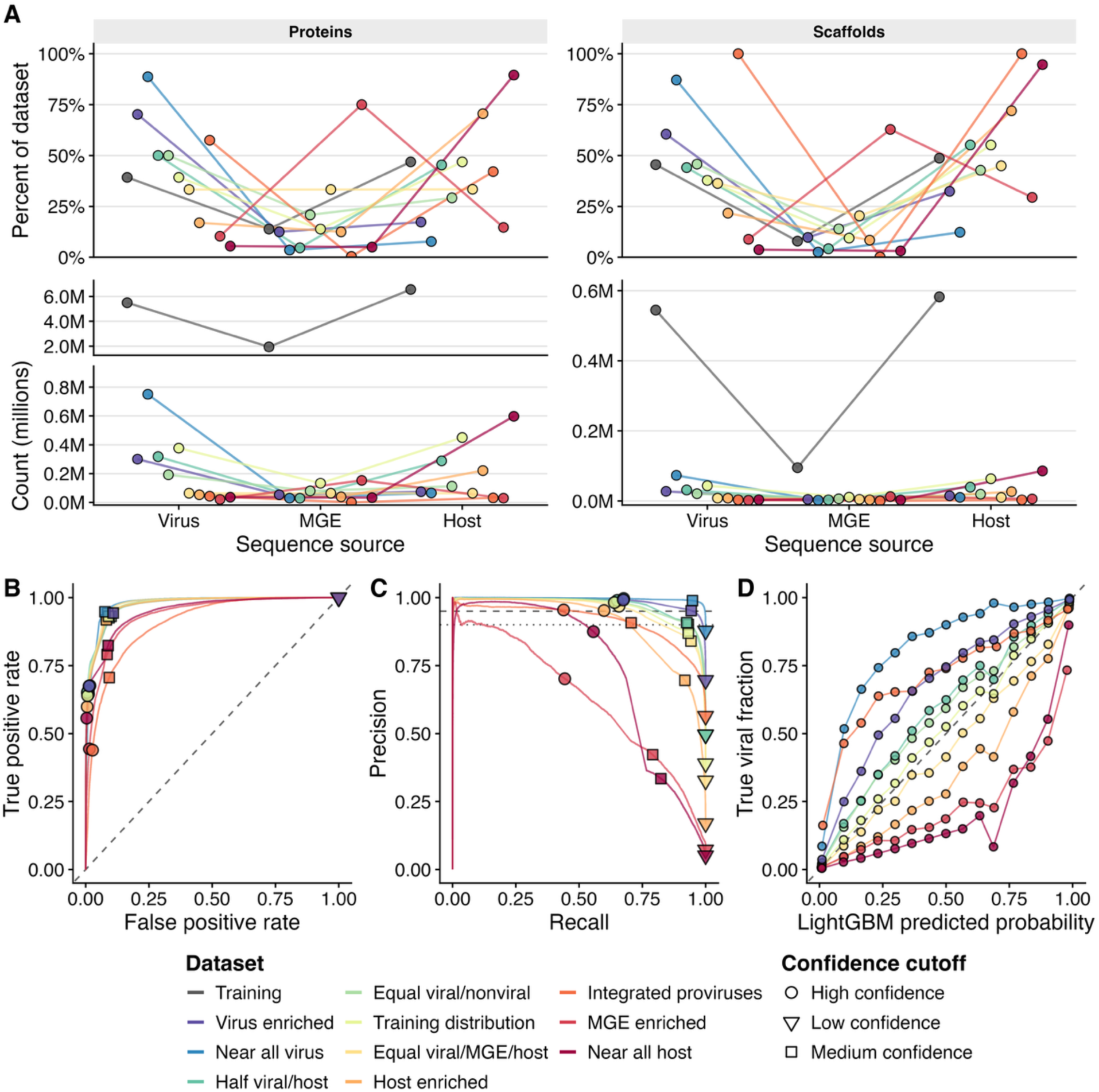
Performance of the LightGBM viral protein classifier across test sets spanning a range of viral, host, and MGE compositions. **(A)** Composition of the training set and each test set by sequence source, shown at the level of proteins (left) and scaffolds (right). The upper row shows the percentage of each dataset’s proteins or scaffolds from each sequence source (viruses, MGEs, and host chromosomes), and the lower row shows the corresponding counts in millions. The protein count axis is broken to fit the much larger training set. **(B)** Receiver operating characteristic (ROC) curves; the dashed diagonal indicates random performance, and markers denote operating thresholds for high (circle), medium (square), and low (inverted triangle) confidence. **(C)** Precision–recall (PR) curves; horizontal reference lines mark precision of 0.90 and 0.95. **(D)** Calibration diagram showing the true viral fraction versus mean predicted probability within each of 15 uniform-width bins per dataset; the dashed diagonal represents perfect calibration.

**Extended Data Figure 3.**
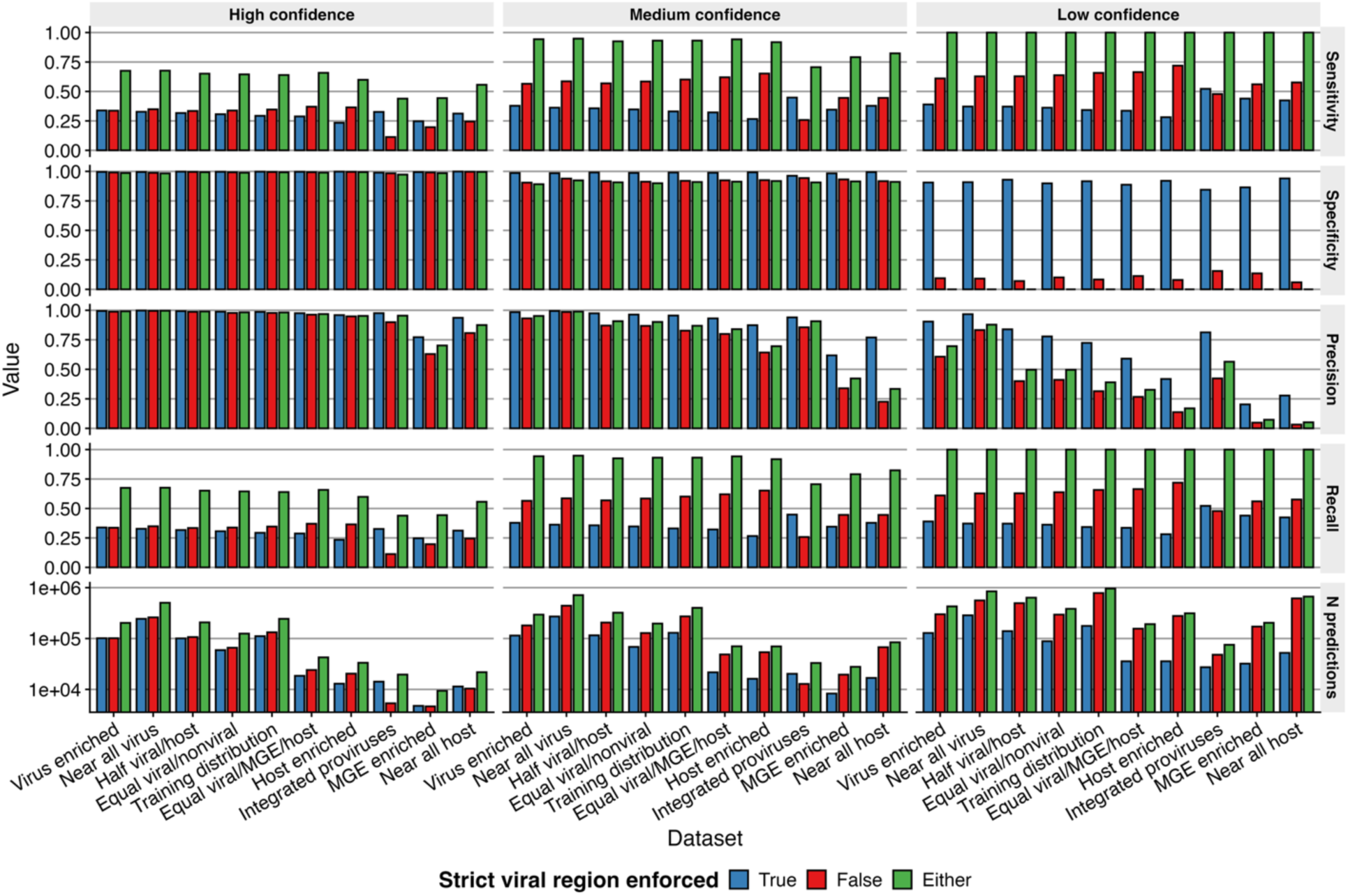
Effect of enforcing the strict viral region requirement on LightGBM predictions across test datasets and confidence tiers. Per-test-dataset sensitivity, specificity, precision, recall, and number of predictions (log_10_ scale) when the LightGBM call is further restricted to proteins inside a strict viral region identified by CheckAMG *annotate*’s V- and V_L_-score-based ambiguous-region algorithm (Methods; Extended Data Figure 1). Bars are colored by whether the strict-region requirement is enforced (blue), not enforced (red), or whether either the LightGBM call or the strict-region criterion is required (green). Rows correspond to the high, medium, and low confidence tiers of the LightGBM classifier.

**Extended Data Figure 4.**
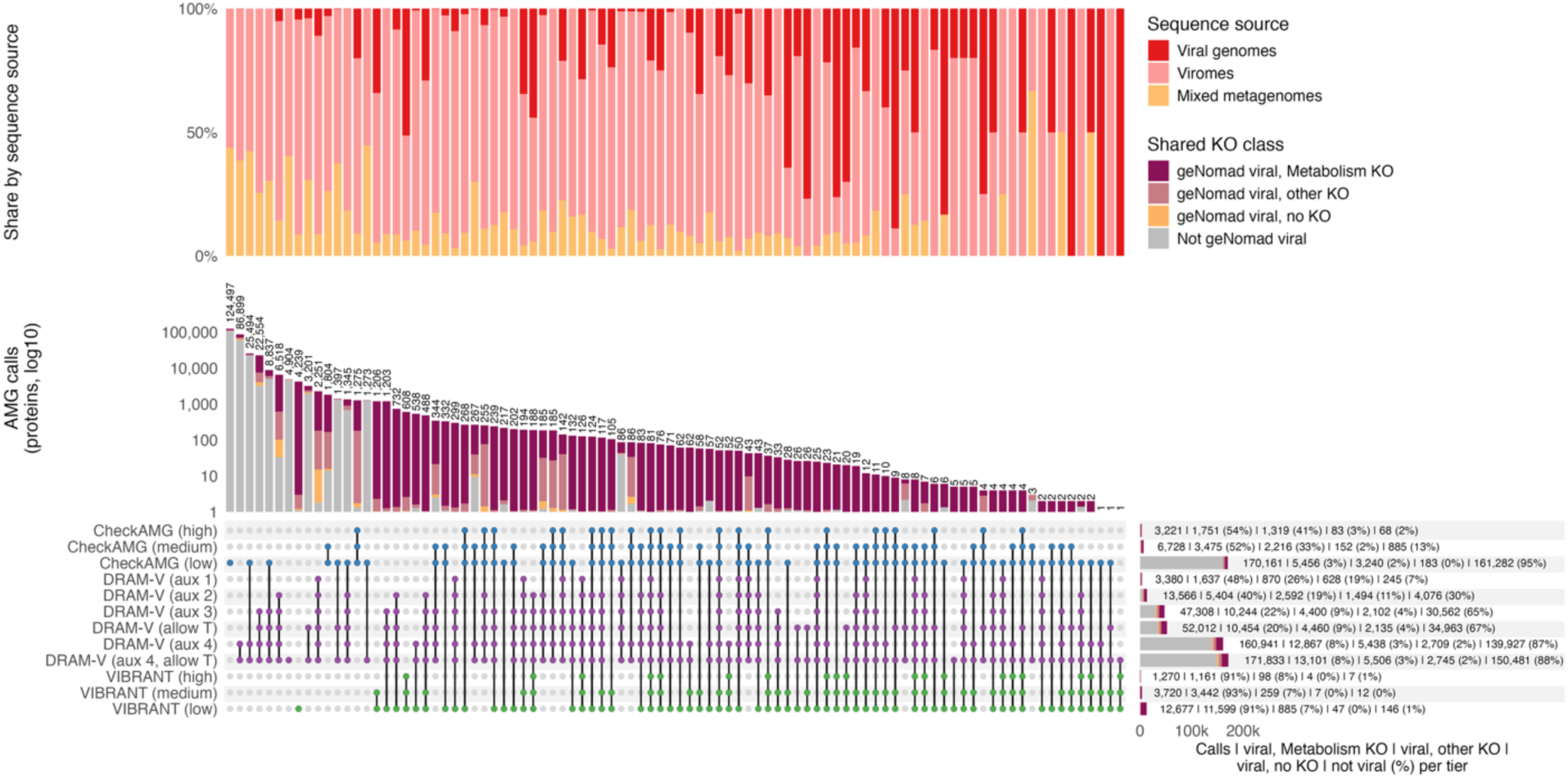
Overlap of AMG calls across all tool tiers and DRAM-V configurations, by sequence source and shared KO class. Rows (sets) are the AMGs called at each tier, and columns (intersections) show the number of AMGs called only by the tiers marked with connected points. All tiers are cumulative, DRAM-V (default) is shown as DRAM-V (aux 1), (aux 2) and (aux 3) as in Figure 3, and DRAM-V (allow T), (aux 4) and (aux 4, allow T) are full configurations that include the default calls. The upper panel gives the share of each intersection by the source of the sequence encoding the call, as viral genomes, viromes or mixed metagenomes. Intersection and set-size bars are split by the shared KO classes of Figure 3A. Intersection counts are on a log scale, so each class takes its share of a bar’s drawn length rather than ending at its count, and cover all 306,534 genes called as an AMG at any tier by any tool, with no intersection omitted. Horizontal bars show the tier totals, labeled with the number and share of calls in each class.

**Extended Data Figure 5.**
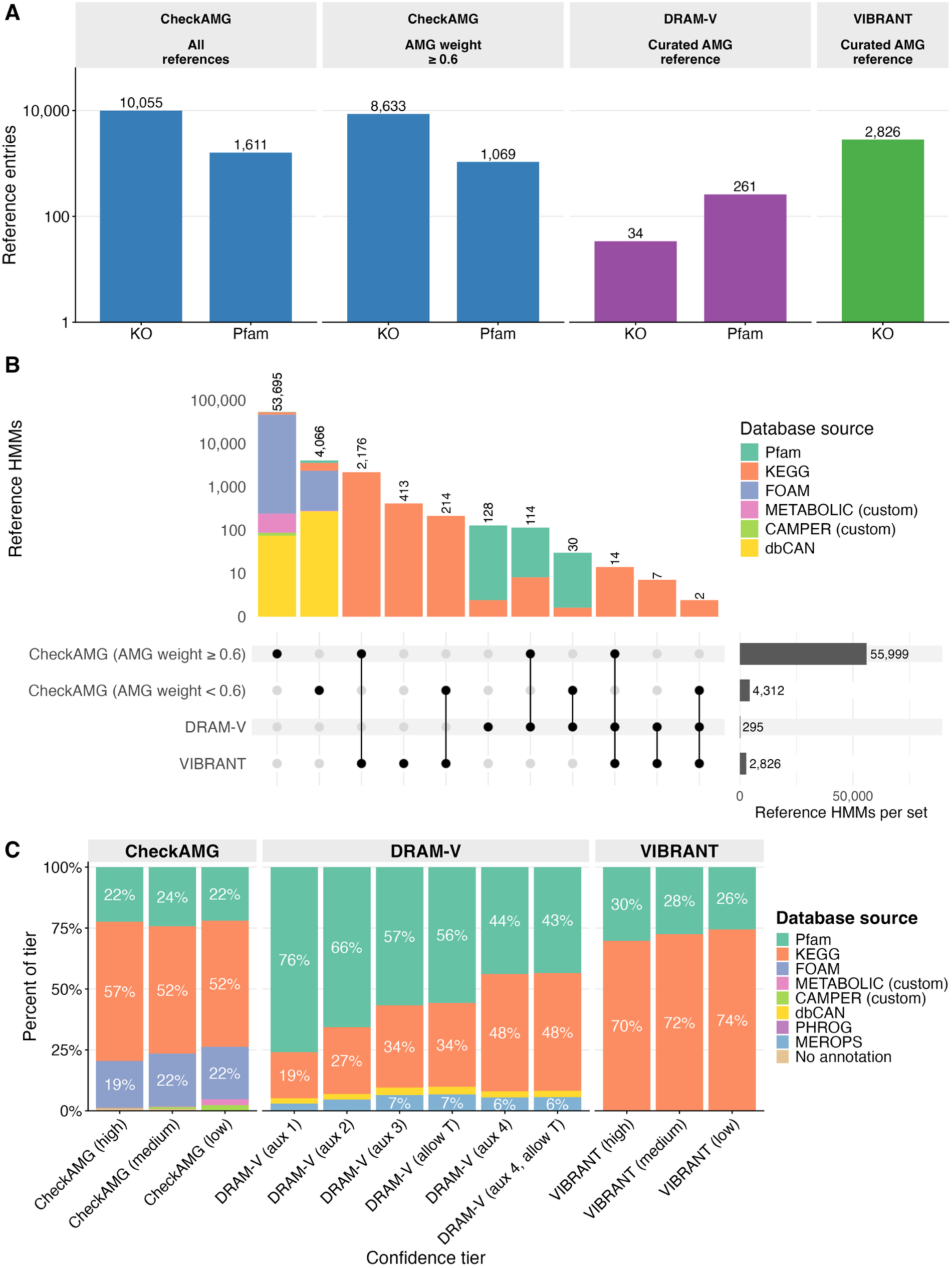
Reference AMG sets used by CheckAMG, DRAM-V, and VIBRANT are largely distinct. **(A)** Reference database size per tool (KO and Pfam entry counts). For CheckAMG, counts are shown for all references and for the subset at AMG weight at least 0.6. **(B)** UpSet plot of overlapping and unique reference AMG HMM profiles across the three tools. Profiles from Pfam, KEGG, and dbCAN that were also present in the METABOLIC or CAMPER databases are not included in the METABOLIC or CAMPER intersection and set size color mappings. Only the HMM profiles that were custom built by METABOLIC or CAMPER are included in their intersection and set size color mapping. **(C)** Composition of AMG annotations by source database for benchmarked predictions produced by each tool and confidence tier. All tiers are cumulative, so a call is counted at every tier it belongs to. DRAM-V (default) is shown as DRAM-V (aux 1), (aux 2) and (aux 3), its calls at auxiliary score 1, at most 2 and at most 3, and DRAM-V (allow T), (aux 4) and (aux 4, allow T) are shown in full, each including the default calls. The databases represented are KEGG, Pfam, FOAM, CAMPER, METABOLIC and PHROG, and for DRAM-V also MEROPS and dbCAN, and where no annotation of a call carries an AMG weight the call takes the source of its strongest hit. CheckAMG’s dbCAN annotations are not present in the benchmark annotation tables and so are not represented here.

**Extended Data Figure 6.**
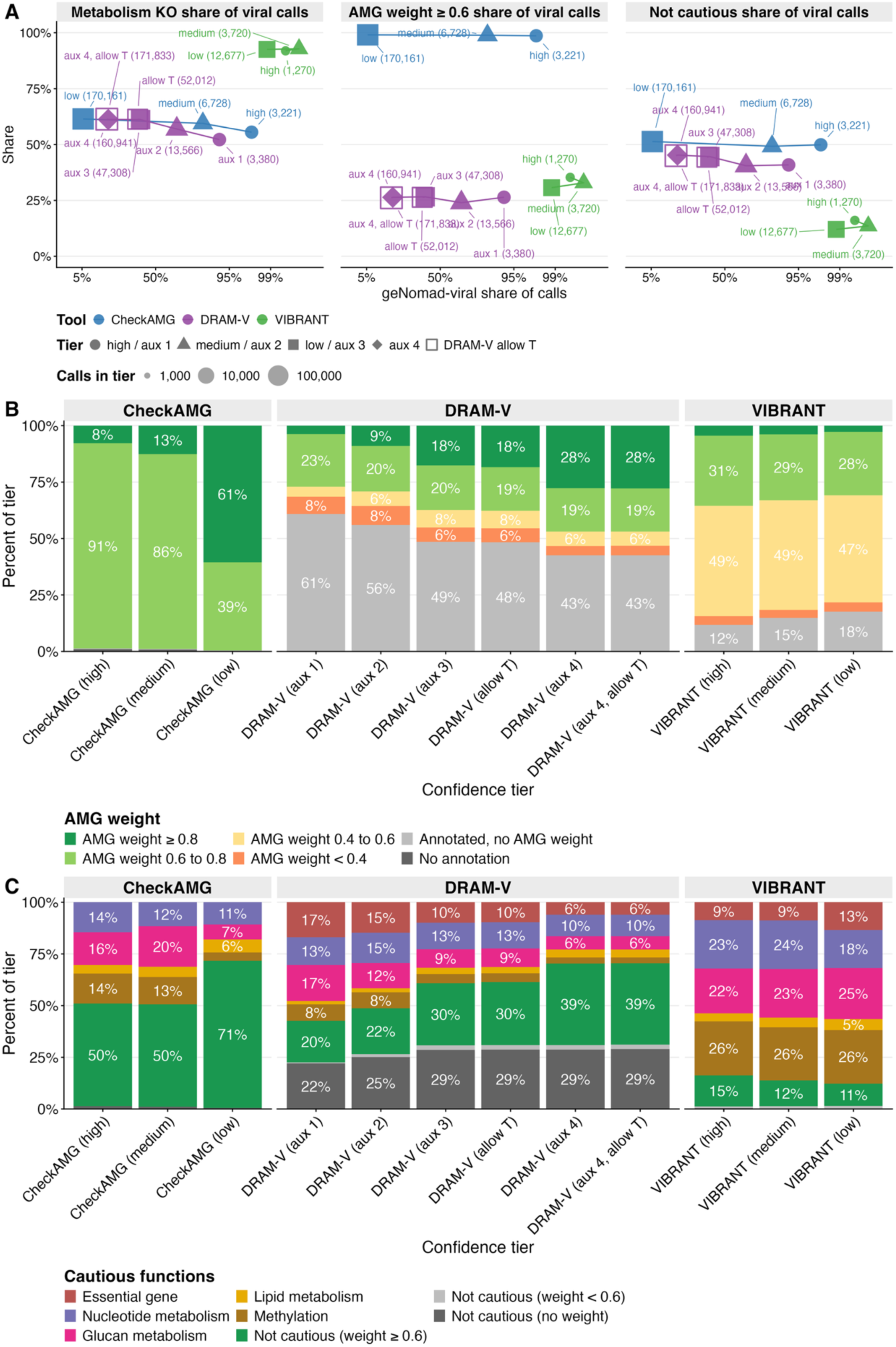
CheckAMG *annotate* produces confident AMG annotations across multiple tiers. **(A)** Share of each tier’s calls that are geNomad viral, on a logit axis, against three properties of those viral calls, the share with a Metabolism KO, with an AMG weight of at least 0.6, and with no cautious-function flag. Point area gives the calls in the tier on a log scale, shape the matched level and color the tool, and open squares mark the DRAM-V configurations that keep T-flagged genes. Viral calls with no annotation carry no cautious assignment and are not counted as not cautious. **(B)** AMG-weight composition of all calls at every tier for all three tools. **(C)** Cautious-function composition of every tool-by-tier combination. In every panel all tiers are cumulative, with DRAM-V (default) shown as DRAM-V (aux 1), (aux 2) and (aux 3) and DRAM-V (allow T), (aux 4) and (aux 4, allow T) shown in full, each including the default calls.

**Extended Data Figure 7.**
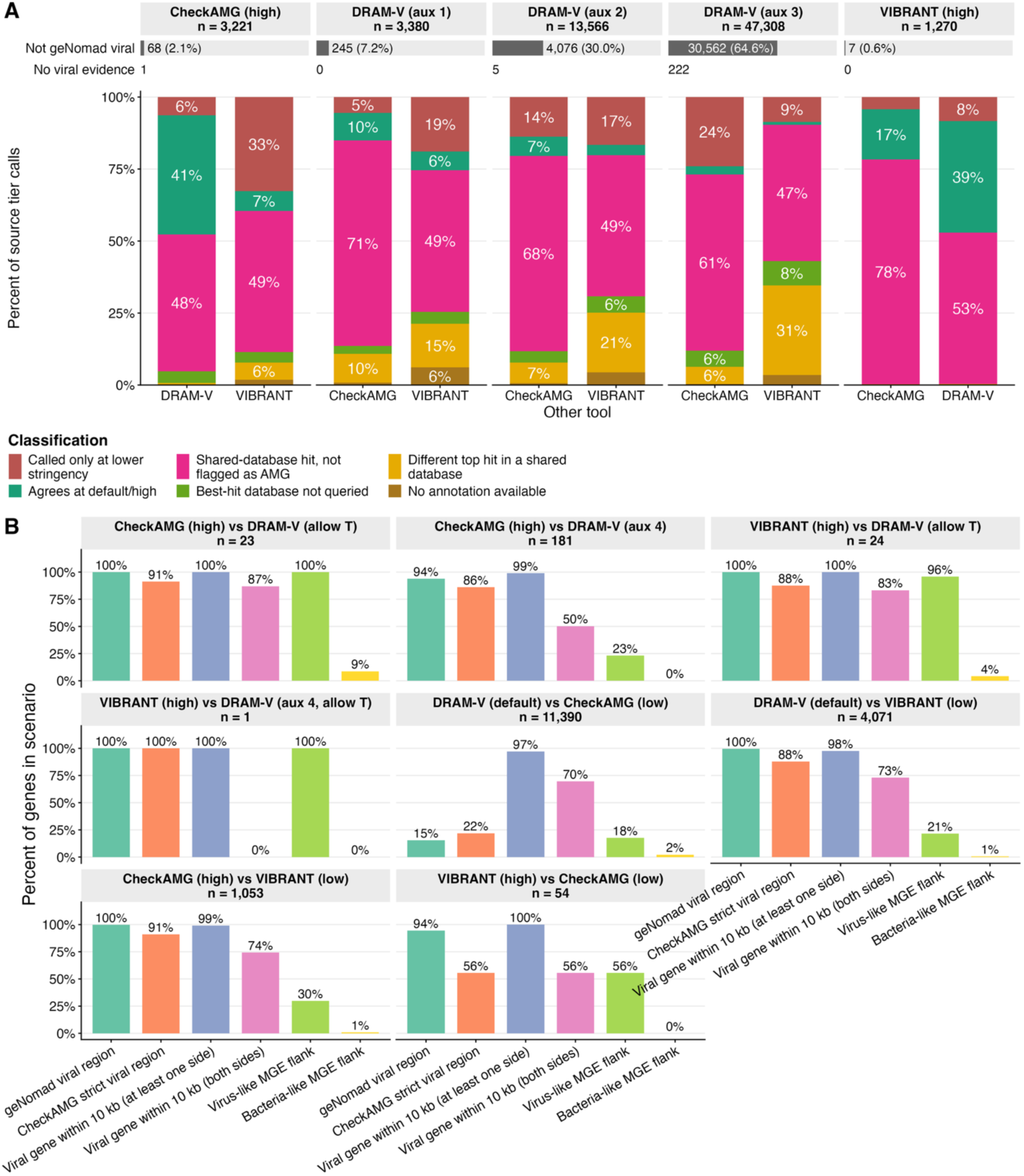
Reasons for non-overlapping AMG calls, and independent validation of viral origin. **(A)** For each source tier (CheckAMG high, DRAM-V (aux 1), (aux 2) and (aux 3), VIBRANT high), the fraction of genes falling into each of six reasons the other tools did not match the same gene: called only at a lower-stringency setting, agrees at default/high, shared-database hit not flagged as AMG, best-hit database not queried, different top hit in a shared database, or no annotation available. Each facet label gives the number of source-tier calls. The DRAM-V levels are cumulative, so each includes the calls of the stricter ones, and DRAM-V (aux 3) is DRAM-V (default). Calls not on a geNomad-viral region numbered 68 of 3,221 for CheckAMG (high) and 7 of 1,270 for VIBRANT (high), and 245 of 3,380, 4,076 of 13,566 and 30,562 of 47,308 at DRAM-V (aux 1), (aux 2) and (aux 3). Calls with no viral evidence numbered 1 for CheckAMG (high), 0 for VIBRANT (high), and 0, 5 and 222 at the three DRAM-V levels. For DRAM-V, a call only at a lower-stringency setting means the gene carried DRAM-V’s metabolism (M) flag together with an auxiliary score of 4 or the T flag. It does not indicate that DRAM-V found viral support for the gene or annotated the same function as the source tool. **(B)** For each tier-conflict scenario, in which a gene is called by a reference tool at its high/default tier but only cleared by the compared tool at a weaker setting, the percentage of genes located in (left to right): a geNomad viral region at FDR at most 0.1, a CheckAMG strict viral region, within 10 kb of a gene with V-score at least 10 on at least one side, on both sides, flanked by a virus-like MGE gene (nearest MGE V-score at least 10), or flanked by a bacterial-like MGE gene (nearest MGE V-score below 10). The two DRAM-V (default) scenarios exclude genes called at the other tool’s high tier. In each scenario the reference tool is the source of the call and the compared tool is the one being tested for why it did not make the same call at its default/high tier.

**Extended Data Figure 8.**
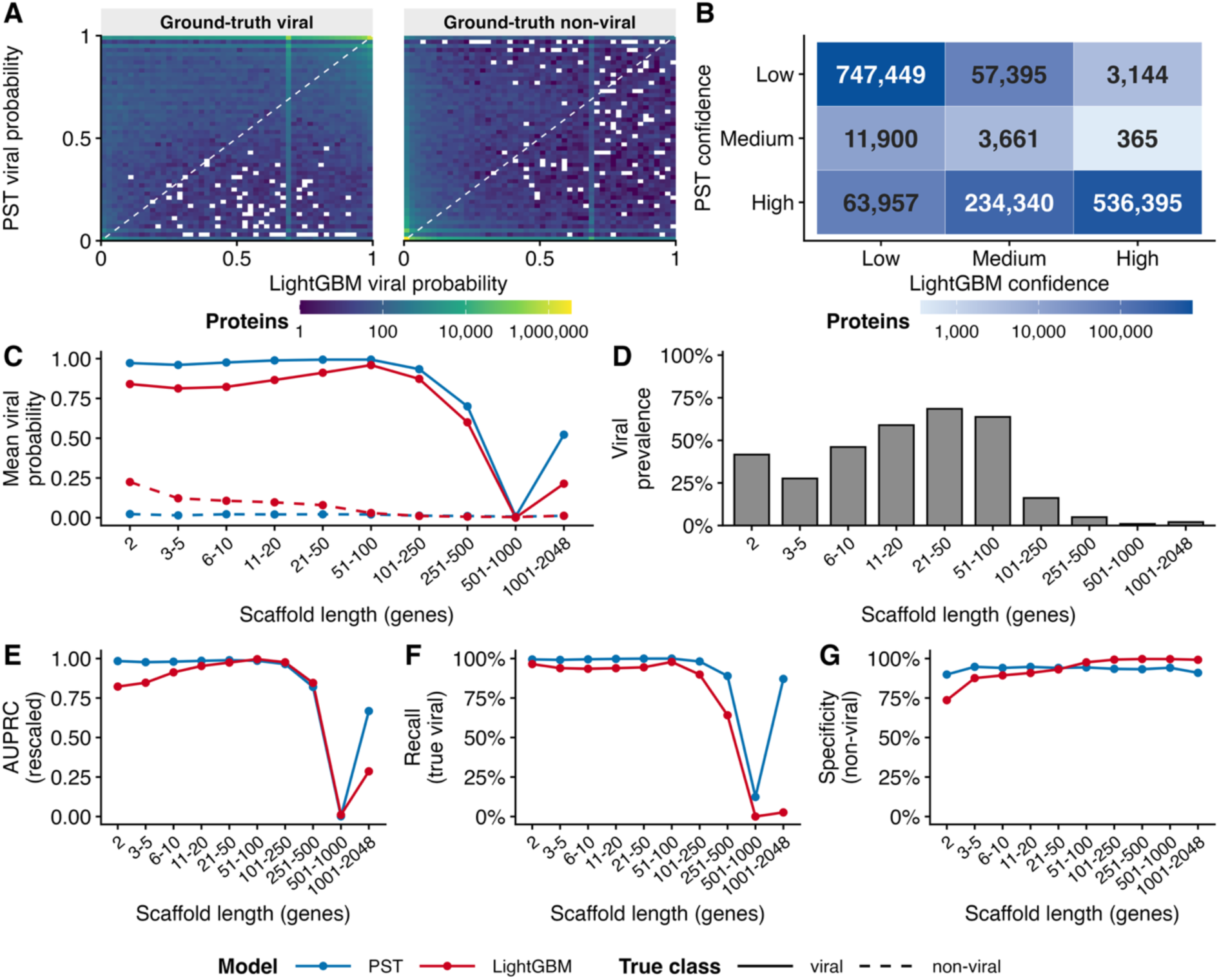
Comparison of the *annotate* (LightGBM) and *de-novo* (CheckAMG-PST) viral classifiers. **(A**) Joint distribution of per-protein viral probabilities from the two classifiers across all test datasets, binned in two dimensions (LightGBM on the x-axis, CheckAMG-PST on the y-axis) and colored by the number of proteins (log_10_ scale), faceted by ground-truth class (viral or non-viral); the dashed line marks equal probabilities. **(B)** Agreement of the discrete viral-origin confidence calls (low, medium, high) between the two classifiers, colored and labeled by the number of proteins in each combination. **(C)** Mean per-protein viral probability as a function of scaffold length (binned by number of genes) for each classifier (CheckAMG-PST, blue; LightGBM, red) and true class (viral, solid line; non-viral, dashed line). **(D)** Viral prevalence (fraction of proteins that are truly viral) in each scaffold-length bin. **(E–G)** Classifier performance within each scaffold-length bin for CheckAMG-PST (blue) and LightGBM (red): prevalence-rescaled Area Under the Precision-Recall Curve (AUPRC) (E), recall among true viral proteins at medium or higher confidence (F), and specificity among non-viral proteins at medium or higher confidence (G).

**Extended Data Figure 9.**
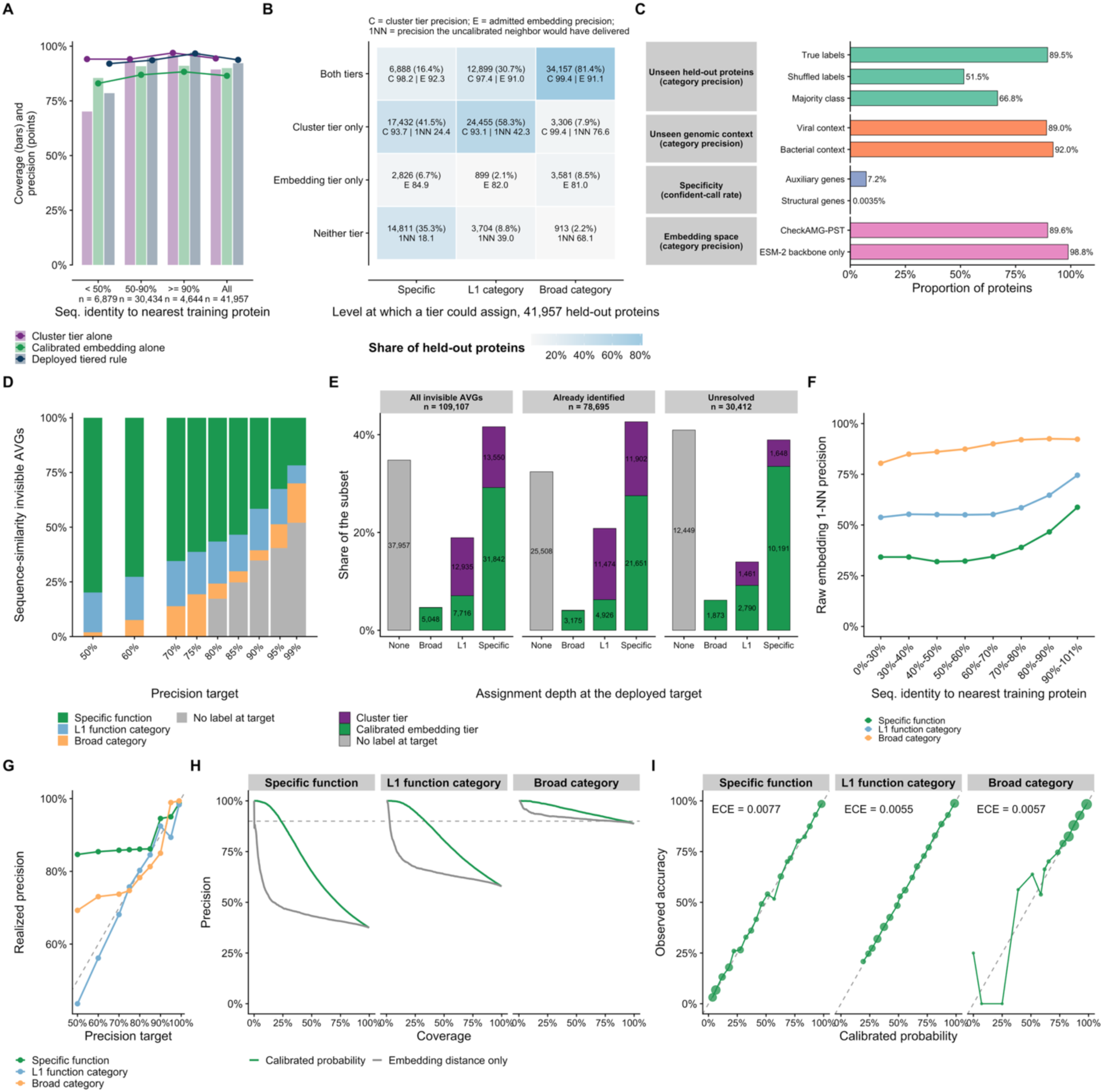
Validation of functional label propagation. **(A)** Functional propagation coverage (bars) and precision (points) for the cluster tier alone, the calibrated embedding tier alone, and the deployed tiered rule, by sequence identity to the nearest training protein. **(B)** The depth at which each propagation tier could assign a functional label, over 41,957 held-out proteins, giving the precision the sequence-cluster tier achieved (“C”), the precision of the admitted embedding labels (“E”), and the precision the uncalibrated nearest neighbor would have delivered (“1NN”). **(C)** Validation controls for functional propagation assignments, at the depth of broad-category depth. Propagation onto unseen held-out proteins compared against shuffled-label and majority-class baselines; proteins whose nearest labeled neighbor lies in a viral or a bacterial genome; the rate of confident calls on auxiliary genes versus structural genes as a negative control; and the finetuned CheckAMG-PST model vs. the unmodified ESM-2 base model at matched sample size. Bars show category precision, except for specificity, which shows confident-call rate as indicated in the panel label. **(D)** Depth of label assigned to the 109,107 sequence-similarity invisible auxiliary viral genes across multiple precision targets that were evaluated. **(E)** Depth of label assigned at the deployed precision target and the tier that supplied it, for all 109,107 sequence-similarity invisible auxiliary viral genes and separately for those already identified, meaning by DefenseFinder or by a structure with a known function, and for those left unresolved. Segments holding under 2% of a subset carry no printed count, which affects the 59 cluster-tier broad-category labels in each of the first two subsets and the none in the third. **(F)** Precision of the uncalibrated nearest-neighbor label as a function of sequence identity to the nearest training protein, at each depth. **(G)** Precision actually realized as the precision target is raised, at each label depth. The dashed line marks exact agreement between target and realized precision. **(H)** Precision versus coverage at each depth for held-out reference proteins, comparing the calibrated probability to ranking by raw embedding distance alone. The horizontal line marks 90% precision. **(I)** Calibration reliability for held-out reference proteins, measured as the observed fraction of correct labels among proteins grouped by their assigned calibrated probability, with point size proportional to group size. The expected calibration error (ECE) is given for each depth.

**Extended Data Figure 10.**
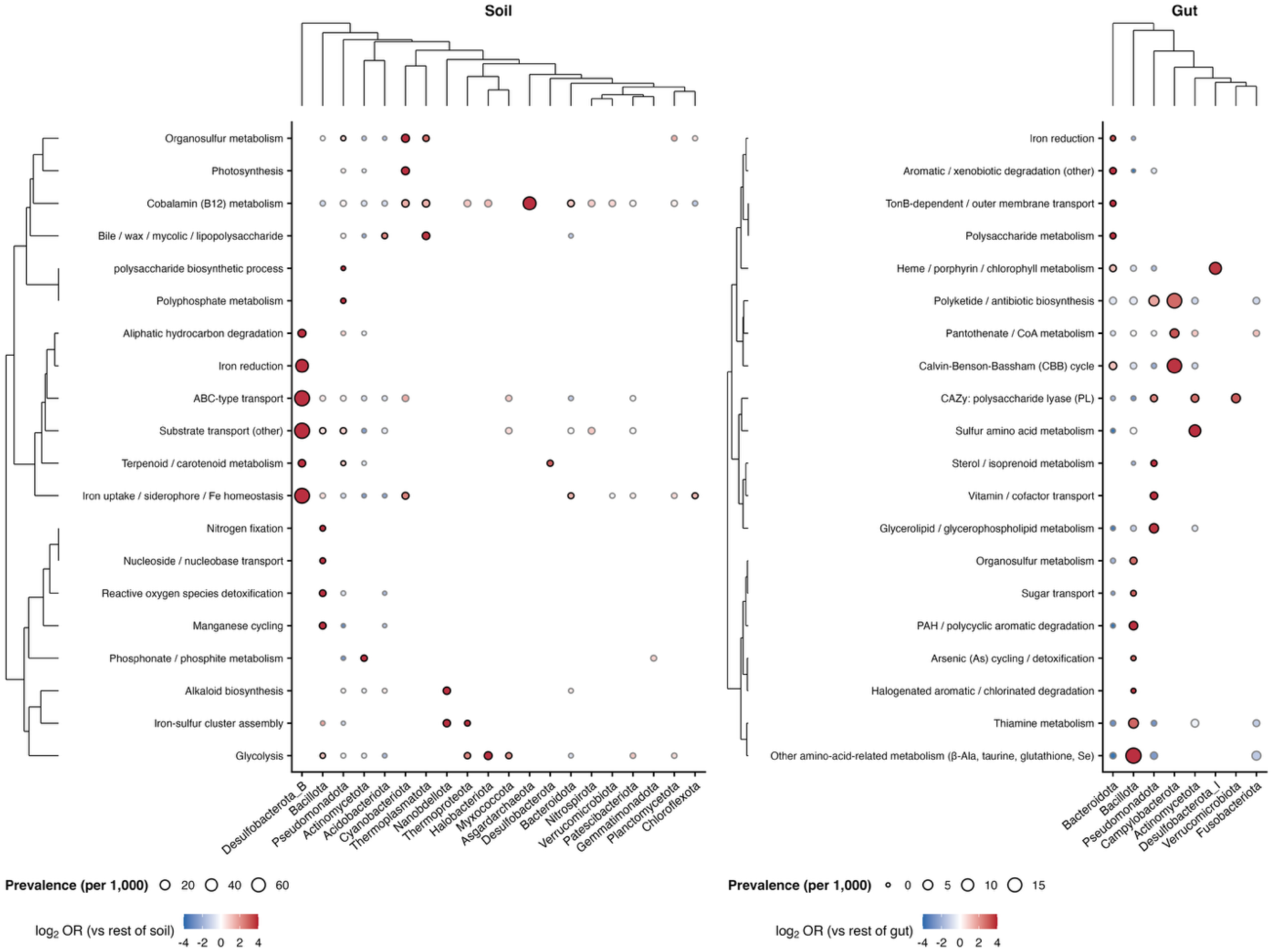
Host-phylum enrichment of CheckAMG-predicted AMG functions recapitulates known host biology. Auxiliary metabolic gene (AMG) functions significantly enriched within specific host phyla, tested separately in soil (left) and human-gut (right) viral genomes. A genome counts toward an AMG function (row) if the function is assigned by CheckAMG *annotate* or by a *de-novo* propagated specific function mapped to its L2-level category in the CheckAMG reference. Each point is one function (row)-phylum (column, iPHoP prediction from MetaVR metadata) comparison (two-sided Fisher’s exact test, Benjamini-Hochberg-adjusted *q* < 0.05, odds ratio > 1.5). Shown are the 20 functions with the largest maximum odds ratio per biome among those significantly enriched in at least one phylum, for simplicity. Point fill corresponds to the log_2_ odds ratio of the function’s prevalence in that host-phylum relative to the rest of the biome, and point size shows prevalence per 1,000 genomes in the phylum. Rows and columns are ordered by hierarchical clustering of the log_2_ odds ratio matrix.

## References

1. Martin, C., Emerson, J. B., Roux, S. & Anantharaman, K. A call for caution in the biological interpretation of viral auxiliary metabolic genes. Nature Microbiology 10, 2122–2129 (2025).

2. Lindell, D. et al. Transfer of photosynthesis genes to and from Prochlorococcus viruses. Proceedings of the National Academy of Sciences 101, 11013–11018 (2004).

3. Breitbart, M., Thompson, L., Suttle, C. & Sullivan, M. Exploring the Vast Diversity of Marine Viruses. Oceanography 20, 135–139 (2007).

4. Breitbart, M., Bonnain, C., Malki, K. & Sawaya, N. A. Phage puppet masters of the marine microbial realm. Nature Microbiology 3, 754–766 (2018).

5. Forterre, P. Manipulation of cellular syntheses and the nature of viruses: The virocell concept. in Comptes Rendus Chimie vol. 14 392–399 (Elsevier Masson SAS, 2011).

6. Urvoy, M., Baumgart, L., Howard-Varona, C. & Sullivan, M. B. Beyond AMGs: Phage-encoded transcription and sigma factors as understudied virocell reprogramming tools. Trends in Microbiology 10.1016/j.tim.2026.05.019 (2026) doi:10.1016/j.tim.2026.05.019.

7. Bragg, J. G. & Chisholm, S. W. Modeling the Fitness Consequences of a Cyanophage-Encoded Photosynthesis Gene. PLOS ONE 3, e3550-(2008).

8. Thompson, L. R. et al. Phage auxiliary metabolic genes and the redirection of cyanobacterial host carbon metabolism. Proceedings of the National Academy of Sciences 108, E757–E764 (2011).

9. Zimmerman, A. E. et al. Metabolic and biogeochemical consequences of viral infection in aquatic ecosystems. Nature Reviews Microbiology 18, 21–34 (2020).

10. Bouvier, T. & Del Giorgio, P. A. Key role of selective viral-induced mortality in determining marine bacterial community composition. Environmental Microbiology 9, 287–297 (2007).

11. Hennes K P, Suttle C A, & Chan A M. Fluorescently Labeled Virus Probes Show that Natural Virus Populations Can Control the Structure of Marine Microbial Communities. Applied and Environmental Microbiology 61, 3623–3627 (1995).

12. Mann, N. H., Cook, A., Millard, A., Bailey, S. & Clokie, M. Bacterial photosynthesis genes in a virus. Nature 424, 741 (2003).

13. Lindell, D., Jaffe, J. D., Johnson, Z. I., Church, G. M. & Chisholm, S. W. Photosynthesis genes in marine viruses yield proteins during host infection. Nature 438, 86–89 (2005).

14. Sullivan, M. B. et al. Prevalence and Evolution of Core Photosystem II Genes in Marine Cyanobacterial Viruses and Their Hosts. PLOS Biology 4, e234-(2006).

15. Seed, K. D., Lazinski, D. W., Calderwood, S. B. & Camilli, A. A bacteriophage encodes its own CRISPR/Cas adaptive response to evade host innate immunity. Nature 494, 489–491 (2013).

16. George, N. A. & Hug, L. A. CRISPR-resolved virus-host interactions in a municipal landfill include non-specific viruses, hyper-targeted viral populations, and interviral conflicts. Scientific Reports 13, 5611 (2023).

17. Al-Shayeb, B., et al. Diverse virus-encoded CRISPR-Cas systems include streamlined genome editors. Cell 185, 4574–4586.e16 (2022).

18. Schwartz D. A., Lehmkuhl B. K., & Lennon J. T. Phage-Encoded Sigma Factors Alter Bacterial Dormancy. mSphere 7, e00297–22 (2022).

19. Fiamenghi, M. B. et al. Meta-virus resource (MetaVR): expanding the frontiers of viral diversity with 24 million uncultivated virus genomes. Nucleic Acids Research 54, D801– D812 (2026).

20. Kosmopoulos, J. C. & Anantharaman, K. Viral Dark Matter: Illuminating Protein Function, Ecology, and Biotechnological Promises. Biochemistry 64, 4609–4627 (2025).

21. Tian, F. et al. Prokaryotic-virus-encoded auxiliary metabolic genes throughout the global oceans. Microbiome 12, 159 (2024).

22. Nayfach, S. et al. CheckV assesses the quality and completeness of metagenome-assembled viral genomes. Nature Biotechnology 39, 578–585 (2021).

23. Garneau, J. R. et al. High-throughput identification of viral termini and packaging mechanisms in virome datasets using PhageTermVirome. Scientific Reports 11, 18319 (2021).

24. Khedkar, S. et al. Landscape of mobile genetic elements and their antibiotic resistance cargo in prokaryotic genomes. Nucleic Acids Research 50, 3155–3168 (2022).

25. Roux, S., Emerson, J. B., Eloe-Fadrosh, E. A. & Sullivan, M. B. Benchmarking viromics: an *in silico* evaluation of metagenome-enabled estimates of viral community composition and diversity. PeerJ 5, e3817 (2017).

26. Pratama, A. A. et al. Expanding standards in viromics: in silico evaluation of dsDNA viral genome identification, classification, and auxiliary metabolic gene curation. PeerJ 9, e11447 (2021).

27. Shaffer, M. et al. DRAM for distilling microbial metabolism to automate the curation of microbiome function. Nucleic Acids Research 48, 8883–8900 (2020).

28. Kieft, K., Zhou, Z. & Anantharaman, K. VIBRANT: automated recovery, annotation and curation of microbial viruses, and evaluation of viral community function from genomic sequences. Microbiome 8, 90 (2020).

29. Emerson, J. B. et al. Host-linked soil viral ecology along a permafrost thaw gradient. Nature Microbiology 3, 870–880 (2018).

30. Krishnamurthy, S. R. & Wang, D. Origins and challenges of viral dark matter. Virus Research 239, 136–142 (2017).

31. Lin, Z. et al. Evolutionary-scale prediction of atomic-level protein structure with a language model. Science 379, 1123–1130 (2023).

32. Mordret, E. et al. Protein and genomic language models uncover the unexplored diversity of bacterial immunity. Science 392, eadv8275 (2026).

33. Zhou, K., Kosmopoulos, J. C., Colón, E. D., Badciong, P. J. & Anantharaman, K. V- and VL-scores unveil viral signatures and origins of protein families. Nature Communications 10.1038/s41467-026-72028-0 (2026) doi:10.1038/s41467-026-72028-0.

34. Martin, C., Gitter, A. & Anantharaman, K. Protein Set Transformer: a protein-based genome language model to power high-diversity viromics. Nature Communications 16, 11123 (2025).

35. Ke, G. et al. LightGBM: A Highly Efficient Gradient Boosting Decision Tree. in Advances in Neural Information Processing Systems (eds Guyon, I. et al.) vol. 30 (Curran Associates, Inc., 2017).

36. Kanehisa, M., Sato, Y., Kawashima, M., Furumichi, M. & Tanabe, M. KEGG as a reference resource for gene and protein annotation. Nucleic Acids Research 44, D457–D462 (2016).

37. Mistry, J. et al. Pfam: The protein families database in 2021. Nucleic Acids Research 49, D412–D419 (2021).

38. Terzian, P., et al. PHROG: families of prokaryotic virus proteins clustered using remote homology. NAR Genomics and Bioinformatics 3, lqab067 (2021).

39. Camargo, A. P. et al. Identification of mobile genetic elements with geNomad. Nature Biotechnology 1303–1312 (2023) doi:10.1038/s41587-023-01953-y.

40. Fullam, A. et al. proGenomes3: approaching one million accurately and consistently annotated high-quality prokaryotic genomes. Nucleic Acids Research 51, D760–D766 (2022).

41. Johansson, M. H. K., Aarestrup, F. M. & Petersen, T. N. Importance of mobile genetic elements for dissemination of antimicrobial resistance in metagenomic sewage samples across the world. PLOS ONE 18, e0293169 (2023).

42. Zheng, J., et al. dbCAN3: automated carbohydrate-active enzyme and substrate annotation. Nucleic Acids Research 51, W115–W121 (2023).

43. Prestat, E. et al. FOAM (Functional Ontology Assignments for Metagenomes): a Hidden Markov Model (HMM) database with environmental focus. Nucleic Acids Research 42, e145–e145 (2014).

44. Zhou, Z. et al. METABOLIC: high-throughput profiling of microbial genomes for functional traits, metabolism, biogeochemistry, and community-scale functional networks. Microbiome 10, 33 (2022).

45. McGivern, B. B. et al. Microbial polyphenol metabolism is part of the thawing permafrost carbon cycle. Nature Microbiology 9, 1454–1466 (2024).

46. The Gene Ontology Consortium. The Gene Ontology knowledgebase in 2026. Nucleic Acids Research 54, D1779–D1792 (2026).

47. Kanehisa, M. et al. KEGG for linking genomes to life and the environment. Nucleic Acids Research 36, D480–D484 (2008).

48. Aramaki, T. et al. KofamKOALA: KEGG Ortholog assignment based on profile HMM and adaptive score threshold. Bioinformatics 36, 2251–2252 (2020).

49. Guo, J. et al. VirSorter2: a multi-classifier, expert-guided approach to detect diverse DNA and RNA viruses. Microbiome 9, 37 (2021).

50. Graham, E. B. et al. A global atlas of soil viruses reveals unexplored biodiversity and potential biogeochemical impacts. Nature Microbiology 9, 1873–1883 (2024).

51. Tesson, F. et al. Systematic and quantitative view of the antiviral arsenal of prokaryotes. Nature Communications 13, 2561 (2022).

52. Bush, M. J. The actinobacterial WhiB-like (Wbl) family of transcription factors. Molecular Microbiology 110, 663–676 (2018).

53. Bei, Q. et al. Extreme summers impact cropland and grassland soil microbiomes. The ISME Journal 17, 1589–1600 (2023).

54. Ibarra, J. A., Pérez-Rueda, E., Carroll, R. K. & Shaw, L. N. Global analysis of transcriptional regulators in Staphylococcus aureus. BMC Genomics 14, 126 (2013).

55. Sharma Vikas, Hardy Aël, Luthe Tom, & Frunzke Julia. Phylogenetic Distribution of WhiB- and Lsr2-Type Regulators in Actinobacteriophage Genomes. Microbiology Spectrum 9, e00727–21 (2021).

56. Rybniker, J. et al. Insights into the function of the WhiB-like protein of mycobacteriophage TM4 – a transcriptional inhibitor of WhiB2. Molecular Microbiology 77, 642–657 (2010).

57. Jamet, A. et al. A widespread family of polymorphic toxins encoded by temperate phages. BMC Biology 15, 75 (2017).

58. Yutin, N. et al. Discovery of an expansive bacteriophage family that includes the most abundant viruses from the human gut. Nature Microbiology 3, 38–46 (2018).

59. Docherty, J. A. D., et al. Diverse defense systems and prophages in human-associated Bifidobacterium species reveal coevolutionary “arms race” dynamics. Cell Reports 44, (2025).

60. Huang, X. et al. Genome-resolved metagenomics reveal soil and viral drivers of keystone bacterial traits shaping nutrient cycling and soybean yield across agroecosystems. Soil Ecology Letters 7, 250346 (2025).

61. Trubl, G., et al. Soil Viruses Are Underexplored Players in Ecosystem Carbon Processing. mSystems 3, (2018).

62. Huang, X. et al. Soil nutrient conditions alter viral lifestyle strategy and potential function in phosphorous and nitrogen metabolisms. Soil Biology and Biochemistry 189, 109279 (2024).

63. Crits-Christoph, A., Diamond, S., Butterfield, C. N., Thomas, B. C. & Banfield, J. F. Novel soil bacteria possess diverse genes for secondary metabolite biosynthesis. Nature 558, 440–444 (2018).

64. Shaw, D. R. et al. Independently evolved extracellular electron transfer pathways in ecologically diverse Desulfobacterota. The ISME Journal 19, wraf097 (2025).

65. Kieft, K. et al. Virus-associated organosulfur metabolism in human and environmental systems. Cell Reports 36, 109471 (2021).

66. Wolf, P. G. et al. Diversity and distribution of sulfur metabolic genes in the human gut microbiome and their association with colorectal cancer. Microbiome 10, 64 (2022).

67. Zhong, Z.-P. et al. Viral potential to modulate microbial methane metabolism varies by habitat. Nature Communications 15, 1857 (2024).

68. Zhou, Z. et al. Unravelling viral ecology and evolution over 20 years in a freshwater lake. Nature Microbiology 10, 231–245 (2025).

69. Kieft, K. et al. Ecology of inorganic sulfur auxiliary metabolism in widespread bacteriophages. Nature Communications 12, 3503 (2021).

70. Anantharaman, K. et al. Sulfur Oxidation Genes in Diverse Deep-Sea Viruses. Science 344, 757–760 (2014).

71. Langwig, M. V. et al. Endemism shapes viral ecology and evolution in globally distributed hydrothermal vent ecosystems. Nature Communications 16, 4076 (2025).

72. Kosmopoulos, J. C., Pallier, W., Malik, A. A. & Anantharaman, K. Ecosystem health shapes viral ecology in peatland soils. Nature Microbiology 11, 142–154 (2026).

73. Wu, R. et al. Structural characterization of a soil viral auxiliary metabolic gene product – a functional chitosanase. Nature Communications 13, 5485 (2022).

74. Bouras, G. et al. Protein structure-informed bacteriophage genome annotation with Phold. Nucleic Acids Research 54, gkaf1448 (2026).

75. Jha, N. et al. Gaia: An AI-enabled genomic context–aware platform for protein sequence annotation. Science Advances 11, eadv5109 (2025).

76. Larralde, M. Pyrodigal: Python bindings and interface to Prodigal, an efficient method for gene prediction in prokaryotes. Journal of Open Source Software 7, 4296 (2022).

77. Hyatt, D. et al. Prodigal: prokaryotic gene recognition and translation initiation site identification. BMC Bioinformatics 11, 119 (2010).

78. Eddy, S. R. Accelerated Profile HMM Searches. PLoS Computational Biology 7, e1002195 (2011).

79. Larralde, M. & Zeller, G. PyHMMER: a Python library binding to HMMER for efficient sequence analysis. Bioinformatics 39, (2023).

80. Eren, A. M. et al. Community-led, integrated, reproducible multi-omics with anvi’o. Nature Microbiology 6, 3–6 (2020).

81. Kananen, K. et al. Adaptive adjustment of profile HMM significance thresholds improves functional and metabolic insights into microbial genomes. Bioinformatics Advances 5, vbaf039 (2025).

82. Figueroa III, J. L., Dhungel, E., Bellanger, M., Brouwer, C. R. & White III, R. A. MetaCerberus: distributed highly parallelized HMM-based processing for robust functional annotation across the tree of life. Bioinformatics 40, btae119 (2024).

83. Mitchell, A. et al. The InterPro protein families database: the classification resource after 15 years. Nucleic Acids Research 43, D213–D221 (2014).

84. Li, W. et al. RefSeq: expanding the Prokaryotic Genome Annotation Pipeline reach with protein family model curation. Nucleic Acids Research 49, D1020–D1028 (2020).

85. Roux, S. et al. Ecology and evolution of viruses infecting uncultivated SUP05 bacteria as revealed by single-cell- and meta-genomics. eLife 3, e03125 (2014).

86. Steinegger, M. & Söding, J. MMseqs2 enables sensitive protein sequence searching for the analysis of massive data sets. Nature Biotechnology 35, 1026–1028 (2017).

87. Van Dongen, S. Graph Clustering Via a Discrete Uncoupling Process. SIAM Journal on Matrix Analysis and Applications 30, 121–141 (2008).

88. Shi, Y., Ke, G., Chen, Z., Zheng, S. & Liu, T.-Y. Quantized Training of Gradient Boosting Decision Trees. in Advances in Neural Information Processing Systems (eds Koyejo, S. et al.) vol. 35 18822–18833 (Curran Associates, Inc., 2022).

89. Head, T., Kumar, M., Nahrstaedt, H., Louppe, G. & Shcherbatyi, I. scikit-optimize.

90. Buitinck, L. et al. API design for machine learning software: experiences from the scikit-learn project. in ECML PKDD Workshop: Languages for Data Mining and Machine Learning 108– 122 (2013).

91. Shkoporov, A. N. et al. The Human Gut Virome Is Highly Diverse, Stable, and Individual Specific. Cell Host and Microbe 26, 527–541.e5 (2019).

92. Tran, P. Q., Bachand, S. C., Peterson, B., He, S. & Anantharaman, K. Viral impacts on microbial activity and biogeochemical cycling in a seasonally anoxic freshwater lake. bioRxiv 10.1101/2023.04.19.537559 (2023) doi:10.1101/2023.04.19.537559.

93. Hazard, C. Soil microbial communities from microcosms at the University of Lyon, Ecully, France. JGI IMG/M 10.46936/10.25585/60008876 (2024).

94. Camargo, A. P. et al. IMG/VR v4: an expanded database of uncultivated virus genomes within a framework of extensive functional, taxonomic, and ecological metadata. Nucleic Acids Research 51, D733–D743 (2023).

95. Mölder, F. et al. Sustainable data analysis with Snakemake. F1000Research 10, 33 (2021).

96. Center for High Throughput Computing. Center for High Throughput Computing. (2006) doi:10.21231/GNT1-HW21.

97. Johnson, J., Douze, M. & Jégou, H. Billion-scale similarity search with GPUs. IEEE Transactions on Big Data 7, 535–547 (2019).

98. Traag, V. A., Waltman, L. & van Eck, N. J. From Louvain to Leiden: guaranteeing well-connected communities. Scientific Reports 9, 5233 (2019).

99. Csárdi, G. & Nepusz, T. The igraph software package for complex network research. InterJournal **Complex Systems**, 1695 (2006).

100. Falcon, W. & The PyTorch Lightning Team. PyTorch Lightning. 10.5281/zenodo.3530844.

101. Roux, S. et al. iPHoP: An integrated machine learning framework to maximize host prediction for metagenome-derived viruses of archaea and bacteria. PLOS Biology 21, e3002083 (2023).

102. Heinzinger, M., et al. Bilingual language model for protein sequence and structure. NAR Genomics and Bioinformatics 6, lqae150 (2024).

103. Kim, W. et al. Rapid and sensitive protein complex alignment with Foldseek-Multimer. Nature Methods 22, 469–472 (2025).

104. Kim, R. S., Levy Karin, E., Mirdita, M., Chikhi, R. & Steinegger, M. BFVD—a large repository of predicted viral protein structures. Nucleic Acids Research 53, D340–D347 (2025).

105. Barrio-Hernandez, I. et al. Clustering predicted structures at the scale of the known protein universe. Nature 622, 637–645 (2023).

