## Supplemental Text for "CheckAMG: Accurate Identification of Auxiliary Viral Genes with Genome Language Models"

### Supplementary Information

#### Supplemental Results

##### AMG call disagreement between CheckAMG, DRAM-V, and VIBRANT

*Most disagreements reflect curation and reference differences, not homology detection*

For every gene called as an AMG at default/high by a reference tool, we classified each compared tool's response into one of six mutually exclusive categories (Extended Data Figure 7A; Methods). Of the 3,221 AMG calls produced by CheckAMG (high), VIBRANT most often produced a hit to the same database entry but did not flag it as an AMG (49.0%) or called the gene only at a lower-confidence tier (32.7%), with smaller shares explained by a different top hit in a shared database (5.9%), a best-hit database VIBRANT does not query (3.7%), or no hit at all (1.8%). DRAM-V flagged 6.3% of the same genes only at its lower-stringency settings. For the 1,270 VIBRANT (high) AMG calls, CheckAMG agreed at its default/high confidence tier 17.4% of the time, but most often produced a hit to the same database entry without flagging it as an AMG (78.0%), and the remainder were called only at a lower-confidence tier (4.3%) or matched a different top hit in a shared database (0.3%). DRAM-V likewise flagged 8.3% of the VIBRANT AMGs only at its lower-stringency settings, and another 52.5% shared an annotation identifier with DRAM-V without being flagged. These patterns show that the disagreement among tools is driven by differences in curation and reference composition rather than by differences in the ability to detect homology.

*Most DRAM-V calls that CheckAMG does not make are not geNomad viral*

The most common reason neither CheckAMG nor VIBRANT matched a DRAM-V (default) call was that they had a hit to the same database entry without flagging it as an AMG (61.2% of the 47,308 DRAM-V (default) calls for CheckAMG, 47.3% for VIBRANT, pooled across the auxiliary scores in Extended Data Figure 7A), and few such calls had no annotation at all (0.37% and 3.5%, respectively). Of the 47,308 genes called by DRAM-V (default), 15,980 (33.8%) are represented by a KEGG annotation and 26,823 (56.7%) by a Pfam annotation (Extended Data Figure 5C, Supplemental Table 8, Supplemental Data 3).

*CheckAMG's high-confidence calls have strong independent evidence of viral origin*

Because DRAM-V flagged 204 of the genes CheckAMG calls with high confidence only at its lower-stringency settings, we asked whether those CheckAMG calls have support for viral origin. Each conflicted gene was cross-referenced against four genomic-context signals: geNomad viral classification at FDR at most 0.1, inclusion in a CheckAMG strict viral region, a gene with V-score at least 10 within 10 kb on one or both sides, and the character of the nearest MGE-annotated flank (Extended Data Figure 7B; Supplemental Table 9). These signals are converging rather than fully independent, because the strict-region and context V-score signals both derive from the annotation-based V-scores used by CheckAMG; geNomad classification is the one fully external signal.

When CheckAMG (high) calls were cleared by DRAM-V only at auxiliary score of 4 (n = 181), 94% lay on geNomad viral regions, 86% inside a CheckAMG strict viral region, 99% had a V-score-at-least-10 viral gene within 10 kb on at least one side, and 50% on both sides, consistent with these genes being virus-encoded. When CheckAMG (high) calls were cleared by DRAM-V only with the T flag allowed (n = 23), viral evidence remained high (100% on geNomad viral regions, 91% inside a strict viral region, 100%/87% with a near-viral gene on one/both sides), and 100% were flanked by a virus-like MGE gene, defined as a neighboring gene with a V-score of at least 10, meaning it is commonly encoded by viruses such as an integrase, recombinase, or transposase of a transposable phage, rather than a non-viral transposon. The T flag is not itself evidence of a transposon near these genes, because DRAM-V v1.5.0 assigns it to every gene on a scaffold on which any gene has a Pfam hit to one of 17 transposase families. Similarly, 77 of the 204 genes carried the B flag, which DRAM-V assigns to any three consecutive genes with its metabolism flag and which raises their auxiliary score to 4.

*Both comparison tools differ from CheckAMG for identifiable reasons*

The two tools differ from CheckAMG for distinct, well-defined reasons. VIBRANT is limited by its reference: the identifiers CheckAMG uses for confident calls are largely absent from VIBRANT's curated set, so no parameter change could recover them. DRAM-V is limited by its default stringency, which pools three auxiliary scores, so most of its calls are on sequences geNomad does not classify as viral. CheckAMG integrates protein viral-origin confidence, AMG weight, and genomic context into a single scoring framework.

#### Biome-associated auxiliary metabolic functions in soil and human-gut viruses

*Biome-level enrichment of AMG biogeochemical categories*

For each AMG biogeochemical category we compared the proportion of soil versus gut viral genomes encoding it (Cochran-Mantel-Haenszel test stratified by quintiles of total gene count, Benjamini-Hochberg [BH]-adjusted, $q < 0.05$, $|\log2 \text{ odds ratio}| > 0.585$; Methods). Of 98 testable categories, 22 were soil-enriched, 42 were gut-enriched, and 34 showed no biome difference (Figure 6B; Supplemental Table 14).

Soil-enriched functions spanned phosphorus, carbohydrate, lipid, amino-acid, and secondary-metabolite categories. Phosphorus metabolism showed the strongest and most exclusive enrichment: phosphonate and phosphite metabolism (578 soil genomes versus 0 gut; odds ratio 93.1 after continuity correction; q = 2.0e-56), polyphosphate metabolism (84 versus 1; odds ratio 42.7; q = 1.1e-9), and phosphatase and phosphate liberation (odds ratio 2.1; q = 7.6e-16). Carbohydrate-active functions were broadly soil-enriched, consistent with the degradation of complex, plant-derived polysaccharides: auxiliary-activity/LPMO CAZymes (51 soil genomes versus 0 gut; odds ratio 8.6 after continuity correction; q = 6.0e-6), other carbohydrate-metabolisms (odds ratio 6.5; q = 8.8e-42), polysaccharide lyases (odds ratio 3.3; q = 7.1e-29), gluconeogenesis (odds ratio 2.6; q = 8.4e-43), general polysaccharide and complex-carbon degradation (odds ratio 2.3; q = 2.8e-146), and sugar interconversion (odds ratio 2.2; q = 5.0e-138). Lipid and fatty-acid functions were also soil-enriched, including a lipid-metabolism (other) category (odds ratio 10.5; q = 1.9e-159), bile/wax/mycolic-acid/lipopolysaccharide metabolism (odds ratio 2.9; q = 3.3e-12), fatty-acid biosynthesis (odds ratio 1.7; q = 5.5e-39), and glycerolipid/glycerophospholipid metabolism (odds ratio 1.7; q = 4.4e-10). Amino-acid functions comprised aminotransferase and transamination reactions (odds ratio 5.2; q = 5.4e-236), D-amino-acid metabolism (odds ratio 4.3; q = 1.1e-14), and branched-chain amino-acid metabolism (odds ratio 1.6; q = 2.4e-5). The remaining soil-enriched functions were an oxygenase/monooxygenase/dioxygenase category (odds ratio 16.8; q = 1.8e-7), polyketide and antibiotic biosynthesis (odds ratio 4.4; q = 9.6e-259), nucleoside and nucleobase transport (odds ratio 3.2; q = 1.1e-2), halogenated-aromatic and chlorinated compound degradation (odds ratio 3.0; q = 7.0e-5), pantothenate and coenzyme A metabolism (odds ratio 2.6; q = 3.0e-16), and denitrification (odds ratio 2.0; q = 1.5e-20). These functions are consistent with phosphorus cycling, secondary metabolism, and the breakdown of plant-derived organic matter that is commonly reported among microbial metabolisms in soil.

The gut-enriched set was larger and dominated by nutrient-transport and cofactor functions that track the gut nutritional environment. Substrate-transport categories were the single largest group: vitamin and cofactor transport (0 soil genomes versus 54 gut; gut:soil odds ratio 118.7 after continuity correction; q = 1.1e-23), TonB-dependent/outer-membrane transport (odds ratio 30.8; q = 1.7e-22), sugar transport (odds ratio 7.2; q = 7.2e-16), ABC transporters (odds ratio 6.8; q = 1.3e-33), a substrate-transport (other) category (odds ratio 6.1; q < 1e-300), inorganic-ion transport (odds ratio 5.1; q = 1.1e-24), phosphotransferase (PTS) system (odds ratio 4.8; q = 4.2e-22), major facilitator superfamily (MFS) transport (odds ratio 3.8; q = 6.8e-39), and ABC-type transport (odds ratio 3.5; q = 3.2e-134). Cofactor and vitamin metabolism was similarly enriched: thiamine metabolism (odds ratio 17.6; q = 5.2e-243), pyridoxal (B6) metabolism (odds ratio 4.6; q = 4.9e-14), NAD(P)/nicotinamide metabolism (odds ratio 3.9; q = 8.9e-245), thiol-cofactor metabolism (odds ratio 1.8; q = 4.4e-2), and riboflavin/FAD metabolism (odds ratio 1.7; q = 1.9e-6). Sulfur- and amino-acid-related functions followed the same pattern: sulfur-amino-acid metabolism (odds ratio 36.9; q = 1.8e-96), cysteine/methionine metabolism (odds ratio 10.1; q < 1e-300), a broader amino-acid category spanning beta-alanine, taurine, glutathione, and selenocompound metabolism (odds ratio 8.6; q < 1e-300), lysine metabolism (odds ratio 3.5; q = 6.1e-80), alanine/aspartate/glutamate metabolism (odds ratio 3.3; q = 1.8e-146), and organosulfur metabolism (odds ratio 2.2; q = 1.1e-12).

The remaining gut-enriched functions were more heterogeneous. Central-carbon and one-carbon pathways included the reductive TCA cycle (1 soil genome versus 65 gut; odds ratio 130.8; q = 1.6e-31), formaldehyde oxidation and fixation (odds ratio 11.0; q = 4.1e-124), the Wood-Ljungdahl pathway and homoacetogenesis (odds ratio 3.1; q = 1.9e-4), the oxidative TCA cycle (odds ratio 2.3; q = 1.2e-12), glycolysis/the Embden-Meyerhof-Parnas pathway (odds ratio 1.9; q = 1.0e-68), the 3-hydroxypropionate/4-hydroxybutyrate cycle (odds ratio 1.7; q = 2.2e-7), the Calvin-Benson-Bassham cycle (odds ratio 1.5; q = 1.4e-7), and pyruvate metabolism (odds ratio 1.5; q = 4.6e-13), alongside polysaccharide metabolism (odds ratio 19.8; q = 6.2e-21), polysaccharide biosynthesis (odds ratio 2.4; q = 2.7e-2), and glyoxylate/dicarboxylate metabolism (odds ratio 2.2; q = 9.3e-18). Inorganic-element-cycling functions included arsenic cycling and detoxification (odds ratio 8.1; q = 1.1e-9) and iron uptake, siderophore, and iron-homeostasis functions (odds ratio 2.0; q = 1.4e-22). Aromatic and xenobiotic-degradation functions included an aromatic and xenobiotic degradation (other) category (odds ratio 13.4; q = 1.0e-32), nitroaromatic/nitrotoluene degradation (odds ratio 4.9; q = 4.1e-7), polycyclic aromatic hydrocarbon degradation (odds ratio 2.7; q = 3.7e-34), and benzoate/aromatic carboxylic-acid degradation (odds ratio 1.8; q = 8.6e-13). Secondary-metabolite functions included terpenoid/carotenoid metabolism (odds ratio 2.6; q = 4.8e-12) and plant-derived/glucosinolate/mycotoxin secondary metabolites (odds ratio 2.1; q = 3.4e-2), and lipid functions included fatty-acid beta-oxidation (odds ratio 2.9; q = 6.7e-27) and lipid degradation (odds ratio 2.1; q = 1.6e-5). Reactive-oxygen-species detoxification was also gut-enriched (odds ratio 4.1; q = 3.1e-51). Together with the transport, cofactor, and sulfur-amino-acid functions above, this set closely tracks the known microbial metabolic functions of the human gut.

The function-level test described in the Methods, which takes the AVG protein as the unit and admits propagated labels alongside reference annotations, was used to ask whether propagation changes the enrichment picture rather than to report enrichment in its own right. Restricted to reference-annotated labels it tested 1,111 functions, all of which were also tested with propagated labels included, while propagation raised a further 29 functions above the 20-protein threshold. Across the 1,111 functions tested both ways, log odds ratios agreed closely (Spearman rho = 0.999), 1,105 (99.5%) kept the same direction of enrichment, and 1,094 (98.5%) received the same enrichment call. The two versions share their reference-annotated proteins, so this is a stability check rather than an independent replication, but it indicates that propagated labels extend the tested function set without redirecting the biome signal.

*Enrichment by ecosystem subtype and predicted host phylum*

Each AMG category was also tested for over-representation within a particular predicted host phylum, relative to the rest of that biome (two-sided Fisher's exact test, BH-adjusted, $q < 0.05$, odds ratio > 1.5; Extended Data Figure 10). In soil, this recovered functions consistent with established host biochemistry, including complex-carbon degradation in Thermoplasmatota (odds ratio 11.5; $q$ = 1.7e-46), manganese cycling and reactive-oxygen-species detoxification in Bacillota (odds ratios 71 and 23; q = 4.1e-63 and 7.8e-45), phosphonate metabolism in Actinomycetota (odds ratio 26; q = 3.9e-91), and polyphosphate metabolism in Pseudomonadota (odds ratio 26; q = 2.5e-25). In the human gut, the strongest host-phylum associations included the Calvin-Benson-Bassham cycle and pentose phosphate pathway in Campylobacterota (odds ratios 15 and 4.7; q = 1.0e-17 and 1.3e-8), cobalamin (B12) metabolism in Patescibacteriota (odds ratio 6.4; q = 8.0e-12), and the phosphotransferase (PTS) system in Bacillota (odds ratio 3.5; q = 2.5e-8).

*Biome- and host-associated AMG functions match known biochemistry*

Auxiliary functions are expected to reflect the metabolism of the virus's host and environment. That the biome- and host-associated AMG functions recovered here match previously reported biochemistry of these hosts and environments provides independent, ecology-based support that the AVGs CheckAMG identifies are biologically coherent rather than annotation artifacts.

#### Generalization of CheckAMG-PST functional propagation to unseen data

*Propagated functional labels reach most sequence-similarity invisible auxiliary genes*

Applying two-tier propagation (Methods) to all 109,107 sequence-similarity invisible AVGs assigned some level of function to 65.2%: 45,392 (41.6%) received a specific function name, 20,651 (18.9%) an L1 function category, and 5,107 (4.7%) only a broad metabolic (14.9% of this group), physiological (15.1%), or regulatory (70.0%) category. The remaining 37,957 (34.8%) received no propagated label at any level (Supplemental Table 10; Extended Data Figure 9D). This depth distribution is consistent with the precision-coverage behavior established on held-out reference proteins below. Propagation reaches a large majority of otherwise-unannotated genes, with depth shallowing as the calibrated evidence for a more specific label runs out.

*Category assignment holds on proteins the model never saw during training*

CheckAMG-PST assigns a broad auxiliary category (metabolic, physiological, or regulatory) to a protein by finding its nearest neighbor among labeled training proteins in embedding space (Methods). We tested whether this works on proteins withheld entirely from training by scoring the true category of each withheld protein against its nearest remaining, non-withheld neighbor (Extended Data Figure 9C). Category assignment was correct 89.5% of the time, far exceeding two controls: randomly shuffling the category labels of the training proteins before propagation, which reduced accuracy to 51.5% (the rate expected by chance), and simply assigning every protein the single most common category, which reached 66.8% (the rate expected without using the model at all). Accuracy was similar whether the nearest matching training protein came from a viral genome (89.0%) or a bacterial genome (92.0%), showing that CheckAMG-PST is not simply relying on the withheld protein sitting in a virus-like genomic neighborhood. Confident category assignments (Methods) were also selective. They were made for 7.24% of likely auxiliary genes but for only 0.0035% of non-auxiliary structural genes used as a negative control (Extended Data Figure 9C).

Because the held-out split is genome-cluster-atomic rather than family-disjoint, most test proteins retain appreciable similarity to a training protein, so precision is reported stratified by sequence identity rather than pooled (Extended Data Figure 9F). Broad-category assignment is stable across the range, from 80.4% in the 0–30% identity band to 92.3% above 90% identity. Specific-function assignment is far more identity-dependent, rising from 34.2% to 58.7% across the same range, which is why specific names are admitted for a much smaller fraction of proteins at a fixed precision target. Sequence identity and embedding distance carry partly independent information: within the highest identity band, precision for specific function names still falls from 0.72 to 0.38 across quintiles of nearest-neighbor embedding distance (Extended Data Figure 9 Source Data). Neither quantity alone is sufficient to decide whether a propagated label is trustworthy, which is what motivates the calibrated model over a simple distance cutoff.

At the 90% realized-precision target used, the deployed two-tier rule assigned a specific function name to 63.5% of held-out test proteins (0.945 realized precision), an L1 function category to a further 26.8% (0.925), and a broad category to a further 1.9% (0.850, over the 806 proteins that reach that depth), leaving 7.8% with no label (Extended Data Figure 9G). Within the embedding tier, ranking candidate labels by the calibrated probability rather than by raw nearest-neighbor distance is what delivers that coverage: at the same target it admits 23.4% of proteins for a specific name, 33.6% for an L1 category and 91.8% for a broad category, against 0.01%, 2.8% and 81.6% for distance alone (Extended Data Figure 9H). Expected calibration error, the average gap between stated and observed accuracy across probability bins, was 0.0077 for specific function names, 0.0055 for L1 categories, and 0.0057 for broad categories, indicating the calibrated probabilities can be interpreted directly (Extended Data Figure 9I).

*Functional propagation ability is largely inherited from the underlying protein language model*

CheckAMG-PST is built by additional training on top of a general-purpose protein language model, ESM-2. To ask how much of the category-assignment ability above comes from that additional training versus from ESM-2 itself, we repeated the same withheld-protein test using the unmodified esm2_t30_150M backbone at matched sample size. The backbone alone assigned the correct broad category 98.8% of the time, exceeding the 89.6% achieved by CheckAMG-PST on the 133,086 proteins matched to the backbone test (Extended Data Figure 9C). Broad auxiliary category is therefore not a property that CheckAMG-PST's additional training confers; it is already largely recoverable from general-purpose protein embeddings, and the genome-context training neither adds to it nor is required for it. Extending the same comparison to the finer levels of the functional hierarchy gave the same ordering rather than the opposite one. At matched sample size the backbone reached 0.925 micro precision for L1 category assignment and 0.823 for specific function names, against 0.587 and 0.382 for CheckAMG-PST (Extended Data Figure 9C; Source Data for Extended Data Figure 9). Stratifying by the maximum sequence identity between each test protein and any training protein shows that the backbone's advantage does not depend on a close training relative being available: the backbone led in every one of the fifteen identity-band-by-depth cells, and at broad-category depth its lead widened as identity fell, from 0.074 in the band above 90% identity to 0.153 for proteins with no recorded identity to any training protein. CheckAMG-PST's contribution therefore lies in the auxiliary-gene call itself rather than in functional label transfer. The fine-tuning objective separates proteins by viral origin and auxiliary-gene likeness, and the set transformer mixes genomic context into each protein's vector, both of which serve the auxiliary-gene call and neither of which preserves the within-class separation that transfer among several thousand specific function names requires. Label propagation would therefore be better served by the untuned backbone, and the function labels reported here are conservative for that reason.

#### Structural matches among the sequence-similarity invisible AVGs

Of the 72,393 sequence-similarity invisible AVGs with a named structural match, 2,586 (3.6%) carry a methyltransferase name, and 2,288 of those fall in the metabolic theme, where they are the largest single class (25.2% of 9,090). DNA methylation is one of five categories CheckAMG flags as warranting additional scrutiny when a gene is called an AMG, alongside essential viral genes and nucleotide, glucan and lipid metabolism. The flag marks a function for interpretation rather than excluding it. 49.0% of CheckAMG's own high-confidence calls carry at least one cautious flag, and the methyltransferase entries in the CheckAMG reference carry a median AMG weight of 0.883, higher than the reference median of 0.813, because they are metabolically specific and less prevalent in viral genomes than the average reference entry. The structural matches show the ambiguity the flag is meant to capture. Of the 2,586, 412 are named as acting on DNA or RNA, including site-specific adenine methyltransferases, type I restriction enzyme methylases and IME4-like RNA methylases, 271 also matched a DefenseFinder model, and 1,665 carry a generic SAM-dependent or unqualified methyltransferase name that a structural match alone cannot resolve. These proteins are therefore reported as auxiliary genes of uncertain metabolic role rather than as evidence of expanded viral metabolism.

#### Supplemental Figures

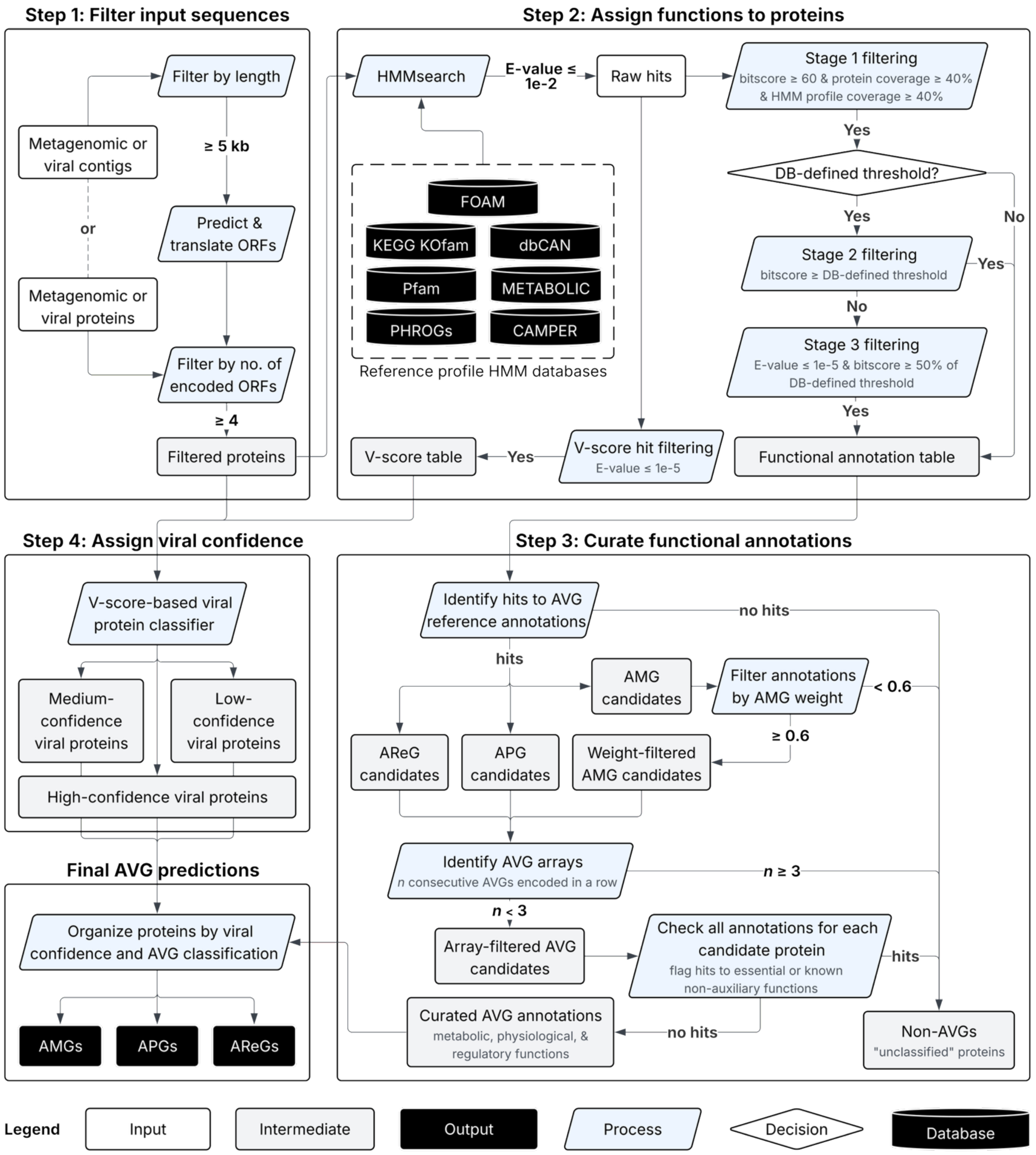


**Supplemental Figure 1. The CheckAMG annotate workflow.** Input contigs or proteins are filtered by length and number of encoded ORFs (Step 1), annotated against seven reference profile HMM databases with staged hit filtering (Step 2), and curated into AMG, APG, and AReG candidates by AMG weight, AVG-array detection, and checks for essential or non-auxiliary functions (Step 3). Each protein is assigned a high, medium, or low viral confidence by the V-score-based classifier (Step 4), and final AVG predictions are organized by viral confidence and AVG type.

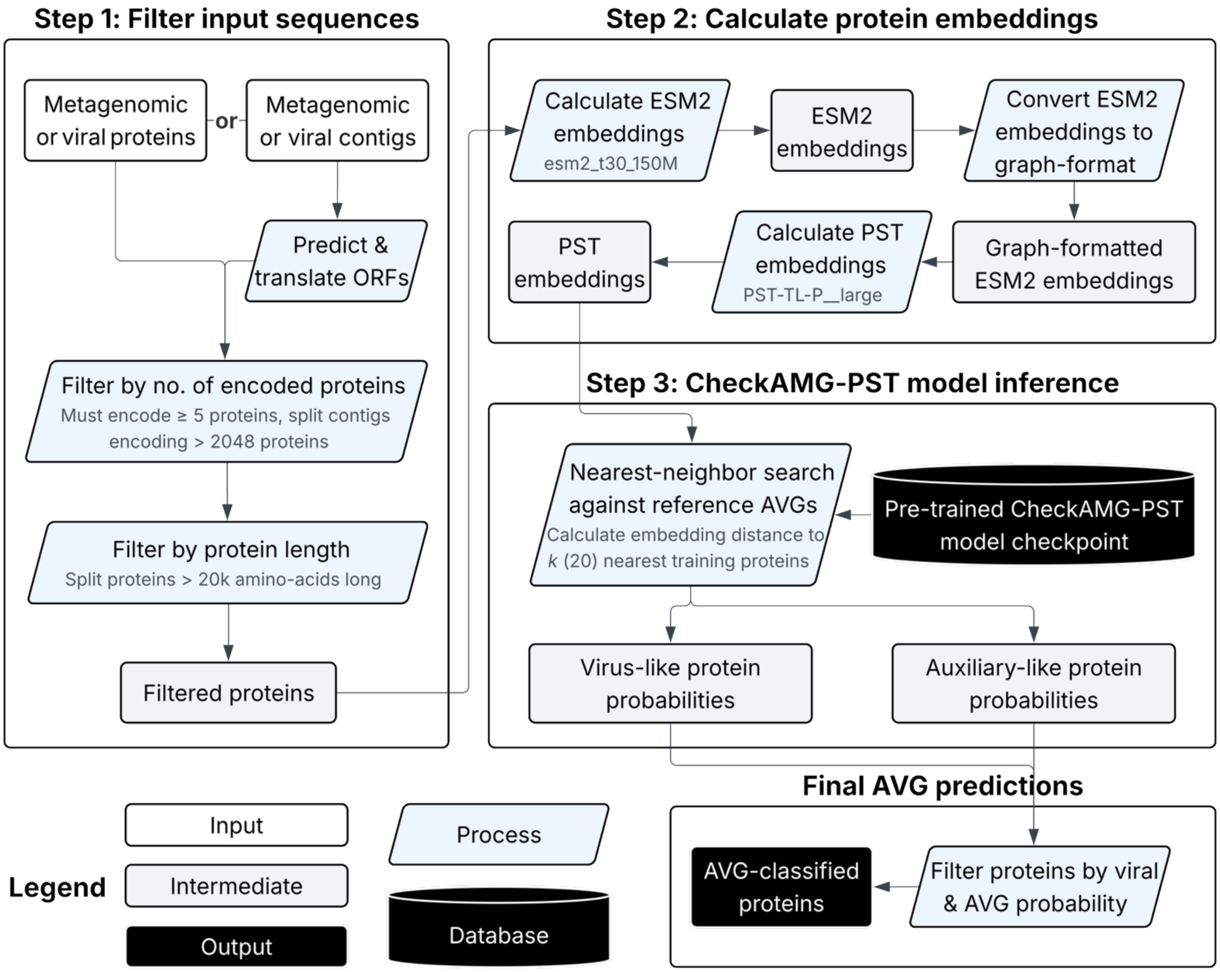


**Supplemental Figure 2. The CheckAMG de-novo workflow.** Input contigs or proteins are filtered by the number of encoded proteins and by protein length (Step 1), embedded with ESM-2 and the Protein Set Transformer (Step 2), and compared to reference AVGs by nearest-neighbor search in CheckAMG-PST embedding space to give per-protein virus-like and auxiliary-like probabilities (Step 3), which are thresholded to produce the final AVG predictions.
